# Deterministic colonization arises early during the transition of soil bacteria to the phyllosphere and is shaped by plant-microbe interactions

**DOI:** 10.1101/2024.10.24.619992

**Authors:** Teresa Mayer, Erik Teutloff, Kerstin Unger, Pamela Lehenberger, Matthew T. Agler

## Abstract

**Background:** Upon seed germination, soil bacteria are activated to transition to the plant and eventually colonize mature tissues like leaves. These bacteria are poised to significantly influence plant health, but we know little about their colonization routes. We studied the mechanisms of the transition of soil bacteria to germinating plants and leaves by experimentally manipulating inoculation times and using *in-planta* isolation to understand bacteria that can make the complex soil-to-leaf transition. Using a trackable, labeled *Pseudomonas viridiflava* (*Pv*3D9) amended to soil, we tested how plant-microbe-microbe interactions shape assembly mechanisms in natural soil communites.

**Results:** We found that the stages of the transition of bacteria from soil to leaves before true leaf emergence were important in establishing uniquely diverse leaf bacteriomes. A wide diversity of leaf-associated taxa can individually make this transition, but most are still shaped by stochastic processes. Interestingly, deterministic processes drove some important taxa only when they transitioned from soil to leaves, but not when they were inoculated later. The opportunistic pathogen *Pv*3D9 promoted plant growth in a natural soil, likely by activating plant immunity. These effects in turn strongly affected the soil-to-leaf transition of almost strictly taxa that colonized deterministically, demonstrating the important role of plant-microbe-microbe interactions in controlling deterministic processes.

**Conclusions:** Diverse, well-adapted bacterial taxa make the soil-to-leaf transition during natural colonization resulting in characteristic diversity in healthy leaf microbiomes. The domination of stochastic mechanisms during this colonization indicates that many taxa must strongly compete to establish their niche. During this complex transition, however, specific important taxa emerge that are driven by deterministic processes, suggesting they occupy unique niches. The malleability of these processes suggests that they may be controlled, for example by targeted soil manipulations. This finding is significant given the important roles of these bacteria in plant health and offers directions forward for engineering beneficial plant microbiomes.

## Background

Plant leaves are colonized by diverse microbes, including bacteria, fungi and oomycetes. This microbiome plays a critical role in determining how plants deal with stresses like drought, disease and herbivory [1, 2]. In particular, the bacterial component of the microbiome aids plant survival in the presence of detrimental soil microbes [3, 4] and can protect leaves against pathogen attack [5]. Soil is thought to be an important source of inoculum for all plant tissues, but only a small fraction of soil microbes will later make up a significant proportion of the plant microbiome [6]. This filtering is not surprising when the many barriers to a successful transition of bacteria to germinating seedlings and ultimately mature leaves are considered. Upon germination, a radicle and then cotyledons emerge into the vast microbial diversity contained in soil [7]. Growth of true leaves follows only later. Thus, for soil bacteria to reach aerial leaf tissue, they must deal with edaphic (soil) factors [8], competitively reach host tissue and establish a niche [9, 10], and either grow together with the plant to establish in or on emerging leaves [11] or invade the tissues later. Although these early stages are likely very important determinants of the complex transition of bacteria from soil to leaves and ultimately leaf microbiome structure and plant health, how they play out and the processes governing them are not well understood.

Stochasticity plays major roles in microbial community assembly in diverse environments [12, 13]. In the context of the plant, stochastic colonization essentially indicates that a niche could be occupied by many different bacteria with similar fitness, making them mostly exchangeable. Given the massively diverse soil inoculum, stochasticity could dominate early immigration processes from a complex inoculum like soil to germinating seedlings. On the other hand, bacteriome structure in mature leaves appears to be determined at least partly by deterministic factors like host selection [14], and is strongly influenced by the interaction between host factors and the environment [6]. Indeed, while the abundance of many leaf taxa are correlated to their abundance in soil, suggesting stochastic dynamics, there is extensive variation on this trend that can in part be explained by host factors [15]. Importantly, recently developed tools can use microbiome structure data to quantify the contribution of stochastic or deterministic processes both on the level of whole microbiomes and on the level of individual taxa, which can reveal taxa that may occupy specialized niches in hosts [16].

The importance of stochastic and deterministic processes in natural colonization may also be shaped by direct or indirect interactions between bacteria [17]. In plant leaves, early arriving pathogens can alter the immune state of the host and thereby facilitate the colonization of late comers [18–20]. Alternatively, early colonizers could occupy essential nutrients like iron [21] or other favorable niches [22]. Experiments simulating invasion of mature leaves have shown that priority effects can strongly influence bacteriome assembly, with early colonizers persisting and influencing subsequent colonization outcomes [23]. During natural colonization from soil, most bacterial taxa making the transition to leaves are likely to start from very low cell numbers, given the massive soil bacterial diversity and relatively low abundance in soil of leaf-colonizing bacteria [15]. Whether or not the arrival of low- abundance soil bacteria can significantly affect colonization processes is unknown. However, this would be important because it may suggest ways to shape colonization mechanisms and community assembly trajectories.

Here, we experimentally dissected the transition of leaf-colonizing bacteria from soil to leaves to better understand how it shapes leaf bacteriomes and ultimately plant health. We used three different *Arabidopsis thaliana* genotypes, including two wild genotypes isolated from Jena, Germany, and compared the assembly mechanisms of bacterial communities in mature leaves when plants germinated in inoculated soil vs when inoculation occurs post- germination. Next, we tested whether and how diverse bacterial taxa can efficiently colonize leaves starting from extremely low levels near a germinating seedling, as a simple soil diversity model suggests they must do in soil. Finally, we amended natural soil with a trackable opportunistic pathogenic bacterium to better understand how community assembly mechanisms are shaped by bacterial interactions. The results reveal new insights into the types of niches occupied by bacteria that transition from soil to leaves and suggests that these niches, and thus their recruitment, is at least partly manageable.

## Materials and Methods

### Plant materials

We utilized three *A. thaliana* wild-type genotypes: Col-0, NG2 and PB. Col-0 is a widely available model genotype of *A. thaliana*, while NG2 and PB are genotypes that were previously recovered from wild populations in Jena, Germany [14]. Near-isogenic lines of each were generated via multiple generations started from single seeds. NG2 is available from NASC (N2110865). PB is being made available there as well and is available upon request from the authors.

### Inoculation time experiment

#### Preparation of microboxes

In total 18 microboxes each containing 9 flow pots [24] were prepared, six for each of the three *A. thaliana* genotypes (Col-0, NG2 and PB). After they were autoclaved, opening the microboxes for inoculation and sampling was performed in a laminar flow hood. At the first day of the experiment (ID0) the Flow Pots were flushed with 25mL sterile MilliQ water to wet the substrate. Seeds had been surface-sterilized with 2% bleach and 70% ethanol and were stratified in the dark for 3 days in 0.1% agarose at 4°C. Between 7 and 10 seeds were added to each pot, and these were later thinned to 1-3 plants per pot. More detailed box preparation instructions can be found the supplementary methods.

#### Inoculations

For each plant genotype, two microboxes were randomly assigned to be inoculated with a natural soil microbial community on the first day of the experiment (ID0), two microboxes at day 7 of the experiment (ID7), one microbox at day 14 of the experiment (ID14), and one microbox was never inoculated with a natural living soil microbial community (Never - control). Inoculation was performed by injecting a natural soil slurry + Murashige and Skoog (MS) medium mixture into the soil via a tube attachment (5mL of inoculum) and by spraying from above with an airbrush system (three sprays for half a second each). To ensure that the only difference between treatments was the time of inoculation with live microbiota, at each inoculation point all plants that did not receive a live inoculum received a double-autoclaved inoculum. The boxes were closed and sealed with medical tape (Duchefa Biochemie). Detailed information on the preparing the natural soil slurry and seed preparation can be found in the supplementary methods.

#### Plant growth, sampling and processing

The prepared boxes with seeds were incubated in growth chambers (PolyKlima PK-520) at 20°C /14°C 16h/8h day/night cycle with 75% light intensity. Humidity was not controlled, but given the environment inside the closed boxes, was likely close to saturation. Plants were sampled on day 14 (SD14), day 21 (SD21), day 28 (SD28) and day 35 (SD35). At each sampling day except SD35, rosettes were collected with a flamed tweezer and placed in a screw cap tube with sterile beads (two 3mm steel beads (WMF) and 0.2g of 0.25-0.5mm glass beads (Carl Roth)) and stored at -80°C until further processing (1 whole plant rosette = 1 sample). At SD35 only three leaves per plant were combined into one sample due to the large size of plants: an old, a middle-aged and a young leaf. DNA was extracted by bead-beating, followed by incubation in a CTAB buffer, direct precipitation in ethanol/sodium acetate then ethanol washing. The DNA was further purified using a magnetic bead-based cleanup. DNA extraction details can be found in the supplementary methods.

#### *In-planta* isolation of leaf bacteria experiment

##### Production of inoculum from leaves

We sampled leaves of wild plants in spring 2018 and 2019. In 2018, we collected 10 *A. thaliana* rosettes of different sizes from the wild population NG2, located in Jena, Germany. They were washed three times with autoclaved ultrapure water to remove dirt, ground with a metal pestle, and suspended in PBS/S (PBS buffer + 0.02% Silwet L-77) by vortexing. After centrifugation (1 min, 19,000 x g), the inoculum was aliquoted, mixed with glycerol (final glycerol concentration: 21.5%) and stored at -80°C. In spring 2019, we sampled *A. thaliana* leaves from populations NG2 and PB and processed them in the same manner. For the experiments, the stocks were diluted in PBS/S so that the inocula finally contained 0-1 cells/seedling (NG2, 2018) or an average of 3.5 and 1.45 cells/seedling (NG2 and PB 2019, respectively).

#### Plant growth, inoculation and identification of successful leaf colonizers

*S*eeds were surface sterilized with 2% bleach and 70% ethanol, washed three times with autoclaved ultrapure water and vernalized for four days in 0.1% agarose at 4 °C in the dark. We sowed one seed in each well of a 24-well plate filled with 1 mL ½ MS + 0.2% sucrose + 1% agar. The plants were incubated in growth chambers (PolyKlima PK-520) at 24°C/18°C 16h/8h day/night cycle with 50% light intensity. 5 µL of leaf microbe extract was inoculated. In 2018, NG2 inoculum was also added to liquid and solid R2A medium in 96-well plates and petri dishes, respectively, and onto the cotyledons of 2-day and 6-day old NG2 seedlings. In 2019, NG2 or PB inocula were added to 3-day old NG2, PB or Col-0 seedlings. In both years we mock inoculated control seedlings with PBS/S, and did not observe contamination. Media inoculations were incubated at 26 °C for 15 days. Inoculated plants were grown for two weeks after which whole rosettes were harvested with flamed tweezers. After removing roots and any flower stems, they were transferred into 1.5 mL tubes (2018) or to deep 96-well plates (2019) and frozen at -80°C until further processing. In the 2018 experiments in NG2 plants, we identified colonized plants by extracting DNA from leaf material and PCR amplifying the V5-V7 region of the 16S rRNA gene, where the bacterial band is clearly separate from the plant band (see detailed supplementary methods). Reactions with a bacterial band were sent for sanger sequencing (Eurofins Genomics). In the 2019 experiments in NG2, PB and Col-0 plants, we crushed and plated dilution series of the rosettes to look for bacterial colonies. They were identified by DNA extraction and PCR amplification of the entire 16S rRNA gene region, followed by Sanger sequencing (see detailed supplementary methods).

#### *P. viridiflava* 3D9 soil amendment experiment

##### Soil preparation and plant growth

Surface-sterilized *A. thaliana* NG2 seeds (2% bleach and 70% ethanol) were stratified in the dark for 3 days in 0.1% agarose at 4°C. On the day of the experiment, a base soil mix was prepared by mixing 4L Floraton 3 potting soil (Floragard), 2L perlite (0-6mm Perlite Perligran, Knob), and 25g Substral Osmocote garden flower fertilizer (Celaflor). This was wetted with either 2L of 10 mM MgCl_2_ (soil “LS”) or with 2L of MgCl_2_ mixed with 200g of dried garden soil (soil “LS+GS”). For the *P. viridiflava* 3D9- 141 treatments, 3D9-141 was added to the MgCl_2_ or MgCl_2_ + soil slurry to target 1,000 CFU/cm^3^ or 1,000,000 CFU/cm^3^ of soil. If the spermosphere (zone of influence) of a germinating seed (0.5mm x 0.25mm x 0.25mm) reaches a distance into the surrounding soil equaling its own dimensions, then this works out to about 0.8 (“low”) and 800 (“moderate”) 3D9-141 CFUs in the spermosphere, respectively. Vernalized seeds were sown with 15 to 20 seeds per pot (5.5 cm upper diameter) and incubated at 193 μmol/(m^2^·s) at 22 °C/18 °C and 16h/8h day/night conditions. To minimize batch and positional effects, the tray positions were switched randomly every two days.

##### Pv*3D9-141 preparation*

*Pv*3D9 was fluorescently labeled by integrating pMRE-Tn7-141 plasmid containing the mTurquoise2 gene as well as chloramphenicol and gentamicin resistance cassettes [25] to generate *Pv*3D9-141. It was grown in LB medium supplemented with gentamycin and chloramphenicol at 220 rpm and 28 °C to OD_600_ of roughly 1.5, the cells were then harvested by centrifugation for 5 min at 2000 rcf, the supernatant removed, and the culture washed once with 10 mM MgCl_2_. After centrifugation, the cultures were resuspended in 10mM MgCl_2_ before diluting into the MgCl_2_ wetting solution.

##### Pv3D9-141 effects on shoot growth

Seven days after germination, seedling shoots were sampled by removing entire plants from the soil and cutting off the root with flamed scissors and tweezers. Because plants were very small, each replicate is the average of 8 plants from one pot that were measured together. At the same time, the number of plants in one pot was thinned out to three.

##### Estimating bacterial and Pv3D9-141 loads in grown plants with CFU counting

To count bacterial loads in the leaves, 14 days after germination, six samples of two to three leaves each were collected and stored at –80 °C until they were processed. For total bacteria and 3D9-141 CFU counting, frozen leaf samples were crushed in 120 µL of 10 mM MgCl_2_ using 3 mm diameter sterilized metal beads (WMF) in a mini bead-beater 96 (Biospec Products) at 1400 strokes/min. After homogenization, a dilution series in 10 mM MgCl_2_ were drop-spotted onto three different media in duplicates and the plates were incubated at 28 °C and counts were made after 72 h (details on the media and their selection can be found in the supplementary methods). Additionally, a fluorescence stereo microscope equipped with a GFP filter was used to record the number of colonies fluorescing in that general emission range, which includes mTurquoise2.

### qPCR quantification of 3D9-141 leaf loads

For qPCR quantification of 3D9-141 and 16S rRNA gene amplicon sequencing, 14 days after germination, six samples of two to three leaves each were collected and stored at –80 °C until gDNA was extracted. DNA was extracted by bead-beating in a CTAB buffer, followed by a phenol-chloroform cleanup, precipitation in isopropanol and ethanol washing. Details on DNA extraction can be found in the supplementary methods. For the qPCR, the single-copy *mTurquoise2* gene integrated in the 3D9-141 genome was targeted. For normalization of copy numbers to the host, we targeted the *A. thaliana EF1-*α gene [26]. Copy numbers were estimated with standard curves generated from a 1:8 dilution series of the linearized pMRE-Tn7-141 plasmid and a 1:8 dilution series of NG2 gDNA. Details on the primers, reaction components, controls and thermocycler setup are provided in the supplementary methods.

### Bacterial community characterization via 16S rRNA gene amplicon sequencing

#### Library preparation

The extracted gDNA was diluted 1:10 in 10 mM Tris-HCl (pH = 8) and used as a template for 16S rRNA gene amplification. The amplicon sequencing library preparation targeted the V3-V4 region of the 16S rRNA gene and used blocking oligonucleotides to avoid plant plastid amplification, as previously described [27] with a few modifications (see detailed steps in supplementary methods). The final library was spiked with 10% PhiX genomic DNA to ensure high enough sequence diversity and was loaded onto an Illumina MiSeq and sequenced for 2x300 cycles. Amplicon sequencing data was split on indices and adapter sequences were trimmed using Cutadapt 1.2.1 [28]. The quality of the reverse reads was not satisfactory, so we proceeded with high quality forward reads only, which were clustered into amplicon sequencing variants (ASVs) using dada2 [29]. First, reads were trimmed and filtered to remove reads with more than 3 errors per 100bp. Sequences were then dereplicated and denoised using the error rate information before calling amplicon sequence variants (ASVs). Chimeric ASVs were removed and taxonomy was assigned using the latest Silva 16S rRNA gene database (v. 138.1) [30].

### Microbial diversity analysis

Downstream analysis was performed in R with phyloseq [31] and other packages listed below. Host-derived reads were removed by filtering any ASVs in the orders “Chloroplast” and “Ricketsialles” from the 16S ASV tables and data was filtered to remove samples with less than 500 reads. For further processing the reads were either not grouped (ASV level) or grouped at the genus and order levels. Beta diversity was based on the Aitchison distance and the “RDA” function was used to perform either unconstrained principle components analysis on the Aitchison distance matrix or a redundancy analysis on the matrix constrained by experimental factors, as stated in the text and figures. Statistical significance of factors was tested with an ADONIS test (a permutational analysis of variance) with 999 permutations. Beta dispersion (between-sample beta diversity) was also based on Aitchison distance with statistics as described in the figures. The alpha diversity metrics Chao1 (estimate total richness of community) and Shannon (index accounting for both richness and evenness) were used and statistical analysis was based on a non-parametric Kruskal-Wallis test. In the inoculation time experimental results, to identify taxa whose abundance in mature leaves significantly varied with experimental variables, we used DESeq2 [32] on data from combined sampling days 21, 28 and 35. The log-transformed abundances of taxa identified by DESeq were plotted in boxplots for visual inspection with a p-value based on a non-parametric Kruskal-Wallis test. In the *Pv*3D9-141 amendment experiment, we identified the taxa that were most correlated to the RDA axes constrained by treatment and made boxplots of those.

For the analysis of processes driving colonization, we employed the tool “Infer Community Assembly Mechanisms by Phylogenetic bin-based null model analysis” (iCAMP 1.5.12) [16]. First, we generated a phylogenetic tree. The 16S rRNA sequences of the ASVs were aligned using the DECIPHER (2.20.0) function AlignSeqs and highly variable regions were masked using MaskAlignment. The masked alignment was then converted to the phangorn package (2.11.1) with the function phyDat and output as a fasta alignment with the function write.phyDat [33]. Finally, the alignment was fed to RAxML Next Generation (1.2.1) to generate a phylogenetic tree using the GTR+G+I model with default parameters (20 tree searches with 10 random and 10 parsimony-based starting trees) [34]. iCAMP was run by loading the OTU table, taxonomy information and tree and generating a treatment table that linked the samples to the plant ecotype and inoculation day. For the environment table, the ecotype information was provided. To optimize parameters, first the phylogenetic niche preference was assessed and then the within-bin phylogenetic signal is measured. We tested a range of values for ds and bin.size.limit and chose those where both the relative abundance of bins with significant phylogenetic signal (RAsig.adj) and the mean correlation coefficient across bins (meanR) showed a peak. For ds = 0.2, and bin.size.limit=20, in addition, we made sure that the relative contribution of stochastic processes was comparable to the phylogenetic normalized stochasticity ratio, as suggested by the authors. iCAMP was then run with these parameters using “Confidence” as the significance index (a non-parametric, one-sided confidence score) and the beta mean pairwise distance (bMPD) as the metric for phylogenetic null model analysis. 1000 randomizations were used for confidence checking. Finally, bin-level statistics were produced using bootstrapping and were output to a text file.

## Results

### Early colonization of germinating seedlings shapes mature leaf bacteriomes

To study how early colonization shapes bacterial communities in *Arabidopsis thaliana* leaves, we disrupted natural colonization by controlling when plants of three genotypes were inoculated by a natural soil microbiota (Day 0, D7, D14 or Never). To ensure that treatment effects are mainly due to live microorganisms and not, for example, immune priming by molecular patterns, all plants that did not receive live inoculum at each treatment time received a heat-treated inoculum. The three inoculation times are all early in development, corresponding to the seed (D0), cotyledon (D7) and first true leaf (D14) stages of plant growth (**Figure 1**).

**Figure 1.**
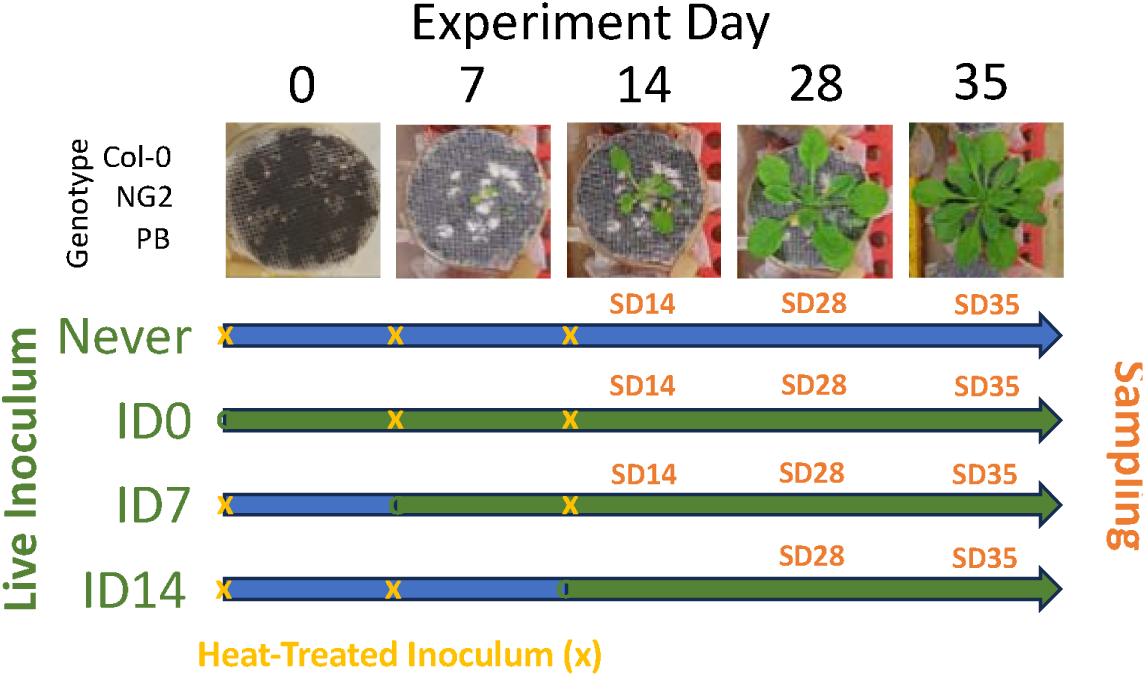
Diagram of flow pot experiment 1, showing for each of three treatments and the control the time of incoulation with live inoculum (start of green arrows), heat-killed inoculum (yellow x) and the sampling times (SD).

Bacterial community structure was more strongly shaped by inoculation day than either sampling day (SD14, SD21, SD28 or SD35) or plant ecotype (Col-0, NG2 or PB) at all investigated taxonomic levels (ASV, Genus and Order). In particular, the inoculation day effect was strongest in the Col-0 genotype (explaining up to 18% of variation at the Order level) but was also significant in NG2 and PB (explaining about 10% of variation in each) (**Figure 2A, S1A and S2A, Tables 1, S1 and S2**). In all genotypes, mature leaf bacteriomes (21 days post inoculation) that had been inoculated at D0 were more similar to those inoculated at D7 than to those inoculated at D14, especially at higher taxonomic levels (**Figure 2B, S1B and S2B and between-sample beta diversity in Fig. S3**). This is because many taxa were significantly enriched in early (D0 or D7)-inoculated plants compared to D14-inoculated plants (**Figures S4-S6**), leading to general patterns of higher alpha diversity and usually higher evenness in D0- and D7-inoculated plants (**Figure 2C, S1C and S2C**).

**Figure 2.**
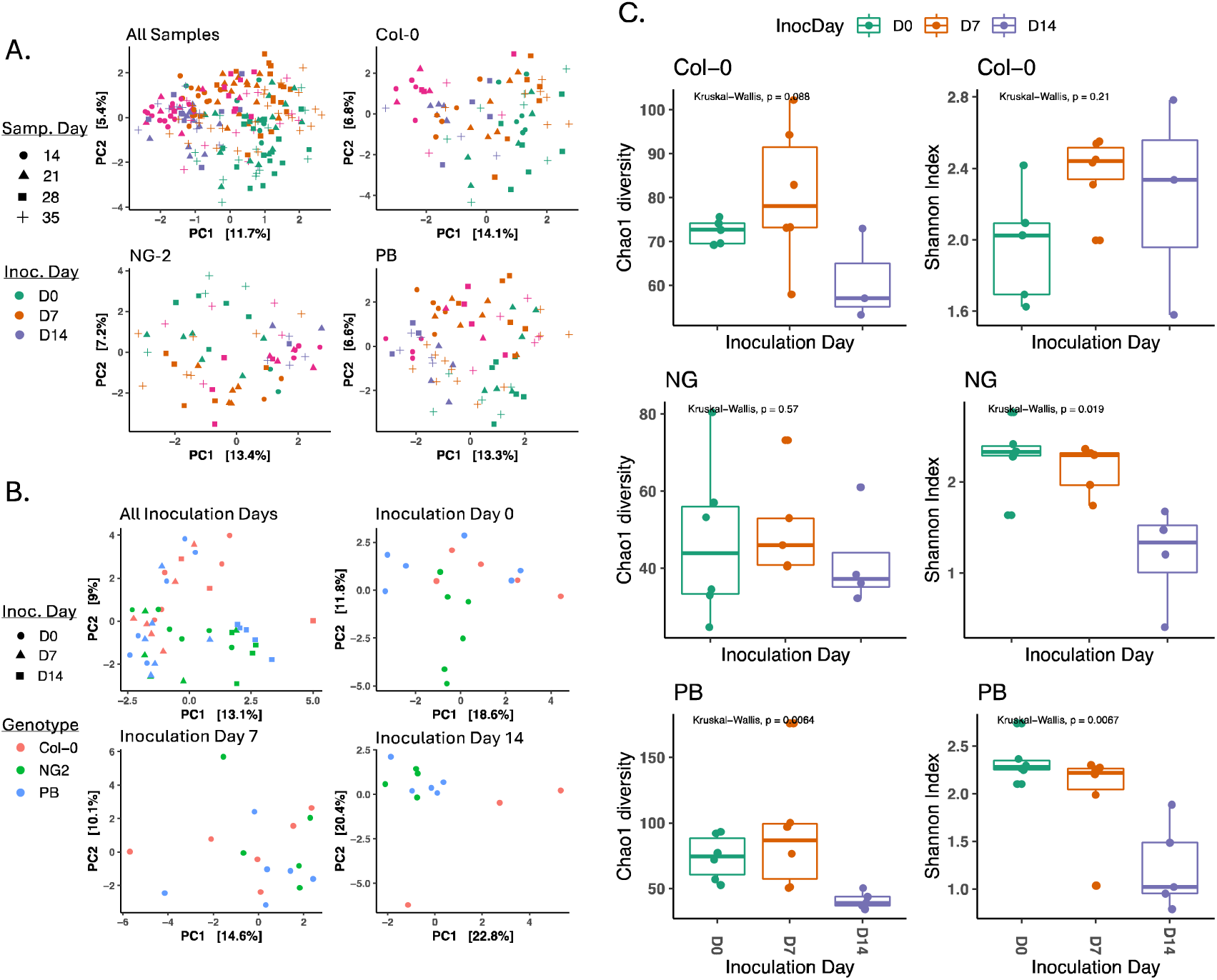
Early-inoculated plants (before the emergence of true leaves) develop distinct and diverse microbiomes in mature leaves. Data is based on genus-level taxonomy of bacteria associated with the mature leaves of *A. thaliana* after inoculation at day 0, 7 or 14 (D0, D7, D14). **A.** RDA ordination of beta diversity (aitchison distance) of all samples including multiple sampling times. Results of a PERMANOVA test for significance of factors is in Table 1. **B.** RDA ordination of beta diversity (aitchison distance), including only samples collected 21 days post inoculation. Results of a PERMANOVA test for significance of factors is in Table 2. **C.** Alpha diversity (Chao1 estimates of total diversity and Shannon Index) of samples collected 21 days post inoculation.

**Table 1:**
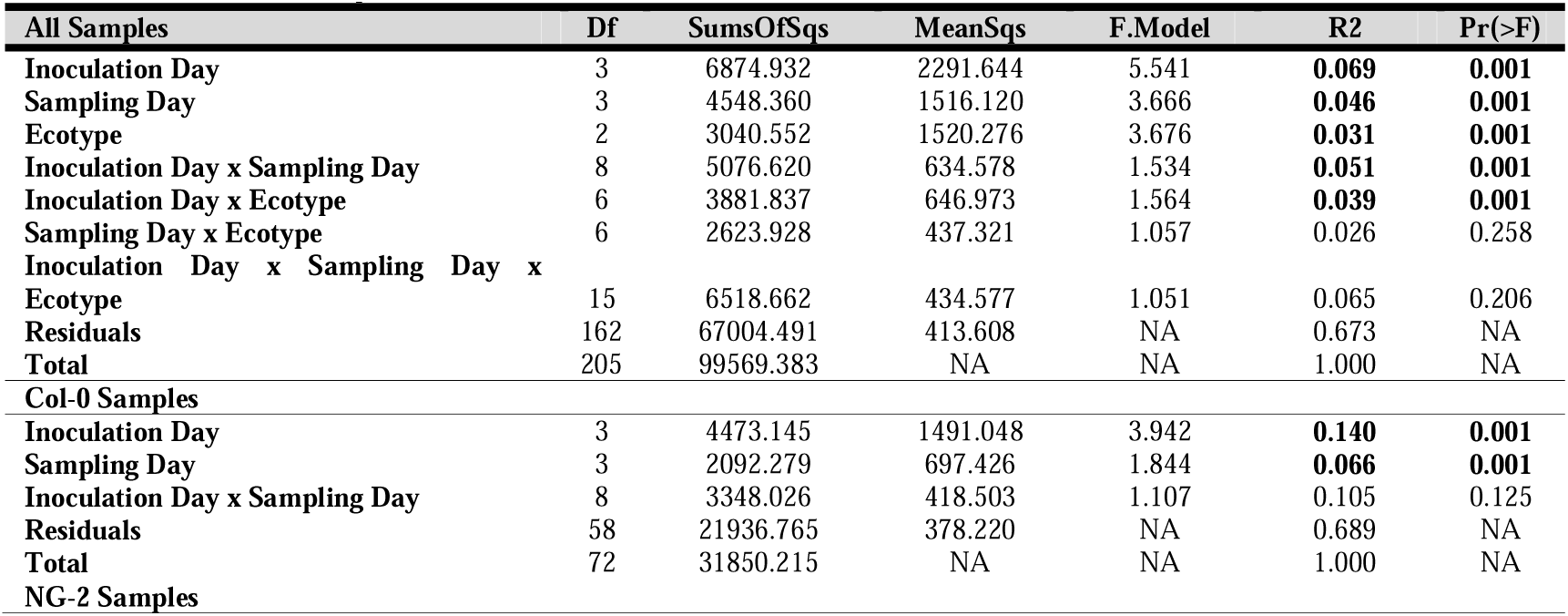
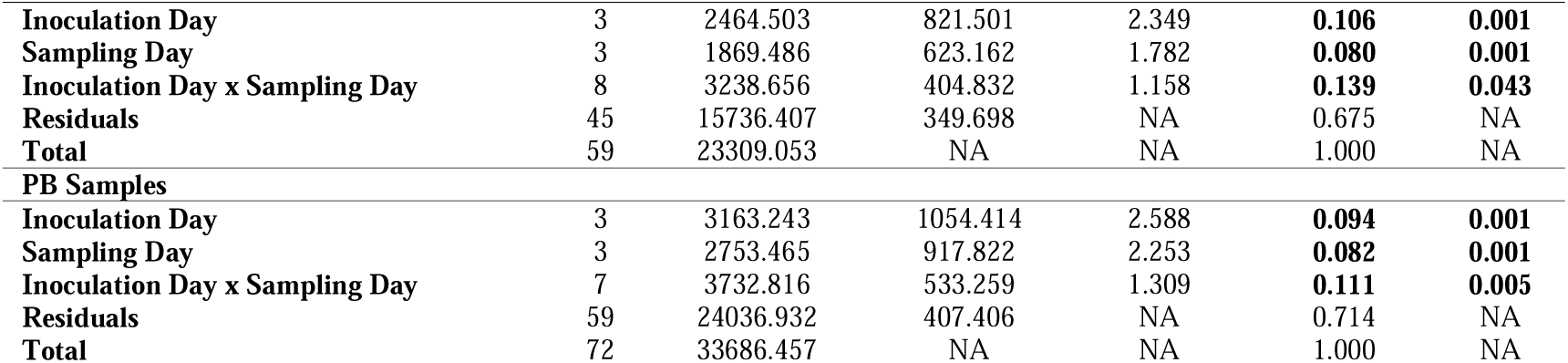
Significance of factors in describing the variation in bacterial communities between samples. Results are based on the Aitchison distance between samples collected at all time points at the Genus level. Analysis was carried out with PERMANOVA with 1000 permutations.

**Table 2:** Significance of factors in describing the variation in bacterial communities between samples. Results are based on the Aitchison distance between samples collected at 21 DPI at the Genus level. Analysis was carried out with PERMANOVA with 1000 permutations.

| All | Df | SumsOfSqs | MeanSqs | F.Model | R2 | Pr(>F) |
| --- | --- | --- | --- | --- | --- | --- |
| Inoculation Day | 2 | 2743.151 | 1371.576 | 3.258 | **0.126** | **0.001** |
| Ecotype | 2 | 1547.307 | 773.654 | 1.838 | **0.071** | **0.002** |
| Inoculation Day x Ecotype | 4 | 1963.144 | 490.786 | 1.166 | 0.090 | 0.078 |
| Residuals | 37 | 15574.781 | 420.940 | NA | 0.714 | NA |
| Total | 45 | 21828.384 | NA | NA | 1.000 | NA |
| **Inoculation Day 0** |  |  |  |  |  |  |
| Ecotype | 2 | 841.835 | 420.918 | 1.456 | **0.172** | **0.022** |
| Residuals | 14 | 4048.468 | 289.176 | NA | 0.828 | NA |
| Total | 16 | 4890.303 | NA | NA | 1.000 | NA |
| **Inoculation Day 7** |  |  |  |  |  |  |
| Ecotype | 2 | 1074.012 | 537.006 | 1.245 | **0.151** | **0.047** |
| Residuals | 14 | 6040.560 | 431.469 | NA | 0.849 | NA |
| Total | 16 | 7114.572 | NA | NA | 1.000 | NA |
| **Inoculation Day 14** |  |  |  |  |  |  |
| Ecotype | 2 | 617.728 | 308.864 | 1.667 | **0.270** | **0.003** |
| Residuals | 9 | 1667.487 | 185.276 | NA | 0.730 | NA |
| Total | 11 | 2285.216 | NA | NA | 1.000 | NA |

The effect of plant genotypes also depended on inoculation time. Specifically, plants inoculated at D0 or D7 had more taxa that were differentially enriched between the genotypes (43 and 57 ASVs, respectively) than plants inoculated at D14 (28 ASVs **Figure S7**). Additionally, 15 were shared between D0- and D7-inoculated plants, while only 3 were shared between D0- and D14-inoculated plants. However, plant genotype explained more beta diversity in D14-inoculated plants (p<0.01, with 26-28% of variation depending on taxonomic level) than in plants inoculated at D0 or D7 (11-17%, insignificant at some taxonomic levels) (**Tables 2, S3 and S4**). This can be explained because although many more taxa were differentially enriched in D0-inoculated plants, some differentially enriched taxa in D14-inoculated plants were highly abundant, especially in the Col-0 genotype (**Figures S8-S9**). Overall, these results show that leaf bacteriome assembly is shaped uniquely by the early transition of bacteria from soil to plant leaves, resulting in higher diversity and more taxa that are influenced by plant genotypes.

### In the overall stochastic transition of bacteria to leaves, some taxa transition deterministically

To understand why plants that germinated in inoculated soil (inoculation D0) were different than those inoculated later (inoculation D14), we studied the mechanisms of assembly of bacteria in leaf microbiomes (stochastic vs. deterministic processes). To do so, we employed a phylogenetic bin-based null model analysis [16], which determines to what extent different colonization processes shape phylogenetically grouped bins of ASVs. Overall, the model showed that in all genotypes and inoculation times, stochastic processes dominated (>60% contribution to assembly, **Figure 3A**), but colonization was slightly more stochastic in D0- than in D14-inoculated plants. Despite the dominance of stochasticity, deterministic processes were still important and were dominated by homogenous selection (HoS), making up ∼15-35% relative importance depending on plant genotype (highest in Col-0, lowest in PB) and inoculation time (**Figure S10A and S10B**). These results were validated based on agreement with a second, independently calculated stochasticity metric, the phylogeny-based normalized stochasticity ratio [35] (**Figure S10D**).

**Figure 3.**
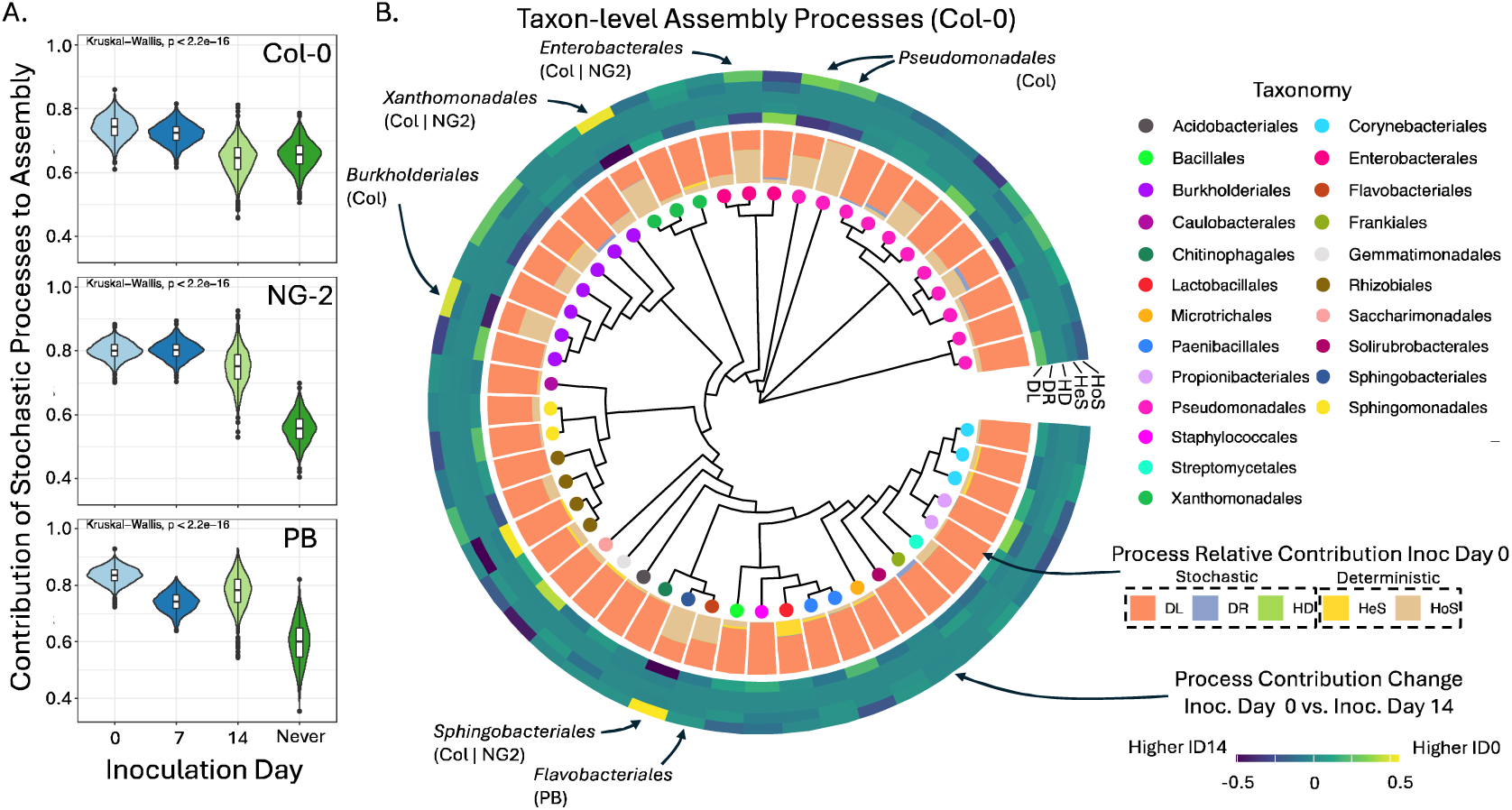
Processes shaping bacterial colonization of leaves of the *A. thaliana* genotypes Col-0, NG2 and PB depend on both inoculation time and genotype. **A.** The relative importance of stochastic processes in colonization of the leaves of plants inoculated at ID0, ID7, ID14 or never inoculated. Stochasticity represents the sum of dispersal limitation, homogenous dispersal and drift, but was consistently dominated by dispersal limitation. Deterministic processes (homologous and heterologous selection), as well as an independend measure of stochasticity (pNST) are shown in Figure S10. **B.** Assembly processes at the level of taxonomic bins, shown for Col-0, trees for other genotypes are shown in Figure S11. The tree is annotated with, from inside to outside: Colored dots representing the majority taxonomy of each bin, A barchart showing for plants inoculated at ID0 the relative contribution of each community assembly process for each bin (DL: Dispersal limitation, DR: Drift, HD: Homogenizing dispersal, HeS: Heterogeneous selection, HoS: Homogeneous selection), and A heatmap showing the change in relative importance of each assembly process for each bin between plants inoculated at ID0 and those inoculated at ID14 (yellow means a given process is more important at ID0, dark blue is more important at ID 14). The labeled taxa are those mentioned in the text that were selected deterministically at ID0 and stochastically at ID14 (homologous selection increased by at least 20% to reach >50% total contribution in D0). The plant genotypes where this difference was observed are labeled.

At the level of phylogenetic bins, colonization processes were usually dominated by dispersal limitation (DL), a stochastic process, and this was consistent across genotypes and inoculation times (**Figure 3B)**. Some taxa, however, were more strongly influenced by deterministic processes. Pseudomonadales were especially strongly and consistently shaped by deterministic HoS, in particular bin Pseudomonadales 41 (including only *Pseudomonas* ASVs) (**Figure 3B, Figure S11** and detailed bin data in **Supplementary File 1**). This determinism can help explain why many outlier ASVs that were unusually abundant for their frequency (fraction of samples where they were observed) were *Pseudomonas* (**Figure S12**). Several other bins also colonized deterministically, but particularly in D0-inoculated plants (>50% contribution from HoS – barchart in Figure 3B). These were Sphingobacterales 19 and Enterobacterales 38 (in Col-0 and NG2, mostly *Pedobacter* and *Enterobacter* ASVs, respectively), Xanthomonadales 35 and Burkholderiales 17 (in Col-0, mostly *Stenotrophomonas* and *Janthinobacterium*, respectively), Pseudomonadales 40 and 41 (in Col-0 and PB, mostly *Pseudomonas*), and Flavobacteriales 20 (in PB, mostly *Chryseobacterium*). Notably, all of these (except Pseudomonadales 41 and 49 in NG2 and Col-0) were only deterministic in D0-inoculated plants but colonized stochastically in D14- inoculated plants (difference between D0 and D14 was >25%) (heatmap in **Figure 3B, Figure S11 and Supplementary File 1**). For several of these same bins, HoS was more important at D0 than D14 in other genotypes as well but did not reach >50% at D0 (Burkholderiales 17 and Flavobacteriales 20 in NG2 and Enterobacterales 38 in PB). Together, these results show that after seedling germination in soil, natural colonization of mature leaves is a stochastic process for many colonizers. Most taxa that do colonize deterministically, however, do so only during the complex transition from soil to leaves, helping to explain why plants colonized naturally from soil assemble distinct leaf bacteriomes.

### Many leaf bacteria have the capacity to transition from soil to leaves

The previous results showed that following plant germination in soil, bacteria mobilize to manage the transition to mature plant leaves via stochastic or deterministic processes. Using a simple diversity model, we estimated cell number distributions for strains in theoretical homogenous soils (see details in **Supplementary Results**). This analysis suggests that most potential leaf-colonizing bacterial strains probably start the colonization process from just a few cells in the spermosphere (the area under influence of a germinating seedling). Since little is known about whether and how different bacteria can manage this task, we developed a technique to inoculate vanishingly low numbers of cells onto germinating, axenic seedlings and to identify bacteria that manage the transition to leaves. In a first experiment, a natural leaf microbiome was extracted from leaves collected from plants in the wild NG2 population. This extract was extinction-diluted to guarantee 0-1 cells per inoculation (two dilutions with ∼0.5 or ∼0.05 cells/inoculation **Table S5**) and inoculated onto R2 liquid or solid media and onto germinating, axenic NG2 seedlings (**Figure 4A**). Mature leaves, individual colonies on solid media or liquid media were harvested two weeks later and PCR/Sanger sequencing was used to identify “isolates”. The frequencies of recovery of individual taxa in media vs. leaves were compared. Colonizers in positive plants were mostly *Pseudomonas* sp. and *Xanthomonas* sp., but two were also positive for a *Rhizobiaceae* and one for a Burkholderiales. The number of plants with *Pseudomonas* sp.*, Xanthomonas* sp. or *Rhizobiaceae* was equal to or more than the expected amount based on the number of colonies of those taxa recovered on media (**Figure 4B, Supplementary Results and Table S6**). Thus, nearly every culturable cell of these taxa that was inoculated onto germinating seedlings reached detectable levels in leaves two weeks later.

**Figure 4.**
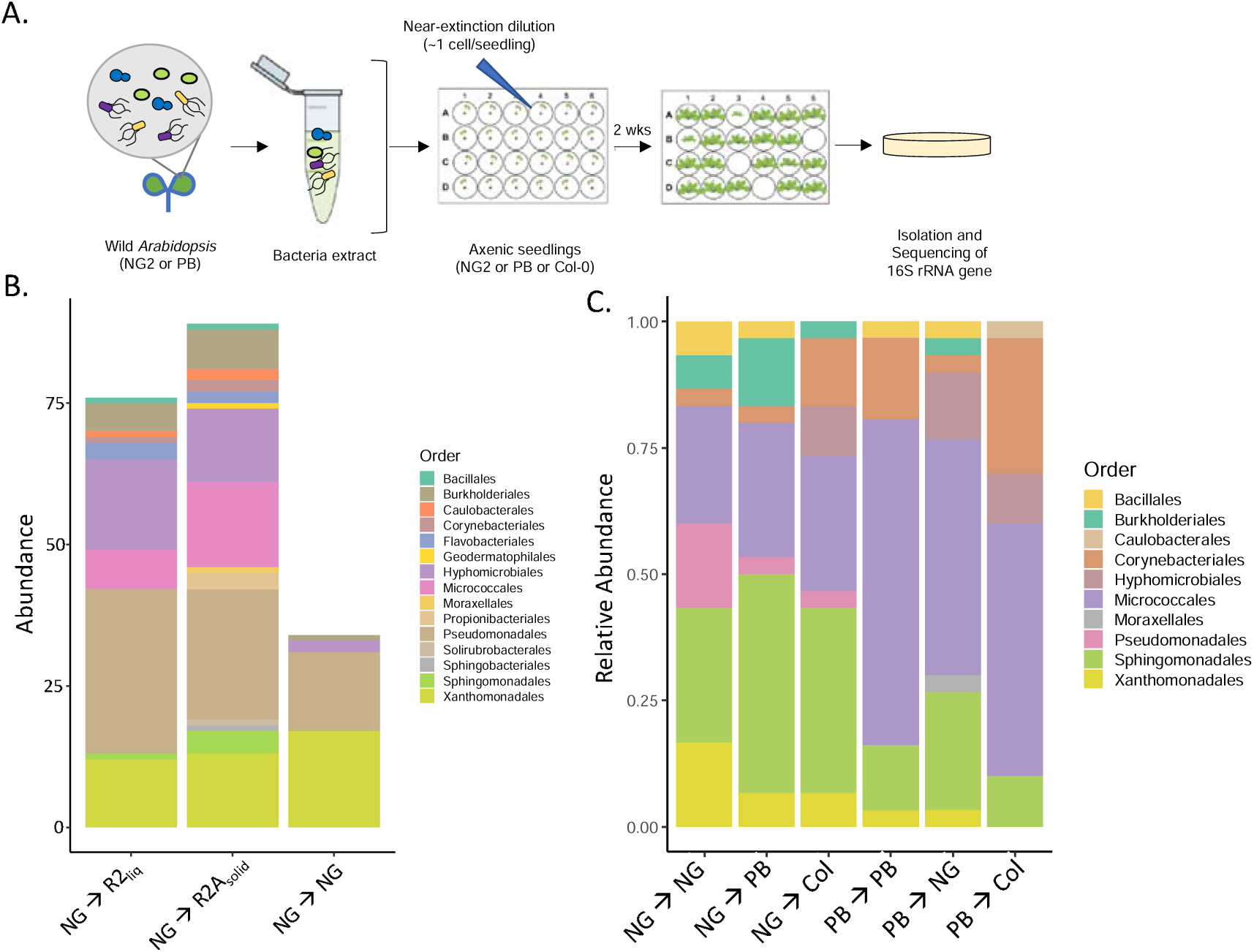
Many leaf bacteria can reach and colonize leaves when only a few cells encounter a germinating seed. (A.) The in-planta isolation method based on dilution-to-extinction of leaf-associated microbes from wild *A. thaliana* leaves and inoculation onto germinating axenic seedlings, followed by two weeks of growth and isolation and identification of colonizers. This was used to identify leaf bacteria that are likely to be efficient colonizers when highly dilute in soil. (B.) Bacteria collected from wild NG2 plants were diluted to near extinction (0-1 cells/plant) and inoculated onto liquid or solid R2 medium or onto germinating seedlings. The bar chart shows the taxonomy of randomly picked isolates from the medium and from mature leaves where colonization was detected by PCR. (C.) Bacteria collected from wild NG2 or PB plants were diluted to 3.5 and 1.45 cells/seedling and inoculated onto germinating NG2, PB or Col-0 plants. Bar chart showing taxonomy of 30 random colonizers from each treatment (31 for PB à PB). Taxonomy is grouped at the order level, taxonomy at the genus level and associated statistics can be found in **Figure S14** and **Table S7.** Detailed data on numbers of plants inoculated and colonized are found in **Table S5**.

In a second, larger experiment, inocula from wild NG2 and PB populations were diluted to about 3.5 and 1.45 cells/seed, respectively. Each was then inoculated onto 120 axenic 3-day old NG2, PB and Col-0 seedlings. Two weeks later, leaf material was collected, crushed, and plated on R2A media. We observed bacteria in 25.0% (NG2 -> PB) to 55.8% (PB -> NG2) (**Table S5**) of inoculated plants. From 160 plants, 181 bacteria were isolated and identified. These were diverse, including 27 genera from three gram-positive (Bacillales, Corynebacterales and Micrococcales) and seven gram-negative families (Burkholderiales, Caulobacterales, Hyphomicrobiales, Sphingomonadales, Moraxellales, Pseudomonadales and Xanthomonadales) **(Figure 4C)**. 140 plants (87.5%) had only one discernable colony. These genera included taxa that colonized both deterministically and stochastically from soil in the previous experiment. *Pseudomonas*, *Janthinobacterium* and *Sphingomonas* isolates were significantly more often enriched from NG2 leaf extracts, while *Clavibacter* was more abundant from PB leaf extracts (**Figure S14, Table S7**). There was generally little discernible effect of the recipient plant genotype, with only *Curtobacterium* from PB slightly more frequently detected in PB plants than in other genotypes (□^2^: p = 0.082**, Table S7**). In 20 plants (12.5%), more than one bacterium was co-isolated. For *Clavibacter, Methylobacterium* and members of the Burkholderiales (*Janthinobacterium*, *Variovorax*, *Acidovorax* and *Rugamonas*), this was significantly more often than expected (□^2^: p<0.05), suggesting they likely benefit from a partner (**Table S8**). Especially the 7 total Burkholderiales isolates (all Comamonadaceae and Oxalobacteriaceae) always were together with a co-colonizer, although this would be expected only once by chance alone (□^2^: p = 2.56x10^-12^). Together, these findings confirm that even stochastically colonizing leaf-associated taxa can efficiently transition to leaves from minute levels near a germinating seedling. Additionally, even from these extremely low inoculation levels, interactions between taxa seem to shape the mechanisms driving colonization.

### Plant-microbe interactions shape colonization of deterministic taxa

Determinic colonization indicates that the bacteria is not easily replaced by another, a sign that they could occupy a specialized niche. Thus, we reasoned that treatments that strongly affect plant phenotypes would most strongly affect colonization of these taxa. To test this, we selected *Pseudomonas viridiflava* 3D9 from the *in-planta* isolation experiment (**Figure 4C, treatment NG**→**NG**). This strain is an opportunistic pathogen native to the NG2 genotype – it was isolated from healthy plant leaves collected in the wild just like other less virulent bacteria, but just a few cells inoculated onto axenic NG2 seedlings is enough to nearly always kill the host plant, whereas less virulent bacteria colonize without symptoms (**Figure S15**). Opportunistic *P. viridiflava* is known to strongly affect plants by interacting with their immune system, among other mechanisms [36]. Thus, we reasoned that it should be adapted to efficiently transition from soil to plants in natural colonization and be poised to exert effects on other colonizers. A labeled version, 3D9-141, was generated by integrating a vector with markers into the genome [25] so that it can be tracked in a complex microbial background via antibiotic selection, fluorescence and qPCR.

First, we evaluated the effects of 3D9-141 colonization in two soils. Plants were grown in a standard laboratory potting soil (LS) or the same soil combined with a microbial extract of natural garden soil (LS+GS). The soils were amended with 3D9-141 at three levels, none (“mock” treatment), ∼0.8 cells in the initial spermosphere (“low”), and ∼800 cells in the initial spermosphere (“moderate”). 3D9-141 reached mature leaves in the moderate treatment as measured by both qPCR (**Figure 5A**) and resistant, fluorescent CFUs (**Figure S16**), but not in the low treatment. Based on total CFU counts, 3D9-141 made up roughly 1% of the leaf bacteriome and its median level in leaves was not significantly affected by the addition of GS. In the mock (no 3D9-141) treatments, LS+GS decreased the average fresh weight of seedlings by roughly 35% compared to LS, possibly due to the addition of detrimental soil microorganisms. The 3D9-141 amendment reversed this, increasing the leaf fresh weight in the low treatment back to LS levels. In the moderate treatment, shoot fresh weight was less than the low treatment, but still higher than mock levels **(Figure 5B)**. Thus, the opportunistic pathogen 3D9-141 can transition to plants and leaves depending on its inoculation level and in natural soil it always had a significant, positive effect on plant phenotypes.

**Figure 5.**
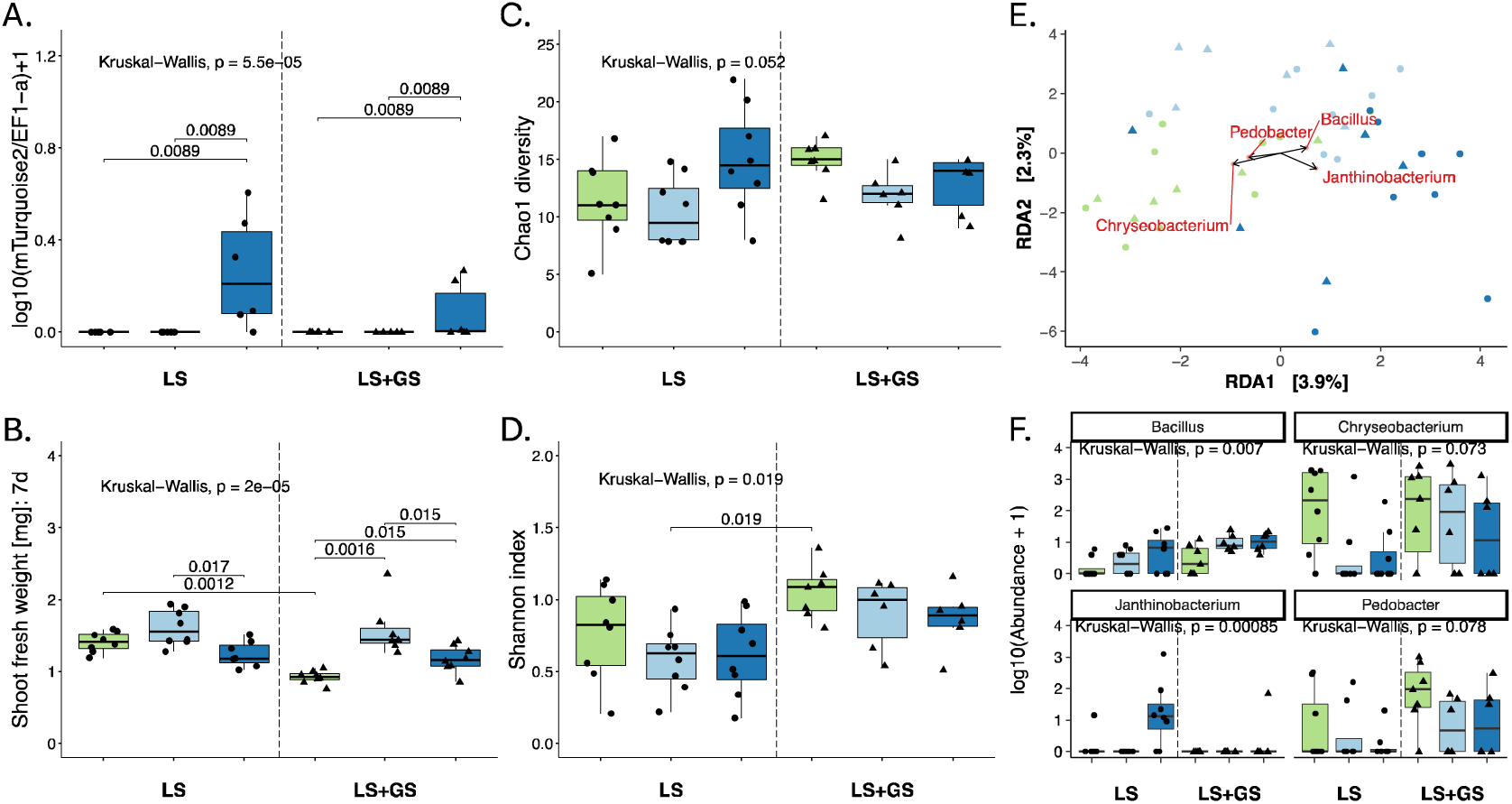
Bacteria that deterministically transition from soil to mature leaves are influenced by other colonizers. Throughout, green corresponds to the mock treatment (no 3D9-141), light blue to the low 3D9-141 soil amendment, and dark blue to the moderate 3D9-141 soil amendment. (A.) *Pseudomonas viridiflava* 3D9-141 abundance in mature leaf tissue based on qPCR measurement of copies of a vector-inserted gene for the mTurquoise2 protein normalized to the host plant elongation factor 1. (B.) The fresh weight of *A. thaliana* shoots 7-days post-germination. (C. and D.) Alpha diversity of bacterial genera in mature leaf tissues (14 days post germination) assessed by Chao1 metric (total estimated diversity) and Shannon index (diversity considering evenness of distirubution of taxa). In A.-D., a Kruskal-Wallis test was performed on the data overall and when it was <0.05, a Wilcoxon test was used to evaluate differences between groups. Signficant p-values are shown. (E.) Principal components analysis of Aitchison distance between samples, constrained for treatment, showing biplot arrows for the top 3% of bacteria that correlate to the two axes. Circles correspond to LS and triangles to LS+GS, as in the boxplots. (F.) Boxplots showing the abundance of the bacteria identified using the biplot analysis in (E) with p-value result of a Kruskal-Wallis test for significant differences between groups.

Next, we tested how 3D9-141 influenced colonization of other bacteria. Based on CFU counts, there was no clear effect of the soil treatment or 3D9-141 on total leaf bacterial loads (**Figure S16**). The addition of garden soil did not significantly increase the bacterial diversity in mature leaves, but it did increase the evenness of the community (**Figure 5C and 5D**). 3D9-141 appeared to slightly affect the medians of both alpha diversity and evenness, but it was not significant with conservative non-parametric statistics. It did, however, have a significant effect on bacterial community structure, with the effect of the soil type and 3D9- 141 treatments each independently explaining about 6-10% of variation depending on the taxonomic level (**Table 3**). Generally, low and moderate 3D9-141 treatments clustered separately, driven by distinct changes to several taxa (**Figure 5E**). Two of the four genera most strongly affected by 3D9-141 were affected similarly in both soils: *Bacillus* (Bacillales), whose abundance always increased with 3D9-141 treatment and *Pedobacter* (Sphingobacterales), which tended to be suppressed by 3D9-141. On the other hand, *Chryseobacterium* (Flavobacterales) was strongly suppressed by 3D9-141 at both low and moderate levels mainly in LS, while *Janthinobacterium* (Burkholderiales) was much higher abundance in LS only in the moderate treatment, when 3D9-141 reached leaves, but was hardly detected at all in LS+GS (**Figure 5F**). Similar patterns at the ASV and order level underscore that effects were highly taxa-specific and depended on both soil and 3D9-141 inoculation level (**Figure S17 and S18**). Notably, three of four genera affected by 3D9-141 (*Chryseobacterium*, *Janthinobacterium* and *Pedobacter*) were also strongly shaped by homogeneous selection in natural colonization of NG2 plants (**Figure 3**). Thus, taxa that behave deterministically in natural colonization appear to also be those that are most responsive to the effects of soil amendment with 3D9-141. The precise taxa that were influenced were different in the low vs. the moderate levels, where 3D9-141 reached leaves.

**Table 3:** Significance of factors in describing the variation in bacterial communities between samples. Results are based on the Aitchison distance between samples collected at 14 DPI. Analysis was carried out with PERMANOVA with 1000 permutations on data agglomerated at the ASV, genus or order levels.

| Factor (PERMANOVA ASV level) | Df | SumsOfSqs | MeanSqs | F.Model | R2 | Pr(>F) |
| --- | --- | --- | --- | --- | --- | --- |
| 3D9-141 Treatment (Mock, low, moderate) | 2 | 1507.641 | 753.820 | 1.593 | **0.068** | **0.004** |
| Soil type (LS/LS+GS) | 1 | 2138.834 | 2138.834 | 4.519 | **0.097** | **0.001** |
| Treatment x Soil | 2 | 971.754 | 485.877 | 1.027 | 0.044 | 0.394 |
| Residuals | 37 | 17511.574 | 473.286 | NA | 0.791 | NA |
| Total | 42 | 22129.803 | NA | NA | 1 | NA |

| Factor (PERMANOVA Genus level) | Df | SumsOfSqs | MeanSqs | F.Model | R2 | Pr(>F) |
| --- | --- | --- | --- | --- | --- | --- |
| 3D9-141 Treatment (Mock, low, moderate) | 2 | 372.982 | 186.491 | 1.363 | **0.062** | **0.075** |
| Soil type (LS/LS+GS) | 1 | 368.900 | 368.900 | 2.695 | **0.061** | **0.002** |
| Treatment x Soil | 2 | 245.005 | 122.503 | 0.895 | 0.040 | 0.648 |
| Residuals | 37 | 5064.163 | 136.869 | NA | 0.837 | NA |
| Total | 42 | 6051.050 | NA | NA | 1 | NA |
| Factor (PERMANOVA Order level) | Df | SumsOfSqs | MeanSqs | F.Model | R2 | Pr(>F) |
| 3D9-141 Treatment (Mock, low, moderate) | 2 | 265.792 | 132.896 | 1.771 | **0.078** | **0.008** |
| Soil type (LS/LS+GS) | 1 | 224.703 | 224.703 | 2.995 | **0.066** | **0.001** |
| Treatment x Soil | 2 | 128.486 | 64.243 | 0.856 | 0.038 | 0.683 |
| Residuals | 37 | 2775.851 | 75.023 | NA | 0.818 | NA |
| Total | 42 | 3394.831 | NA | NA | 1 | NA |

## Discussion

### The transition of bacteria from soil to phyllosphere is an important colonization route

Bacteriomes that develop in mature leaves after seedling germination have important implications for host plant health [37]. The goal of this work was to gain better insight into how the well-known effects of host and environmental factors [6] shape natural colonization processes in these important communities. Leaf bacteriomes are composed of cells that must arrive in one of two ways: Transitioning following seed germination, either from soil or from cells attached to or in seeds that were transmitted vertically from parent plants, or later arrival and “invasion” of already emerged leaves. Transitioning from soil is thought to be a very important route, since leaf bacteria are typically mostly composed of a subset of the taxa found in soils [6]. The importance of the “soil route” is also recognized in commercial applications, where seed impregnation with biological treatments is meant to improve colonization success of beneficial microorganisms [38]. However, this route is complex: Bacteria must find and reach germinating seedlings and ultimately multiply on or in the growing plant to eventually establish in true leaves that only emerge later. Additionally, our simple model of soil bacterial diversity illustrated the complication that there will usually just be a few chances for success, since most strains will start from just a handful of cells in the vicinity of the seedling. To our knowledge there is little known about which bacteria efficiently manage this complex task in a natural context, how they do so and what influences them. However, we here experimentally showed that many bacteria can and do manage this transition and that it leaves distinct signatures in the leaf microbiome.

### Natural colonization results in uniquely diverse leaf microbiomes

In the transition to leaves from a diverse natural soil, we found that most bacterial colonizers exhibited stochastic assembly patterns. In neutral theory, stochasticity indicates that the differences in fitness between taxa is not significant, making their distribution largely subject to neutral processes including dispersal limitation, homogenizing dispersal, and drift [16]. Thus, while many bacteria can reach and colonize leaves, most apparently do so with similar fitness. This can enable simultaneous transitioning from soil of many taxa that are essentially exchangeable and can help explain the unique and high diversity of taxa in naturally colonized plant leaves (inoculation day 0 in this study). It should be noted that stochasticity in this context refers only to niche fitness *differences* among leaf-colonizing bacteria and does not imply that leaf colonization is just a random selection of environmental bacteria. On the contrary, leaf bacterial diversity is usually far less than soil bacterial diversity, indicating that “phyllosphere-associated bacteria” are different than other soil bacteria that do not make the transition [39]. Supporting this, in our 1-on-1 plant-bacteria enrichment experiment, a wide diversity of typical leaf bacteria could start from just a few cells near a germinating seedling and weeks later reach detectable levels in mature leaves. Thus, these taxa are apparently well-adapted to thrive on primary resources from host plants and to deal with the harsh conditions in the leaf environment, even if they do so stochastically. Host physiology, and thus primary resources that stochastic taxa depend on, will be variable across host species and environments [40]. Thus, an approach like what we have used here can help identify these efficient colonizers. This would be important for applications, to know which taxa will thrive when given an advantage, for example via seed impregnation or soil amendments.

At least two factors could explain stochastic recruitment of many or most leaf bacteria. For one, many leaf colonizers could use similar resources and have similar fitness so that none drastically outcompete others. Resource overlap, however, was previously found to be a relatively poor predictor of colonizer success on leaves [41]. On the other hand, resource overlap is usually calculated from resource preferences from metabolic models or *in-vitro* experiments because of the difficulty of measuring activity of bacteria *in-planta*. Indeed, we still know very little about niches during host colonization [42] and it is possible that there is extensive overlap among commensal bacteria. A second possibility is that space is limited and occupied on a “first-come-first-served” basis. This is supported by various studies showing that bacteria colonizing healthy leaves are limited spatially, for example to epidermal cracks [43] or microscopic droplets [44] and that colonizers that establish there tend to be stable to subsequent invasion [23]. Regardless, the process of transitioning from soil to mature leaves is complex and may involve bacteria transitioning between multiple niches before reaching leaves. Thus, to understand and manipulate it, improved techniques to understand niches and niche diversity of non-pathogenic bacteria *in-planta* will be required.

### The emergence of determinism in the midst of stochasticity

Our results suggest that most leaf-colonizing bacteria transitioned from soil to leaves stochastically. However, a few taxa showed strong signs of determinism (specifically homogeneous selection). It is not surprising that several *Pseudomonas* bins consistently colonized deterministically, given the well-described adaptation of *Pseudomonas* to *A. thaliana* colonization [45]. On the other hand, bins dominated by *Pedobacter, Enterobacter, Stenotrophomonas, Janthinobacterium*, and *Chryseobacterium* also colonized deterministically. In contrast to stochastic selection, which occurs when multiple taxa can occupy a niche with similar fitness (i.e., are exchangeable), homogenous selection implies that the taxa likely have more specialized niches [16]. Supporting this, we previously found that several taxa, including Burkholderiales taxa in the families Comamonadaceae and Oxalobacteraceae (like *Janthinobacterium*), are strongly enriched in some *A. thaliana* genotypes due to aliphatic glucosinolates, specialized metabolites of *A. thaliana*. We were able to identify a specialized metabolic niche whereby only certain bacteria with myrosinase enzymes (including an Enterobacterales) can metabolize glucosinolates as a carbon source, and specific cross-feeding interactions support enrichment of other taxa (*Pseudomonas* and perhaps *Janthinobacterium*) [14]. A similar phenomenon was recently observed in maize, where specialized lactonases enable specific bacteria to grow on benzoxazinoids, leading to rhizosphere enrichment [46]. Given the wide diversity of specialized metabolites that plants produce [47], we speculate that these can be the basis of many specialized niches that underly deterministic recruitment in plants. However, new approaches are needed to understand what is available to colonizers and how bacteria take advantage of them. The approach we use here to identify deterministic taxa may be a good way to identify those whose colonization niche should be studied more closely.

### Deterministic processes in natural colonization rely on the soil-to-leaf transition

Notably, here we found that except for *Pseudomonas,* all the taxa that were deterministic in the transition from soil to leaves colonized stochastically when soil inoculum was applied directly onto leaves (inoculation day 14). It is currently unclear why deterministic niche establishment in leaves would be dependent on at least some part of the complex transition from soil. However, several explanations are plausible. First, germinating seedlings exude a broad mix of compounds into the soil matrix that shift in composition over time [48]. Exudates can serve as signals that cause soil bacteria to alter their phenotypes to promote host colonization [49]. Therefore, exudates secreted early on could serve as a signal to activate the metabolic pathways required to establish a specialized niche. Second, plants use molecular recognition machinery to react specifically to different types of microbes [50], where responses include altered secondary metabolism. Thus, the makeup of the soil microbiome at early colonization timepoints, by altering plant physiology and chemistry, can lead to altered patterns of deterministic colonization. This is consistent with our finding that even very low-level natural soil amendment with *Pv*3D9, an opportunistic pathogen, promoted plant growth, likely indicating that it altered the plant immune status. These changes to plant physiology had either direct effects or effects that rippled into the leaf microbiome, but these were almost strictly limited to deterministically colonizing taxa, suggesting that processes they depend on had been altered.

A third possibility could be that important interactions between deterministic taxa develop during early colonization. Our results strongly suggest that deterministic colonization of leaves by some taxa arises out of inter-bacterial interactions. In the *in-planta* isolation experiments, *Oxalobacteraceae* and *Comamonadaceae* bacteria were always only found with a partner bacterium. Consistently, our previous work showed that *Oxalobacteraceae* and *Comamonadaceae* colonization in *A. thaliana* leaves depended on aliphatic glucosinolates, but *in-vitro* tested isolates probably relied on another bacterium that could metabolize the compound [14]. Similarly, in an enrichment of bacteria from peppers on the secondary metabolite capsaicin, two partners (a *Pseudomonas* and the Comamonadaceae *Variovorax*) fully relied on one another to use it as the sole carbon and nitrogen source [51]. More broadly, Comamonadaceae and Oxalobacteraceae have been highlighted as being strongly positively correlated to a range of bacteria in wild *A. thaliana*, including links to both Sphingobacteriaceae and Flavobacteriacae (which include *Pedobacter* and *Chryseobacter*ium, taxa found here to colonize deterministically with *Janthinobacterium*) [52, 53]. Thus, we hypothesize that these bacteria enter into diverse interactions with other bacteria that define their niche and promote deterministic colonization. Regardless, once taxa are identified that colonize deterministically in a given environment, an important next step will be elucidating the molecular underpinning of the colonization mechanisms to realize targeted manipulations.

## Conclusions

Understanding how plant-associated bacteria assemble in tissues like leaves is a critical step toward harnessing their potential for sustainable plant protection. While soil clearly is an important inoculum of bacteria for leaf tissues, the complex transition of soil bacteria to leaves following seedling germination has largely remained a black box. This work demonstrated that early stages following germination are critical for establishment of diverse bacteriomes in mature leaves. This is because of the unique mix of stochastic and deterministic mechanisms that arises during early colonization to shape individual bacterial taxa. Deterministic mechanisms especially are shaped by both host genotype and host-microbe interactions. Thus, characterizing the mechanisms shaping bacterial taxa offers a way forward for targeted manipulation and optimization of leaf-associated bacteriomes.

## Declarations

### Ethics approval and consent to participate

Not applicable

### Consent for publication

Not applicable

### Availability of Data and Materials

The 16S rRNA sequencing datasets generated and analysed during the current study are available in the NCBI SRA repository BioProject number PRJNA1174406, https://www.ncbi.nlm.nih.gov/bioproject/PRJNA1174406.

Scripts used to analyze the data in R to create the figures and tables, as well as the raw data tables and metadata files will all be made publicly available before publication on Figshare: https://doi.org/10.6084/m9.figshare.27290064.v1.

### Authors’ Contributions

MTA and TM conceived and designed the study. TM performed experiments on inoculation times, PL and KU performed experiments for in-planta enrichment, ET performed the experiments with *Pv*3D9. TM, PL, KU, ET and MTA contributed to data analysis and generating figures. TM, KU, ET and MTA contributed to writing the text. All authors helped edit and approved the final text.

## Acknowledgements

We acknowledge René Maskos, Stefan Riedel, and Kirsten Küsel (Aquatic Geomicrobiology, Friedrich-Schiller-University Jena) for sequencing our libraries on their MiSeq instrument.

