## Supplementary Information for "Deterministic colonization arises early during the transition of soil bacteria to the phyllosphere and is shaped by plant-microbe interactions"

Supplemenary Information for:

**The supplementary information consists of:**

**Figures S1 – S18**

**Tables S1 – S8**

**Supplementary Methods**

**Supplementary Results**

**Figure S1.**

| 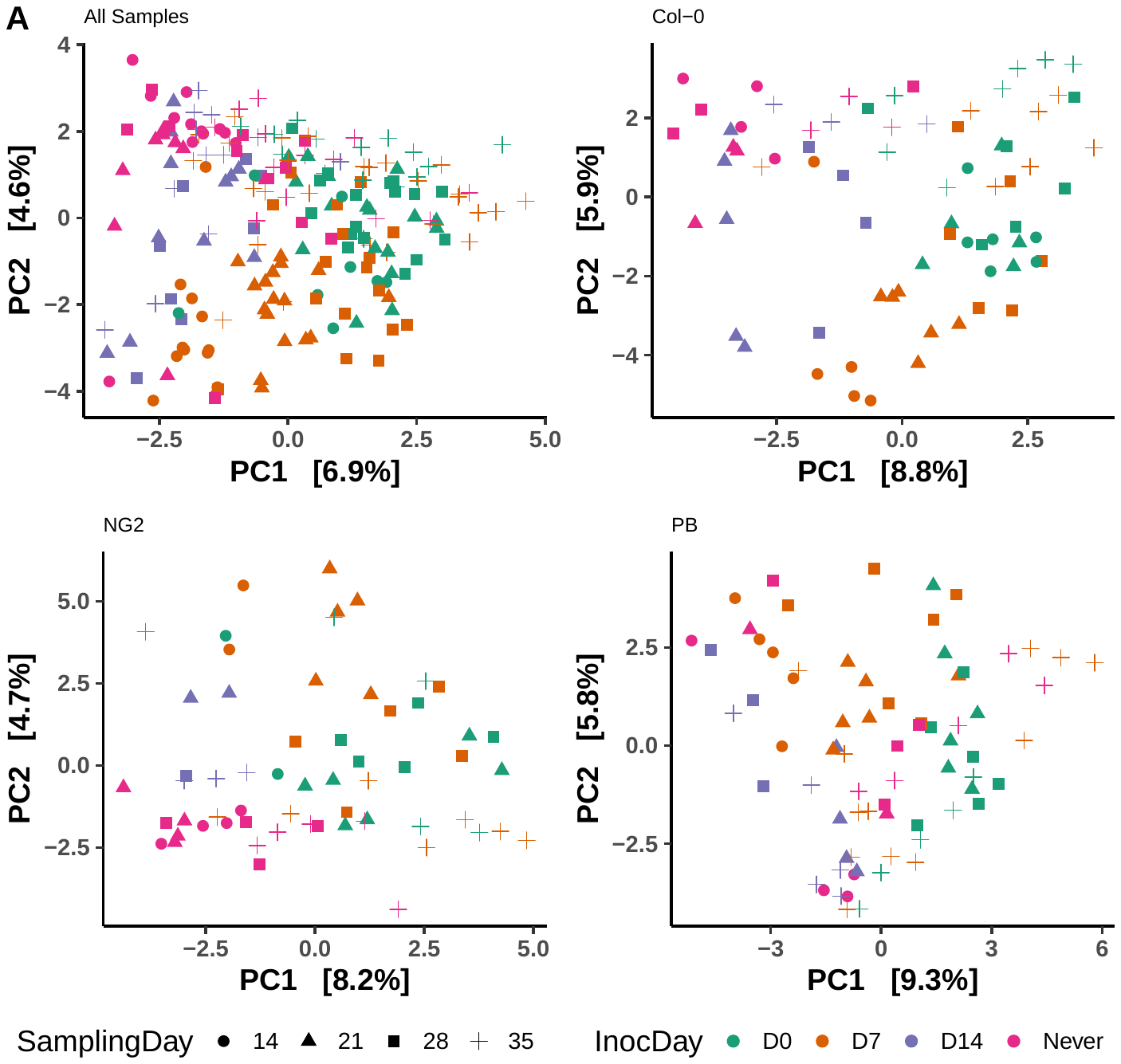 | **C**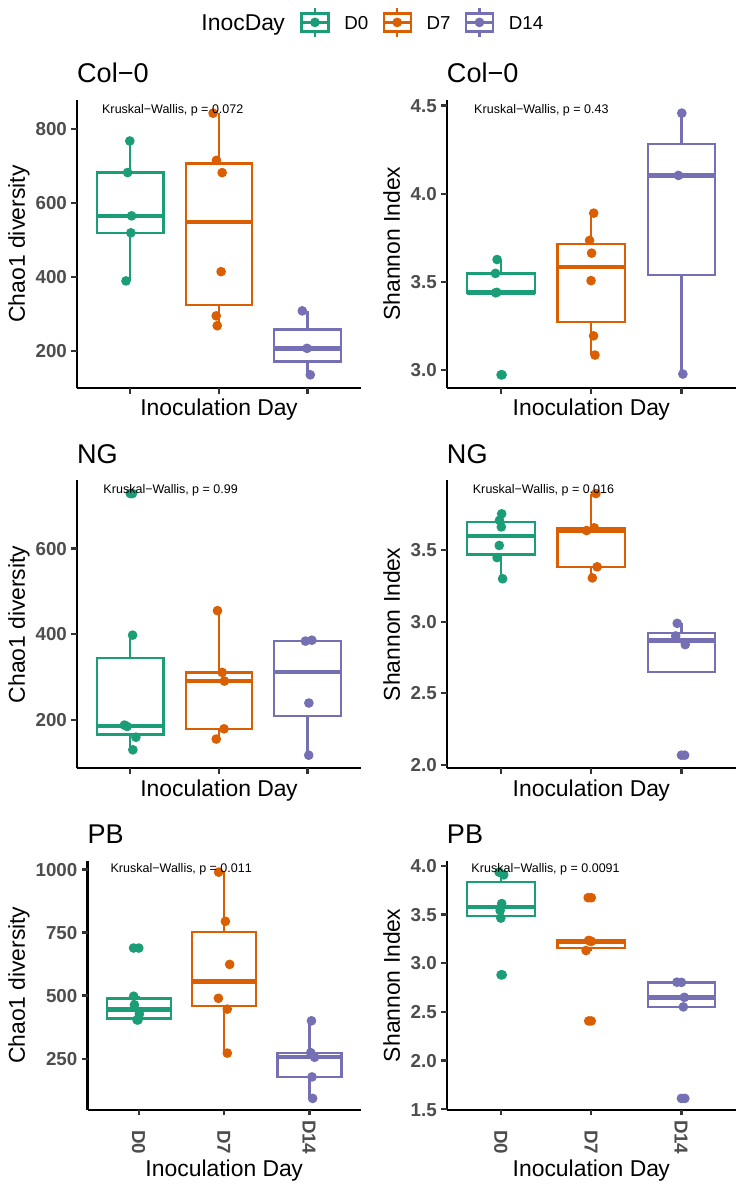 |
| --- | --- |
| 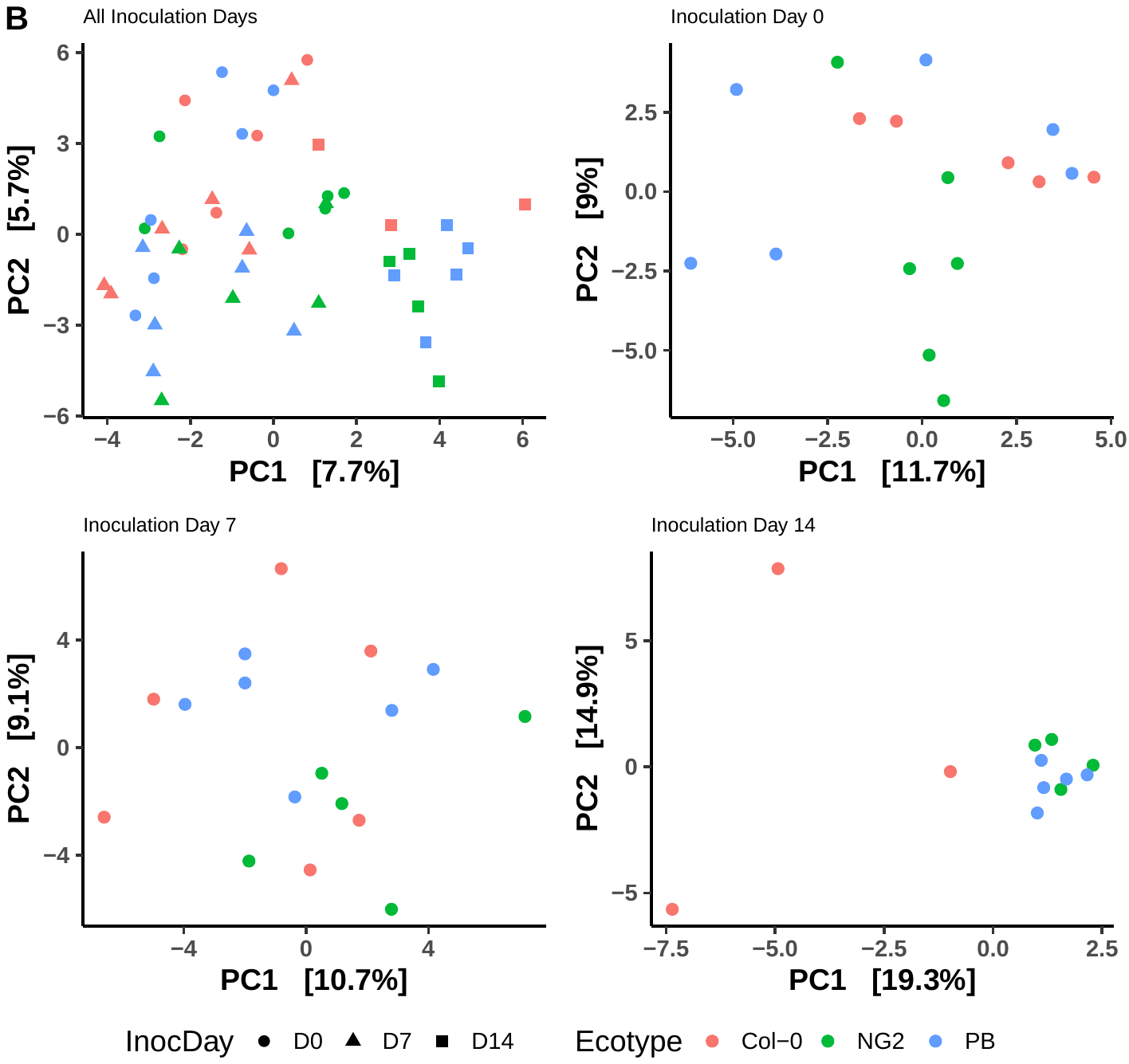 |  |

**Figure S1.** **Early-inoculated plants (before the emergence of true leaves) develop distinct and diverse microbiomes in mature leaves.** Data is based on ASVs associated with the mature leaves of *A. thaliana* after inoculation at day 0, 7 or 14 (D0, D7, D14). **A.** RDA ordination of beta diversity (aitchison distance) of all samples including multiple sampling times. Results of a PERMANOVA test for significance of factors is in Table 1. **B.** RDA ordination of beta diversity (aitchison distance), including only samples collected 21 days post inoculation. Results of a PERMANOVA test for significance of factors is in Table 2. **C.** Alpha diversity (Chao1 estimates of total diversity and Shannon Index) of samples collected 21 days post inoculation**.**

**Figure S2.**

| 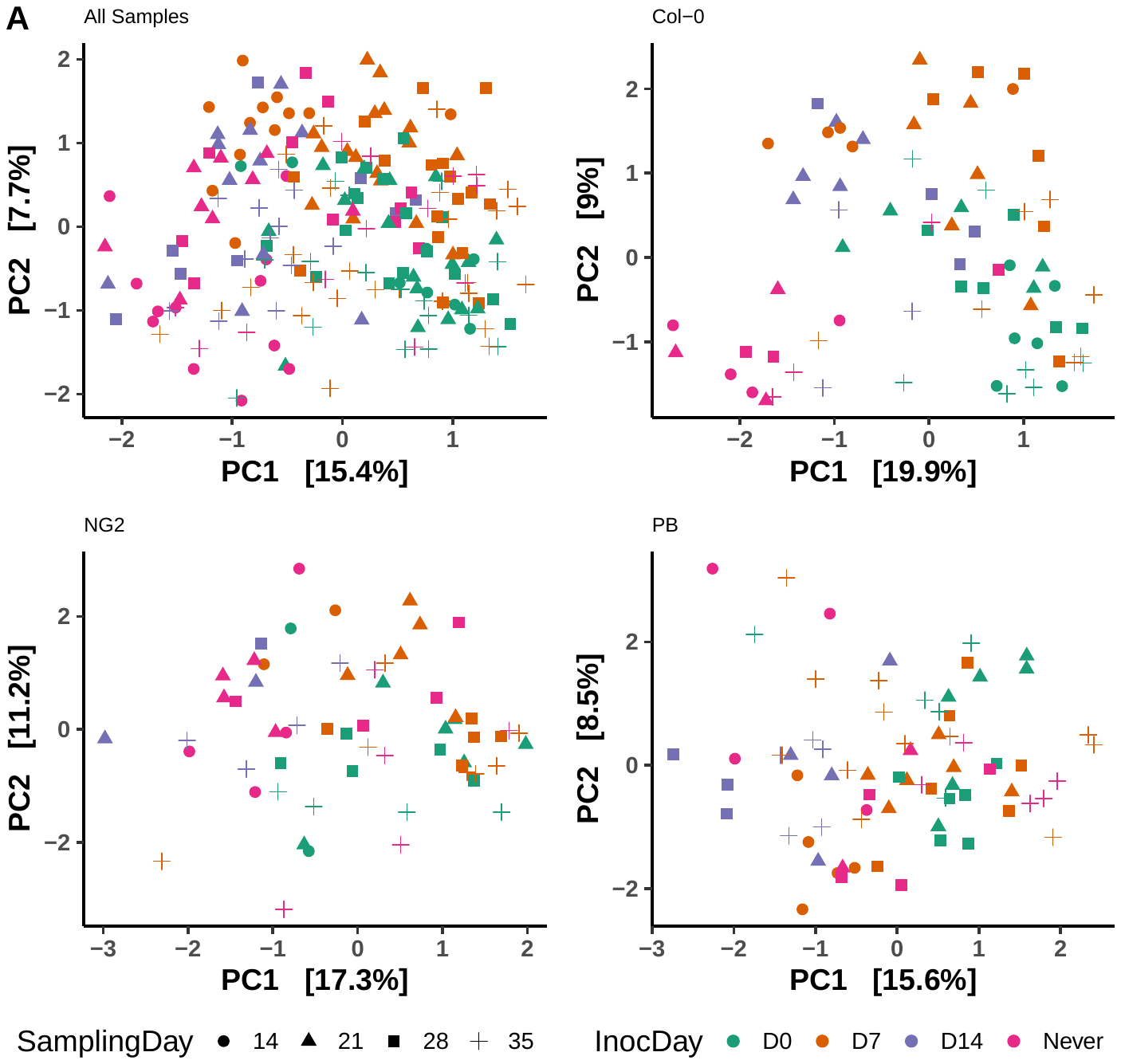 | **C**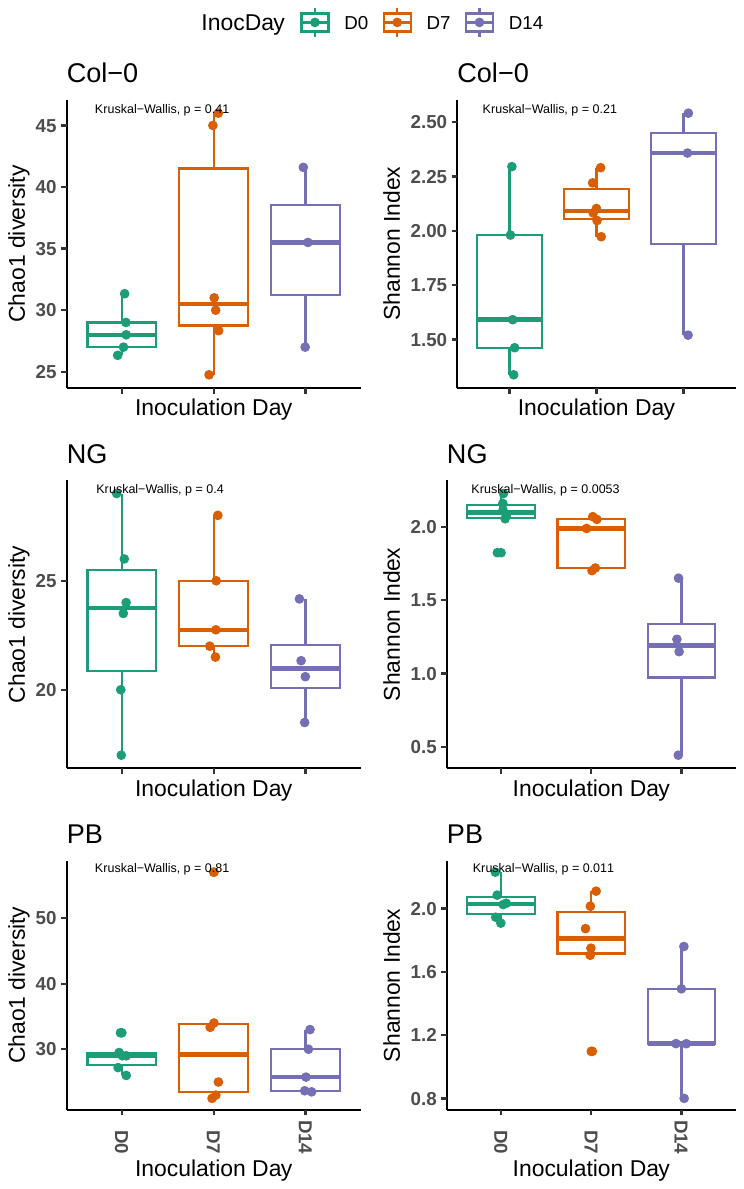 |
| --- | --- |
| 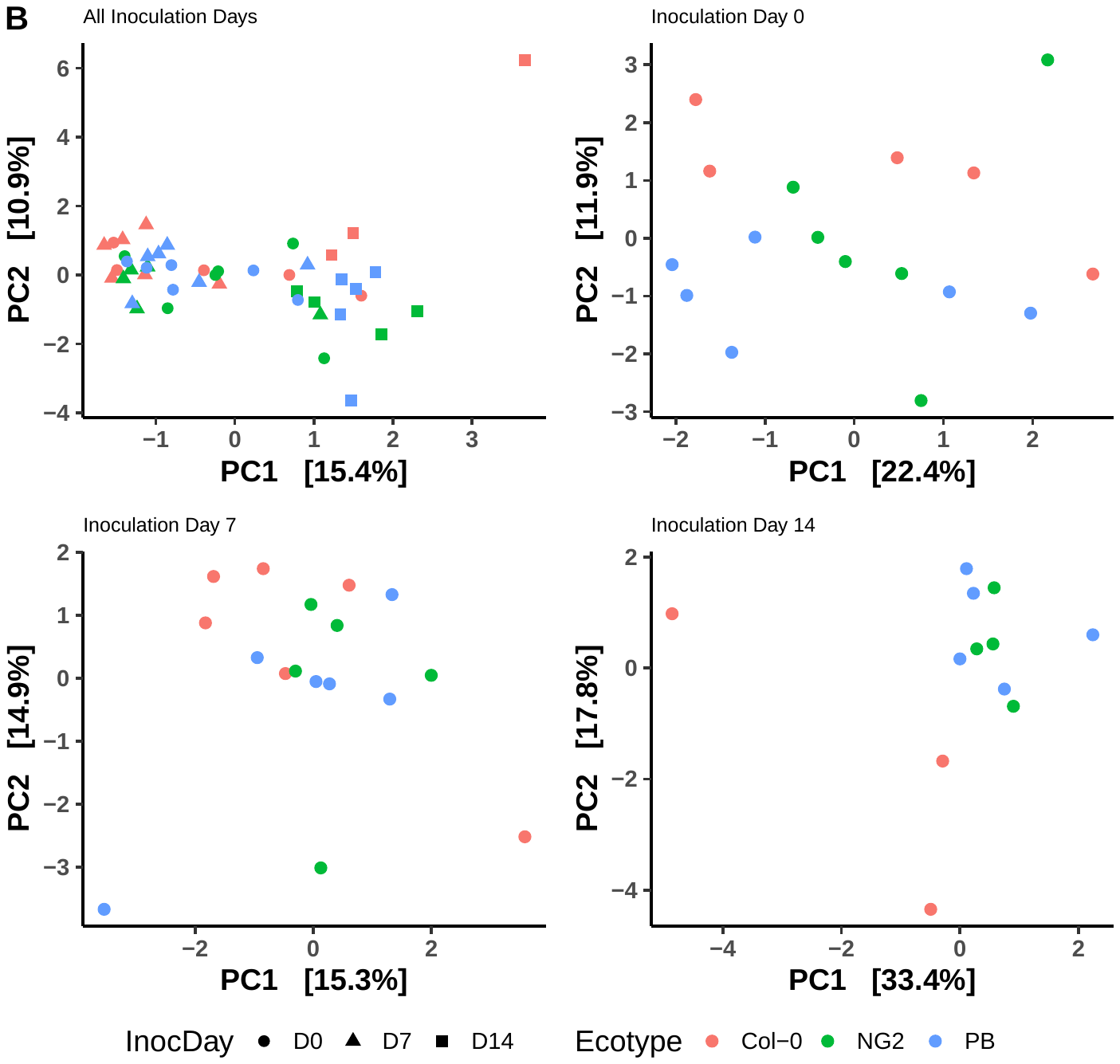 |  |

**Figure S2.** **Early-inoculated plants (before the emergence of true leaves) develop distinct and diverse microbiomes in mature leaves.** Data is based on Order-level taxonomy of bacteria associated with the mature leaves of *A. thaliana* after inoculation at day 0, 7 or 14 (D0, D7, D14). **A.** RDA ordination of beta diversity (aitchison distance) of all samples including multiple sampling times. Results of a PERMANOVA test for significance of factors is in Table 1. **B.** RDA ordination of beta diversity (aitchison distance), including only samples collected 21 days post inoculation. Results of a PERMANOVA test for significance of factors is in Table 2. **C.** Alpha diversity (Chao1 estimates of total diversity and Shannon Index) of samples collected 21 days post inoculation**.**

**Figure S3**

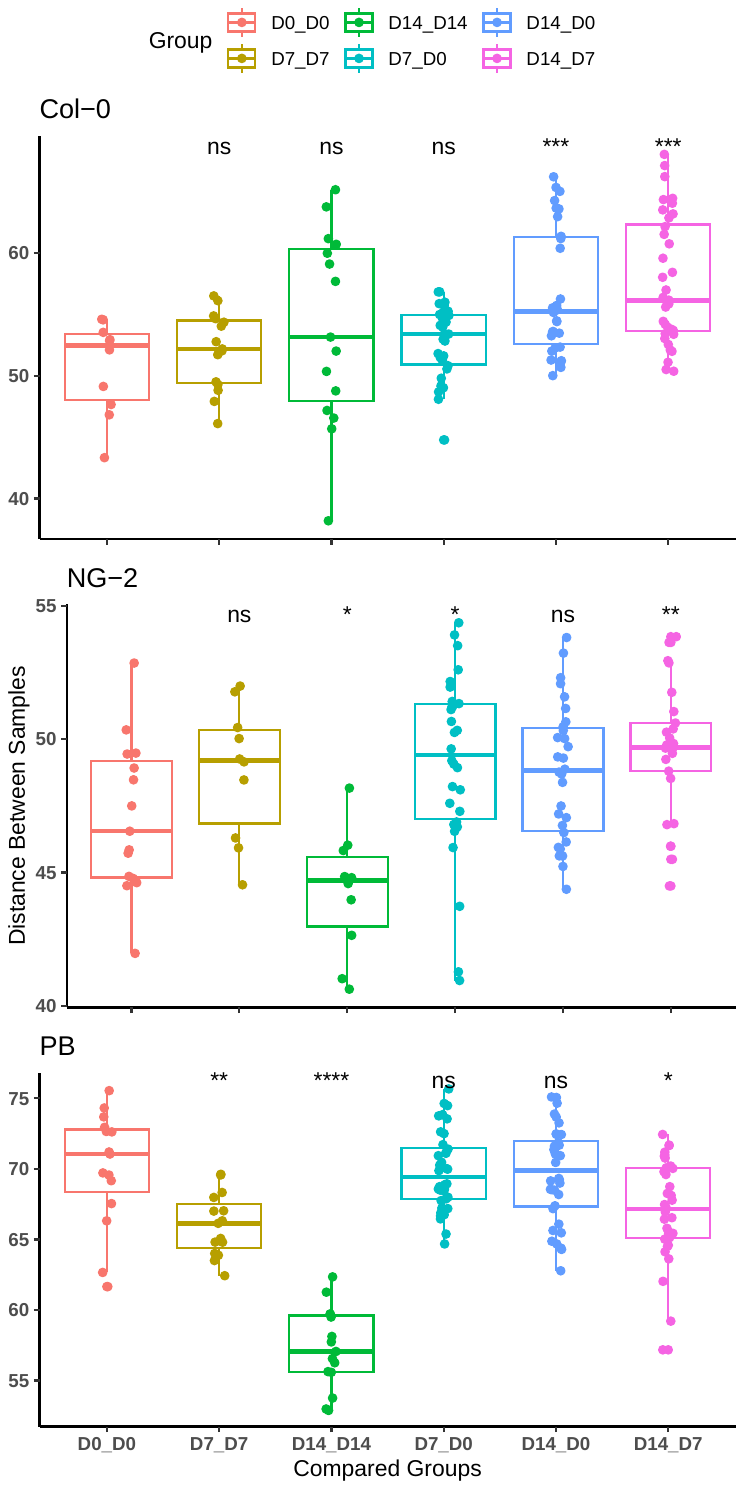

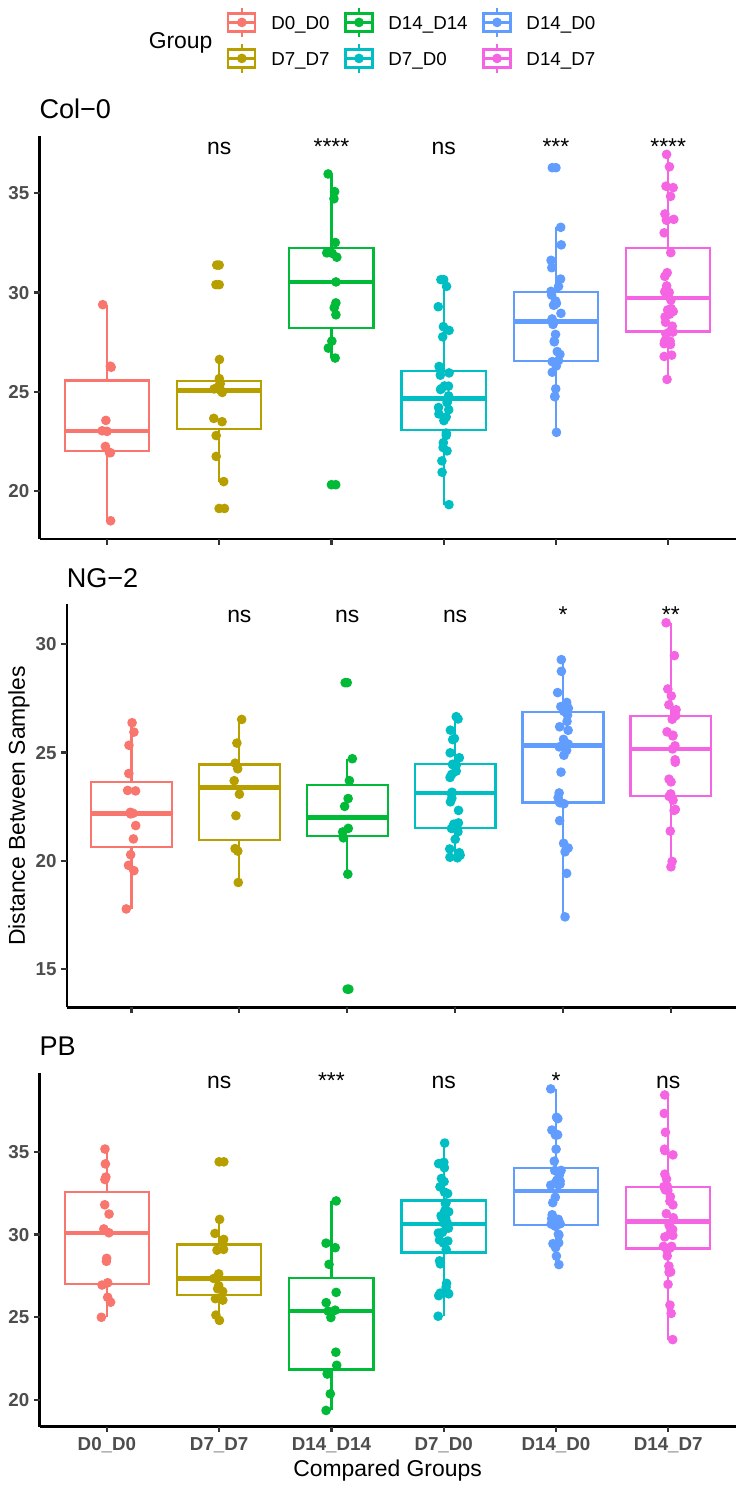

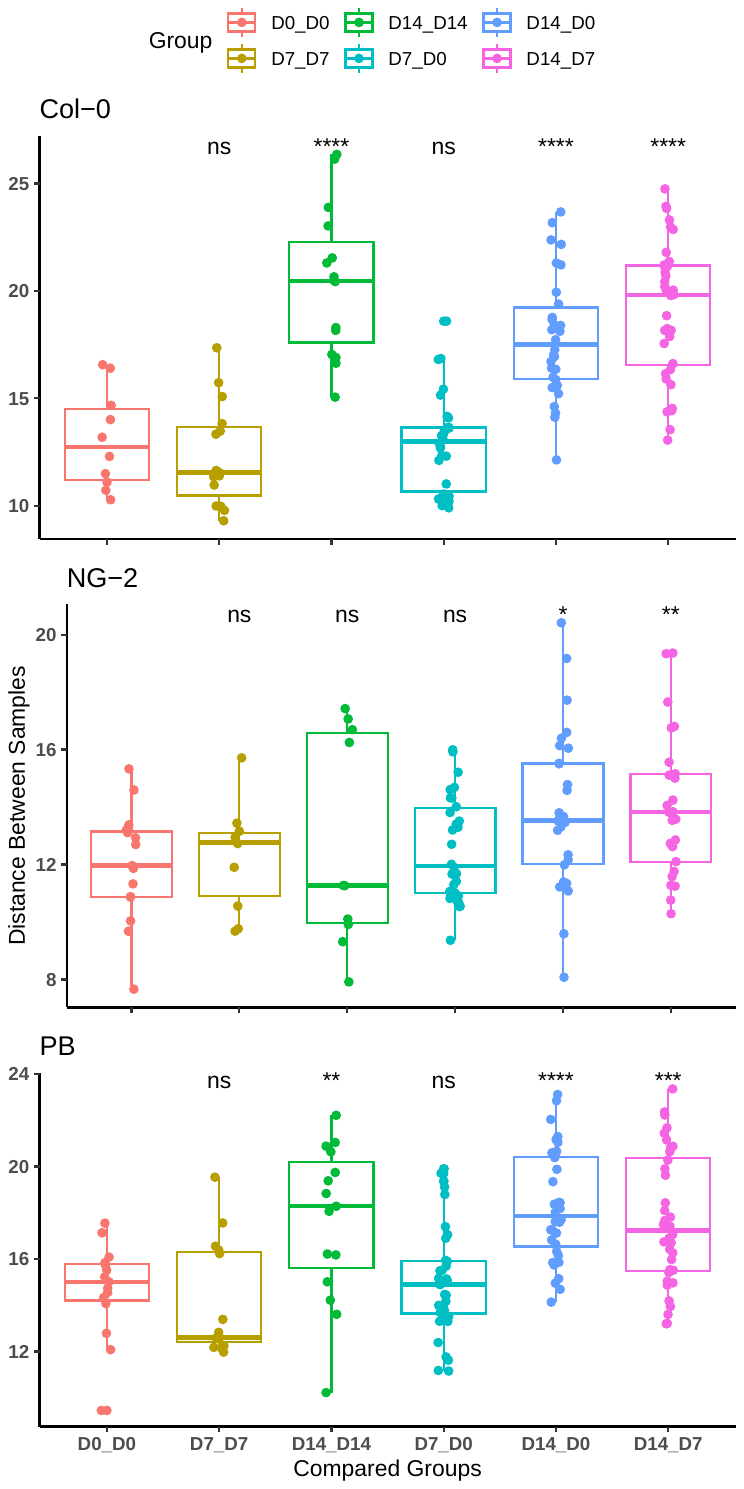

**Figure S3.** Between-sample beta diversity (Aitchison distances) comparing bacterial communities associated with mature *A. thaliana* leaf samples (21 DPI) at the ASV (left), Genus (middle) and Order (right) levels. Within-group distances (D0_D0, D7_D7 or D14_D14) show the variation in the bacterial community structure in a given treatment group. Between-group distances (D7_D0, D14_D0 and D14_D7) show how similar or different samples are between two treatments. In all cases, the stars show sinificant differences compared to the D0_D0 group.

**Figure S4**

A.

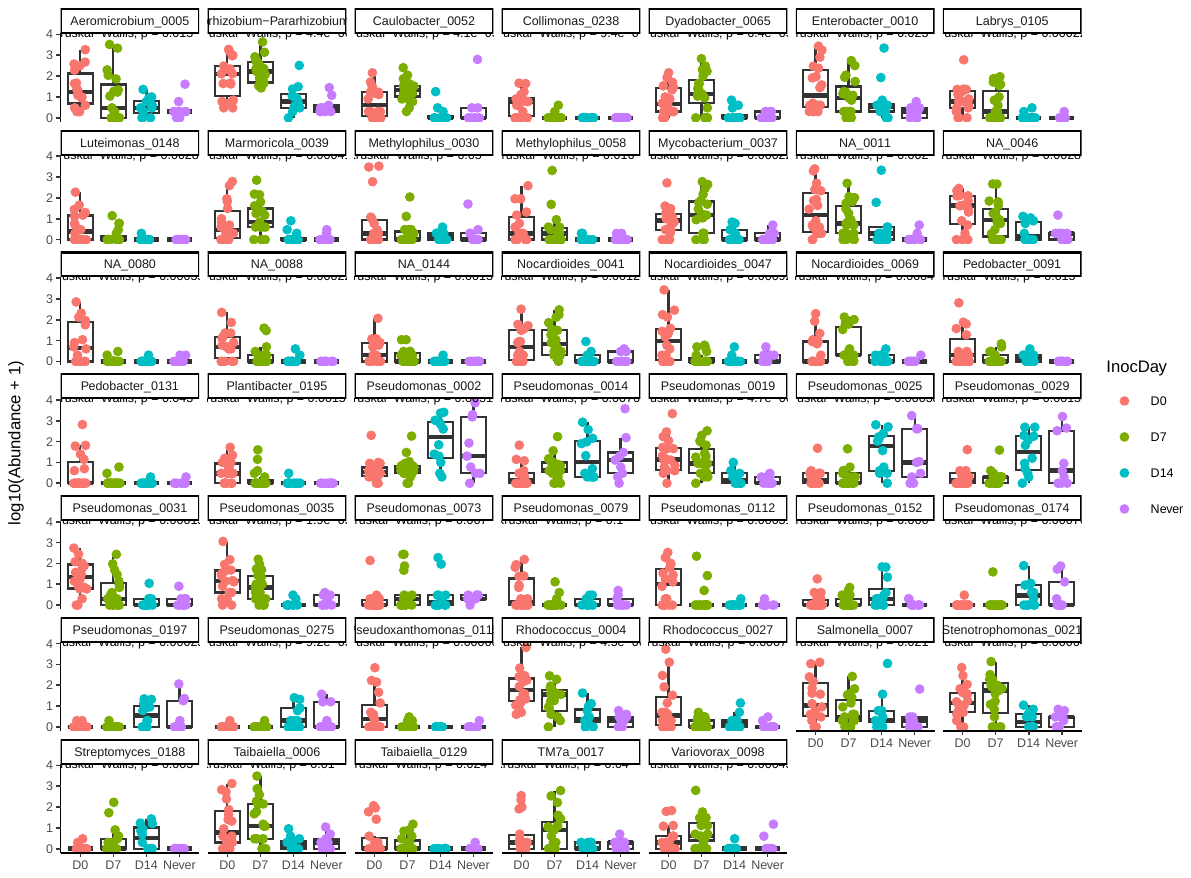

B.

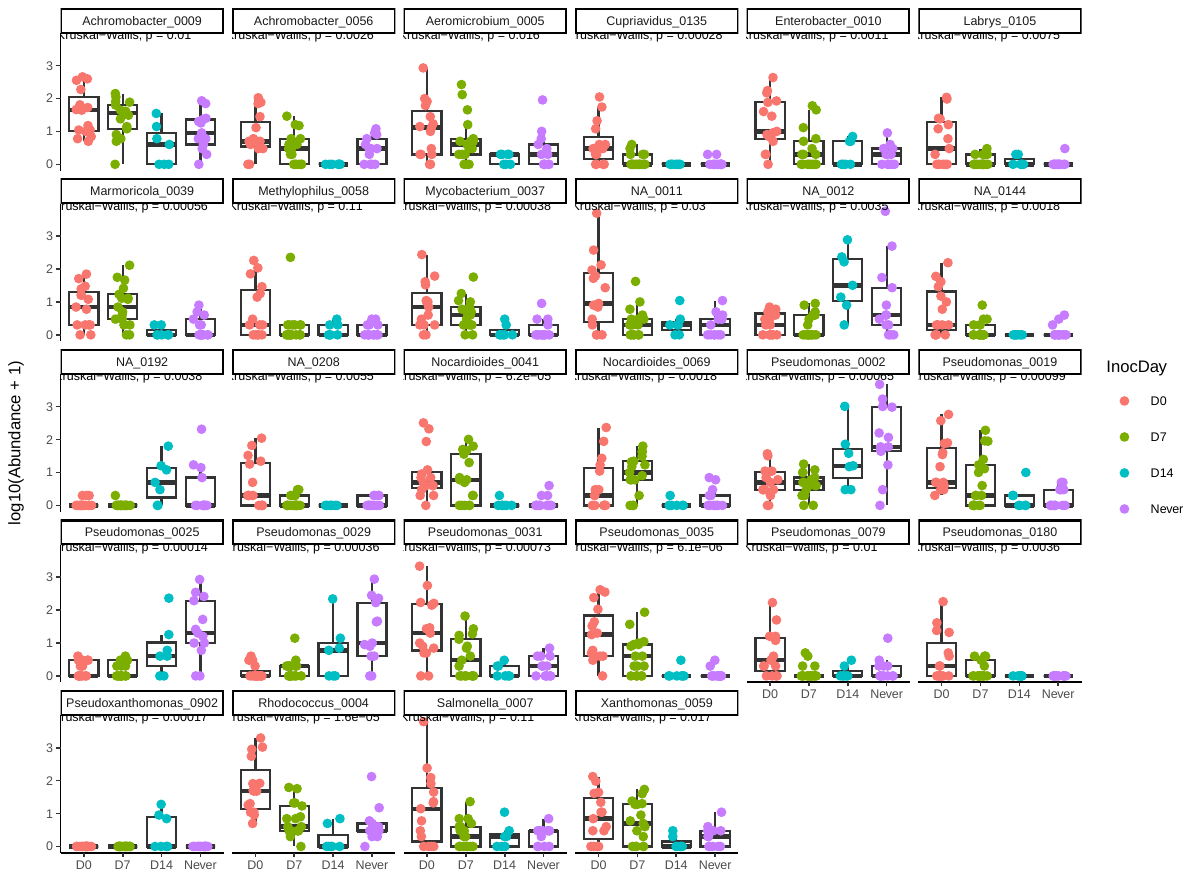

C.

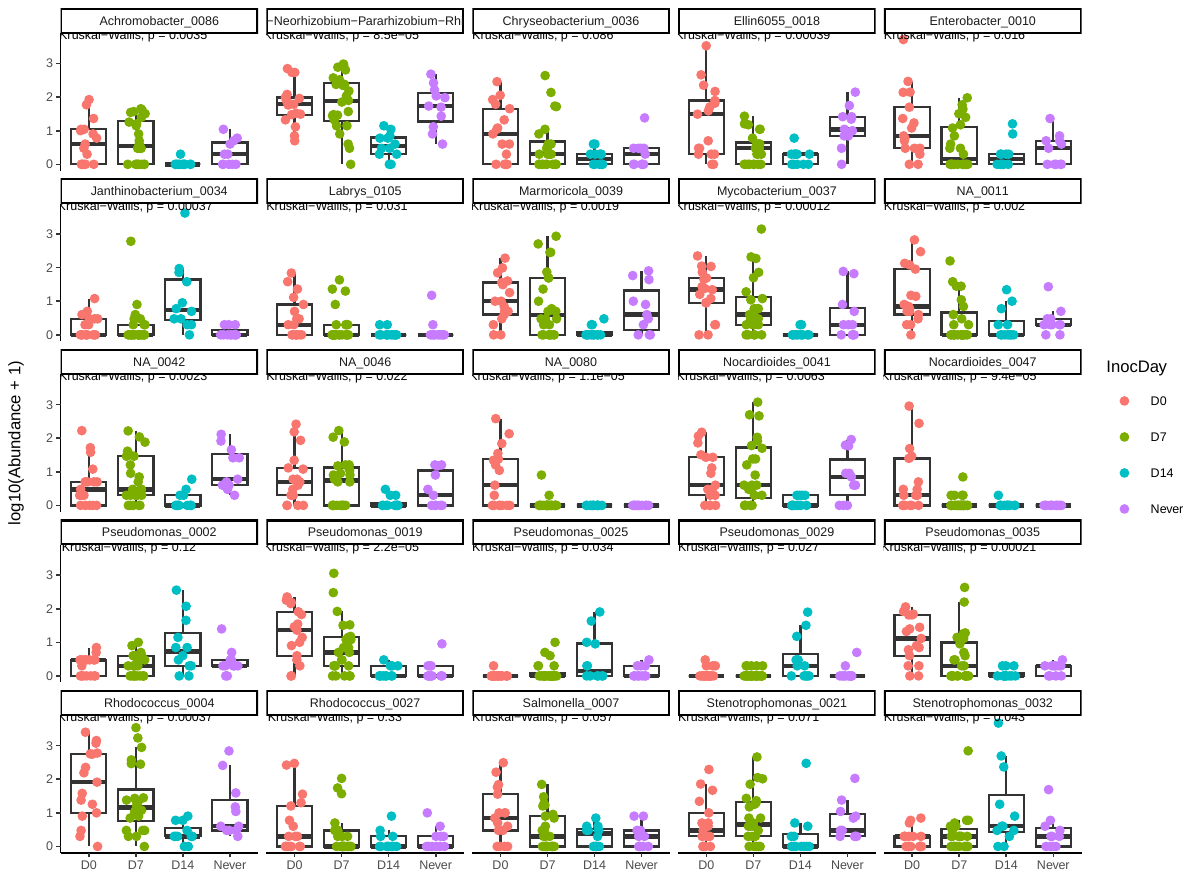

**Figure S4.** Bacterial taxa (ASVs) in mature leaf samples (combined SD21, 28, 35) that were differentially enriched between plants inoculated at D0 and D14. A. in Col-0, B. in NG-2, C. in PB. Detections are based on a DESeq2 analysis with p < 0.001. Kruskall-wallis p-values are additionally shown for each plot.

**Figure S5**

A.	B.

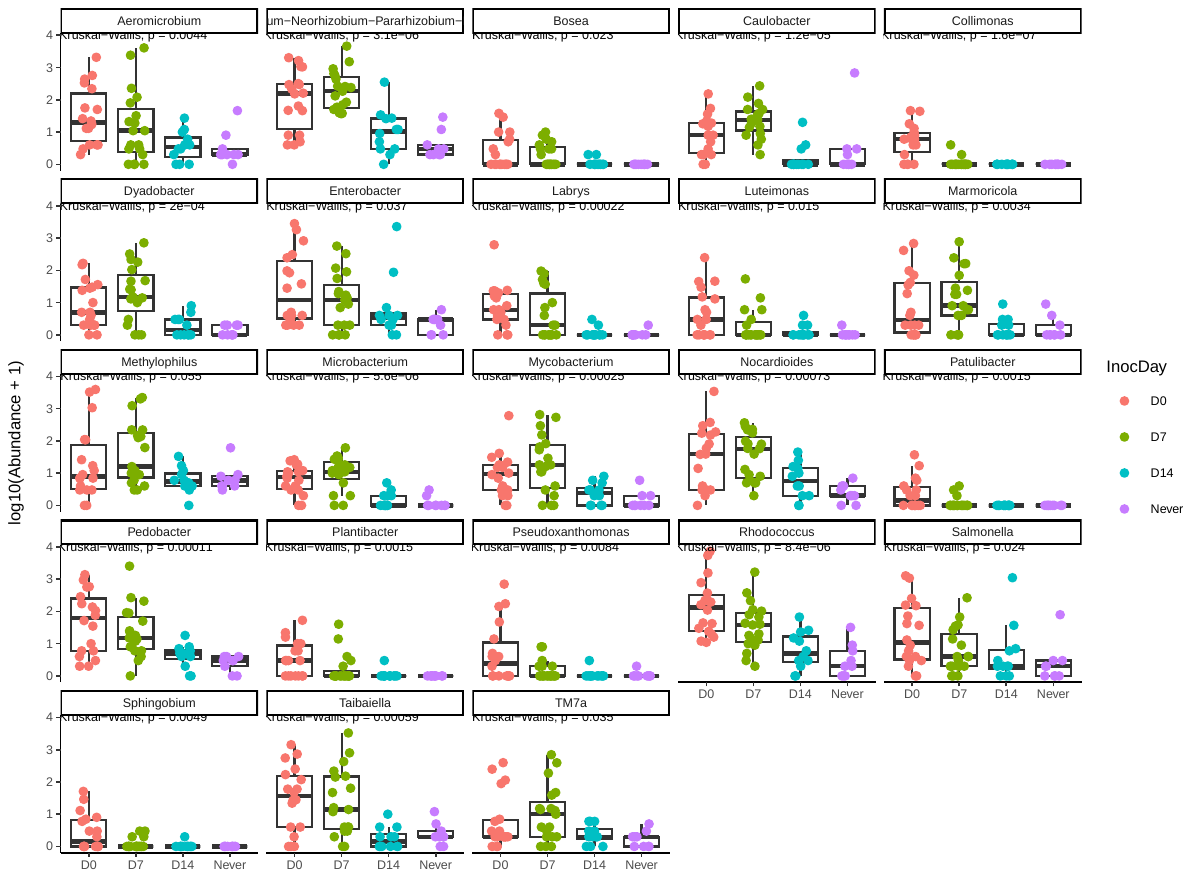

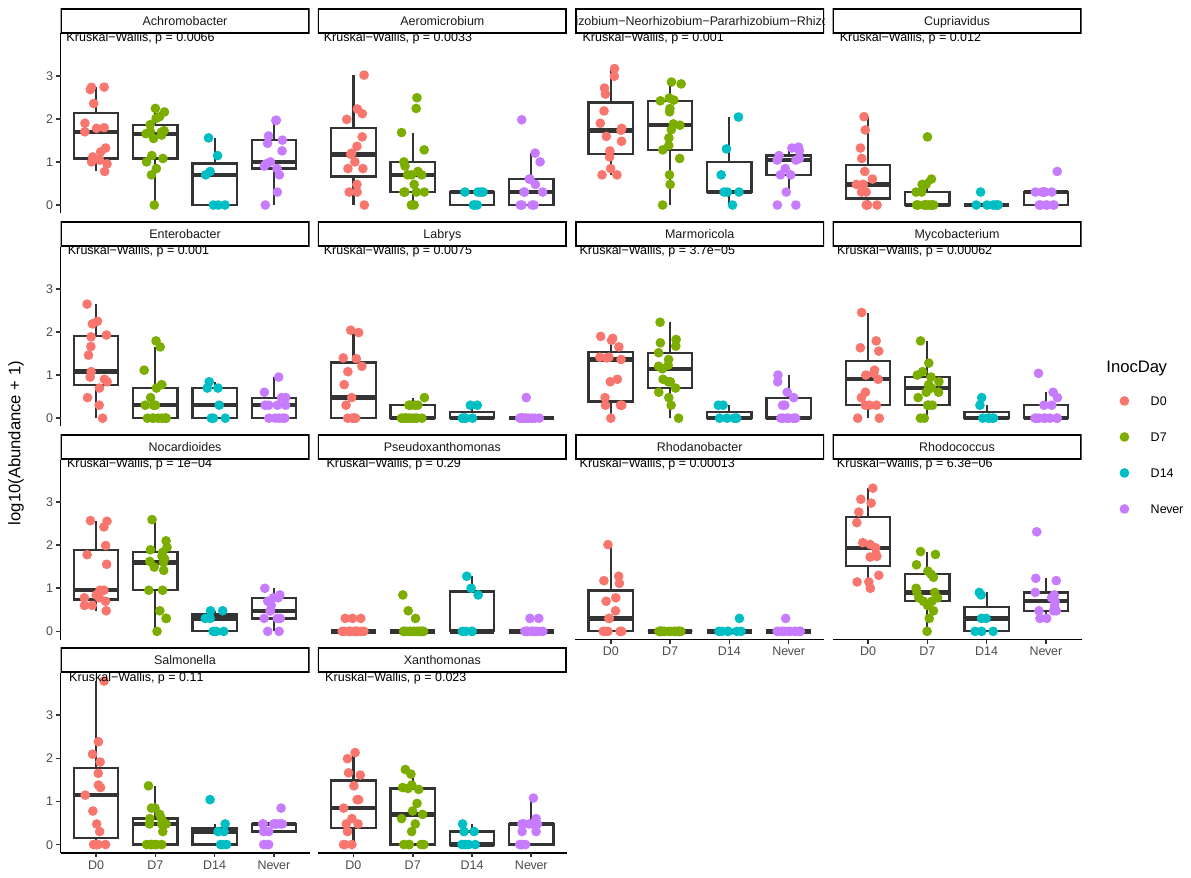

C.

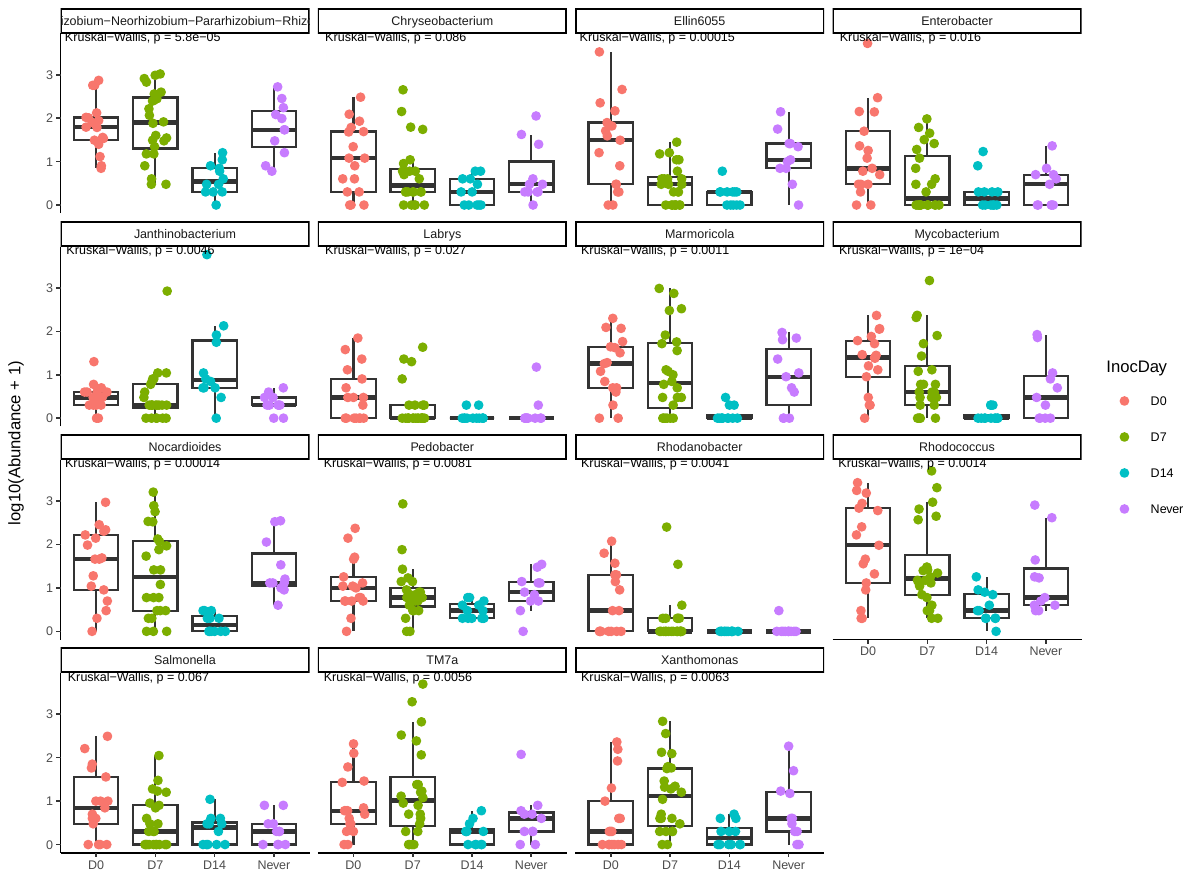

**Figure S5.** Bacterial taxa (Genera) in mature leaf samples (combined SD21, 28, 35) that were differentially enriched between plants inoculated at D0 and D14. A. in Col-0, B. in NG-2, C. in PB. Detections are based on DESeq analysis with p < 0.001. Kruskall-wallis p-values are additionally shown for each plot.

**Figure S6**

A.	B.

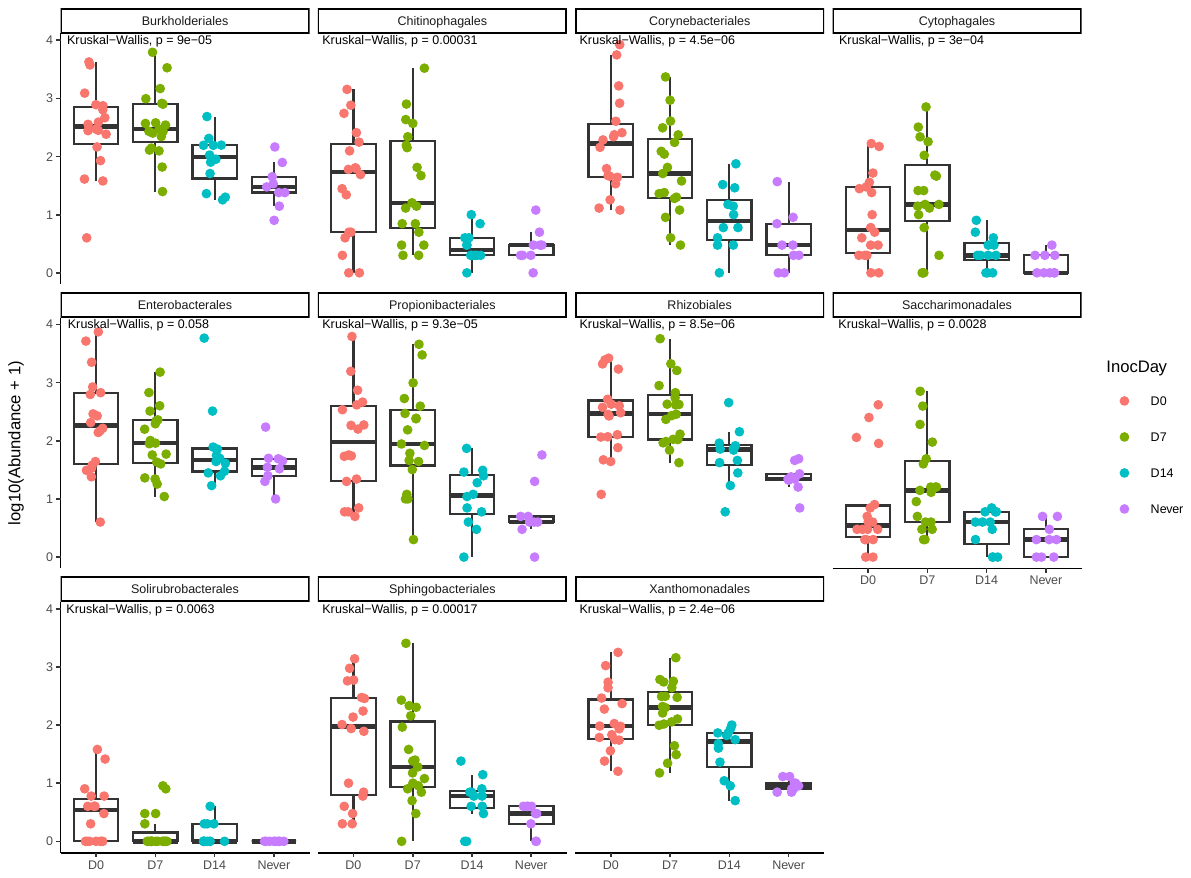

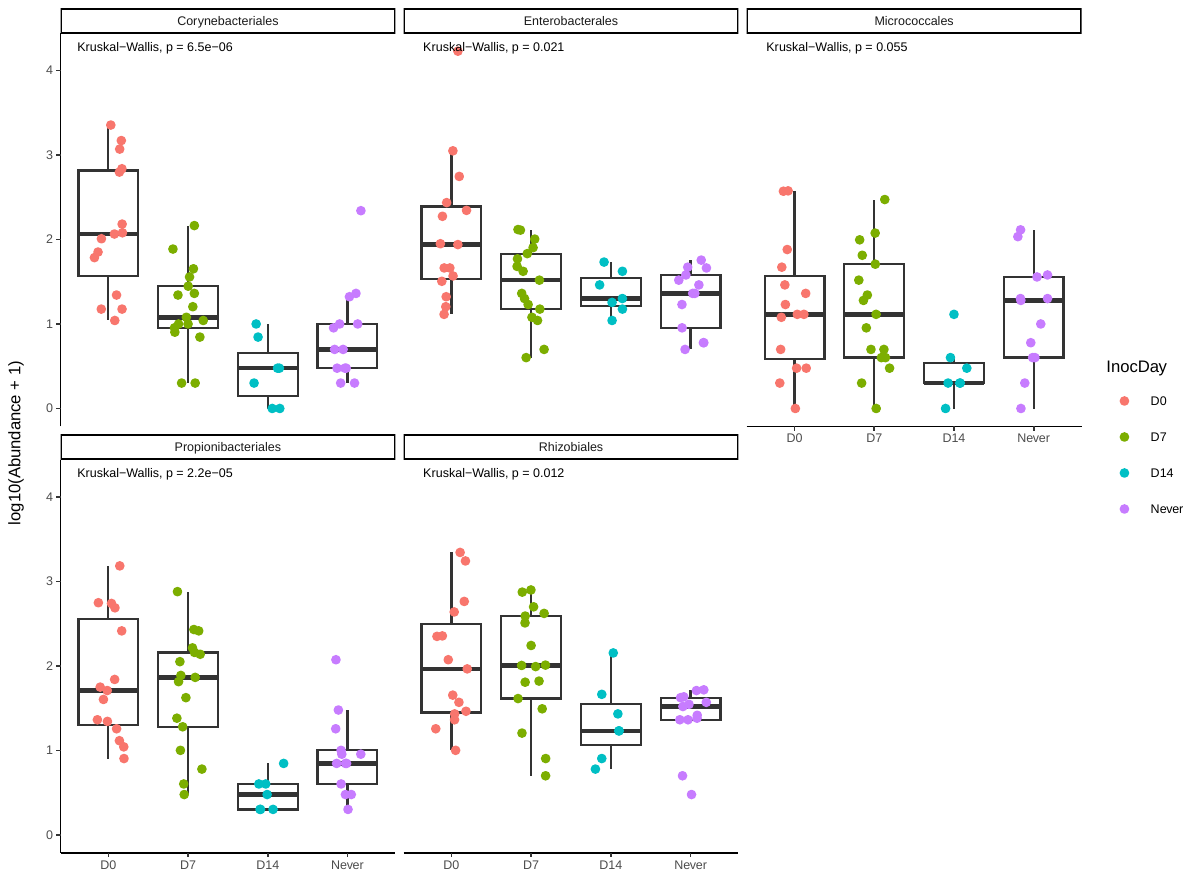

**Figure S6.** Bacterial taxa (Order-level) in mature leaf samples (combined SD21, 28, 35) that were differentially enriched between plants inoculated at D0 and D14. A. in Col-0, B. in NG-2 (no differentially abundant Order-level taxa were detected in PB). Detections are based on DESeq analysis with p < 0.001. Kruskall-wallis p-values are additionally shown for each plot.

**Figure S7**

A.	B.	C.

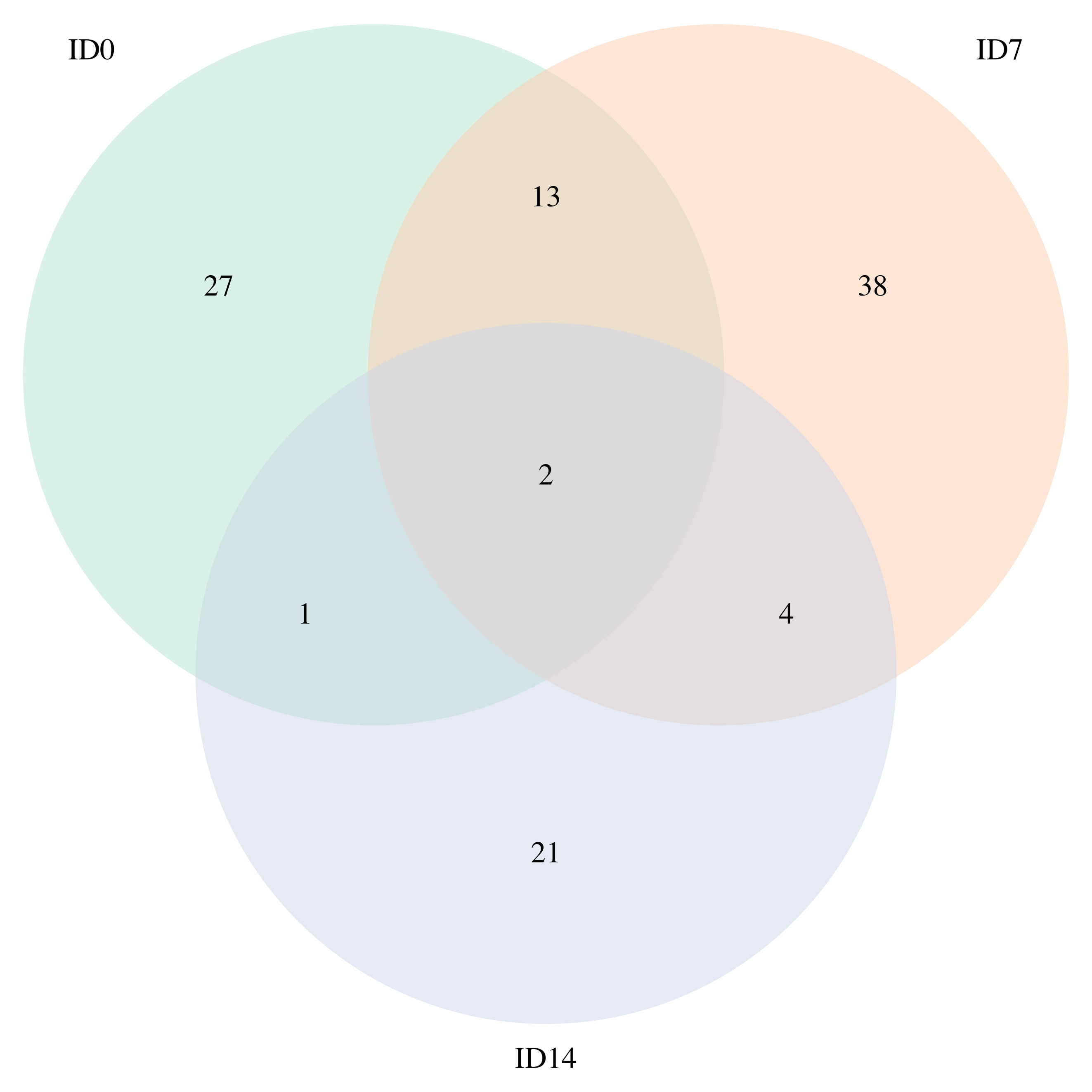

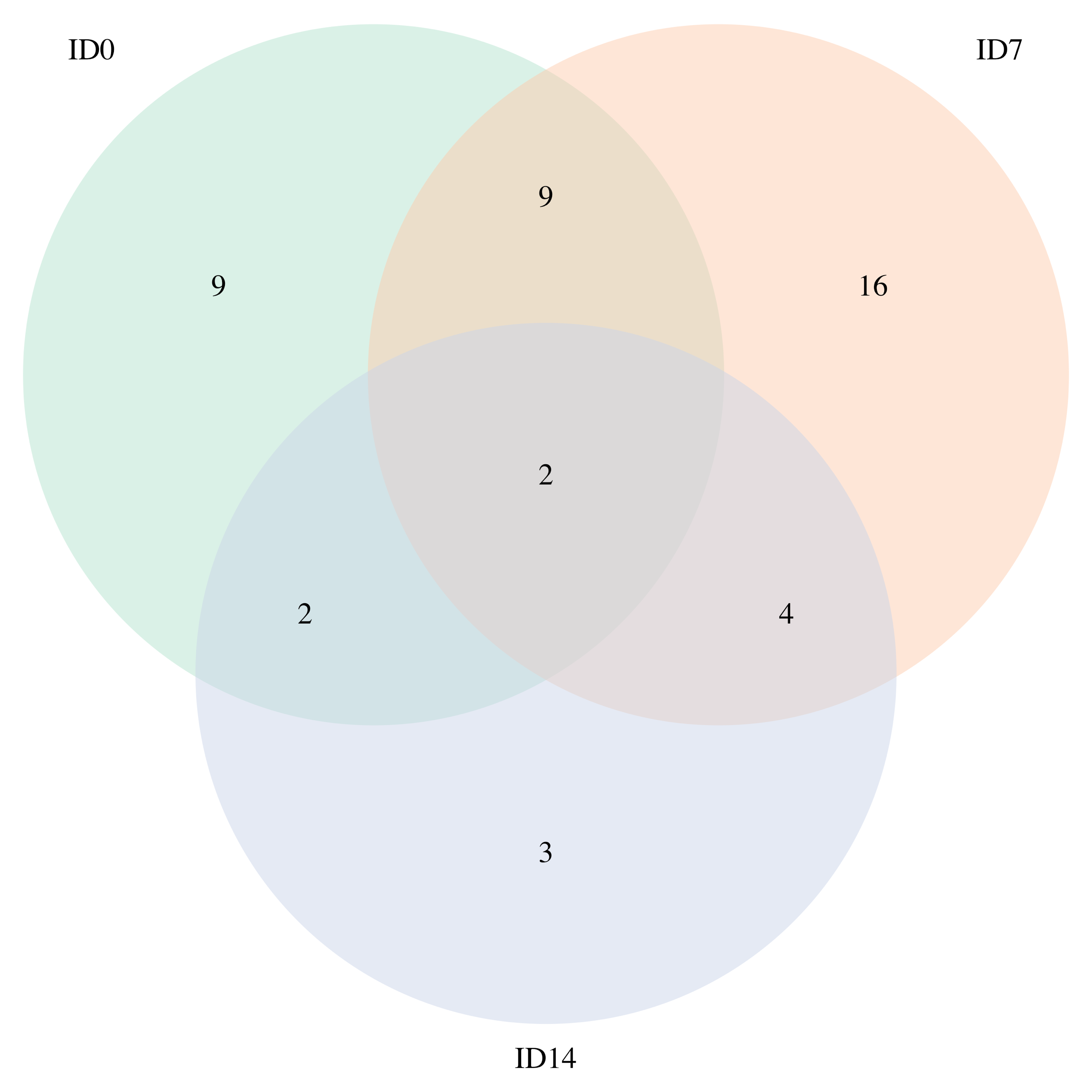

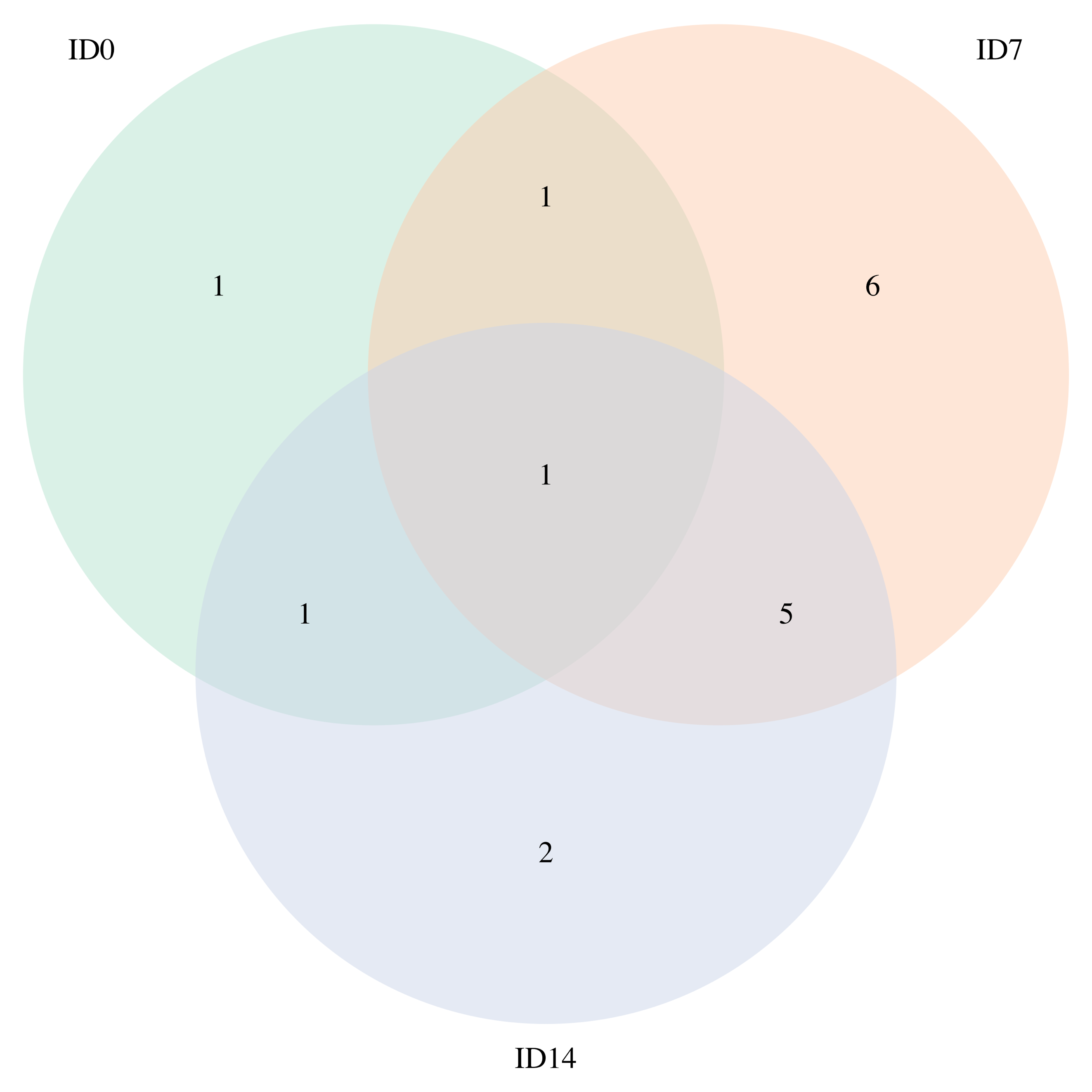

**Figure S7.** Number of taxa correlated to ecotype in mature plants (21 days post inoculation) that had been inoculated at D0, D7 or D14. A. ASV-level taxa, B. Genus-level taxa, and C. Order-level taxa. Detections were based on DESeq analysis with p < 0.01.

**Figure S8**

A.

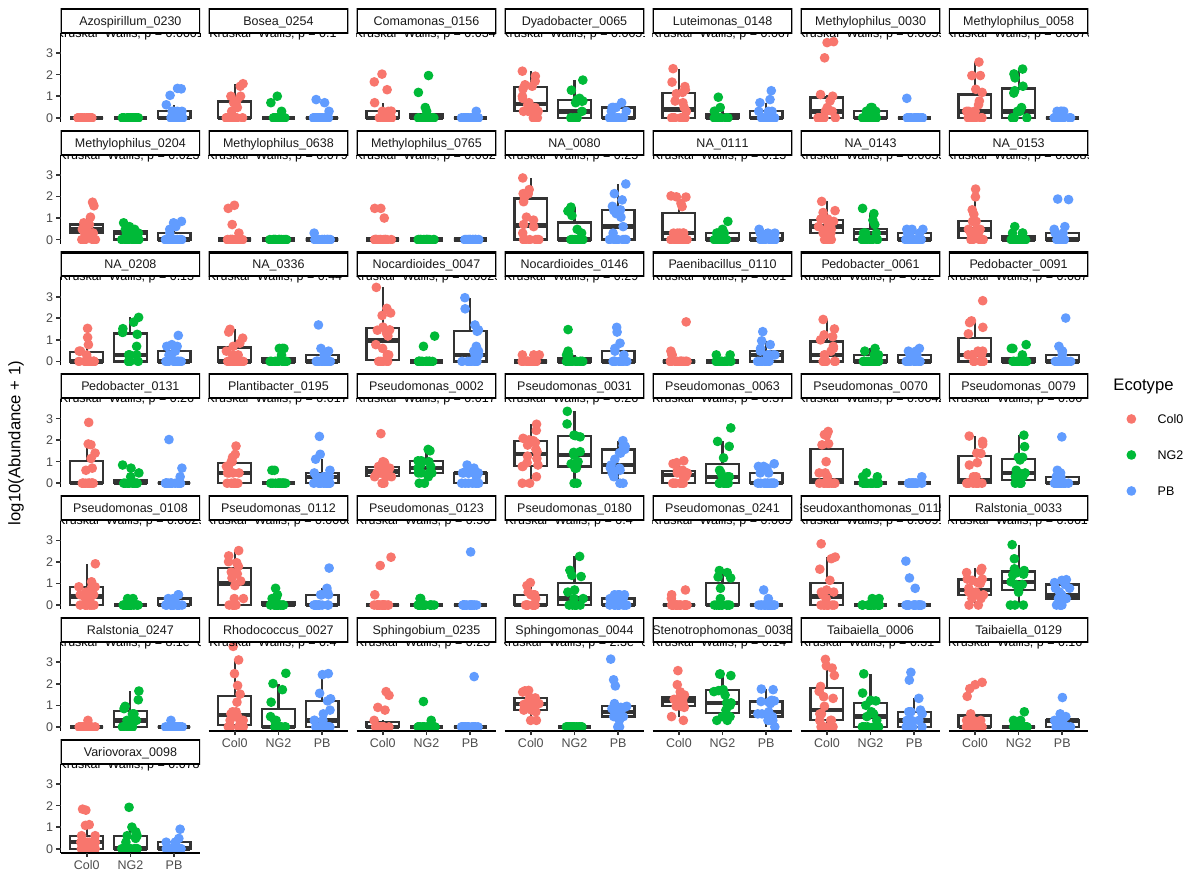

B.

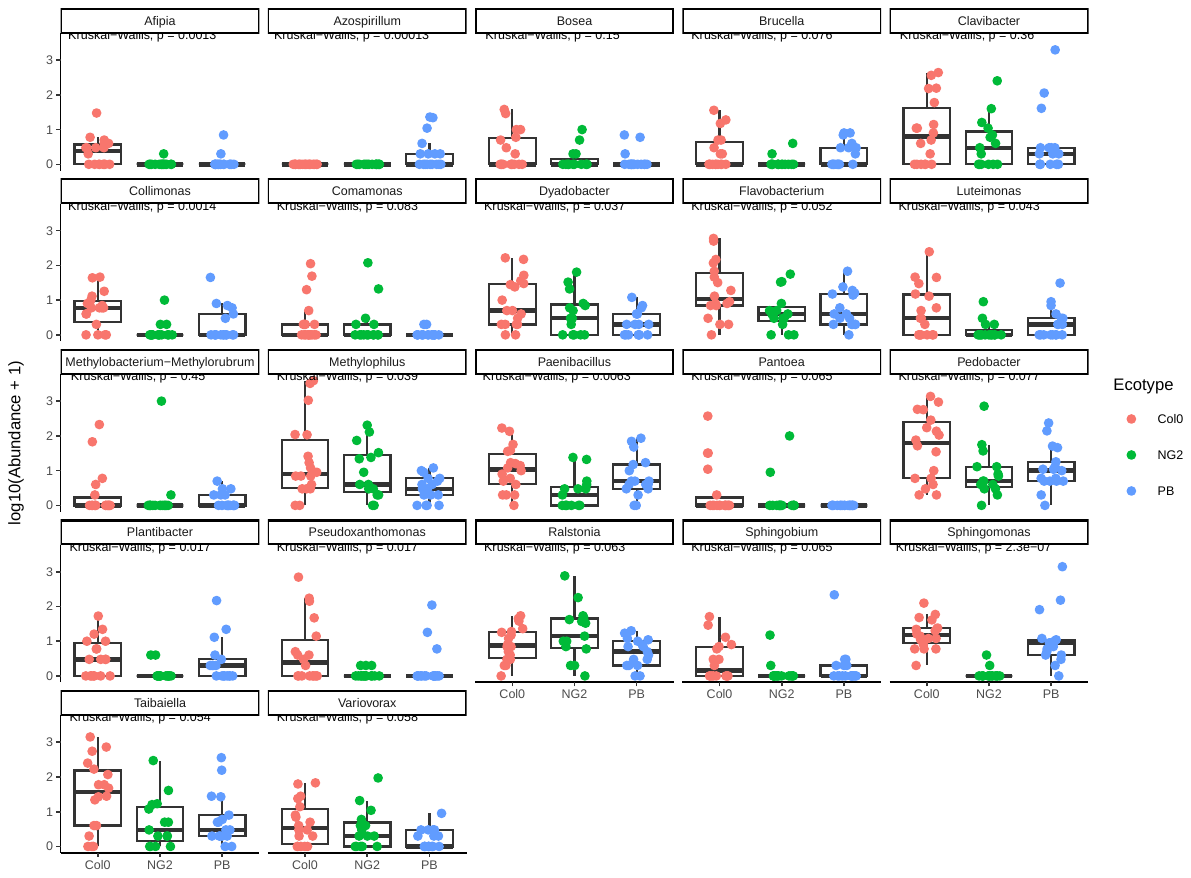

C.

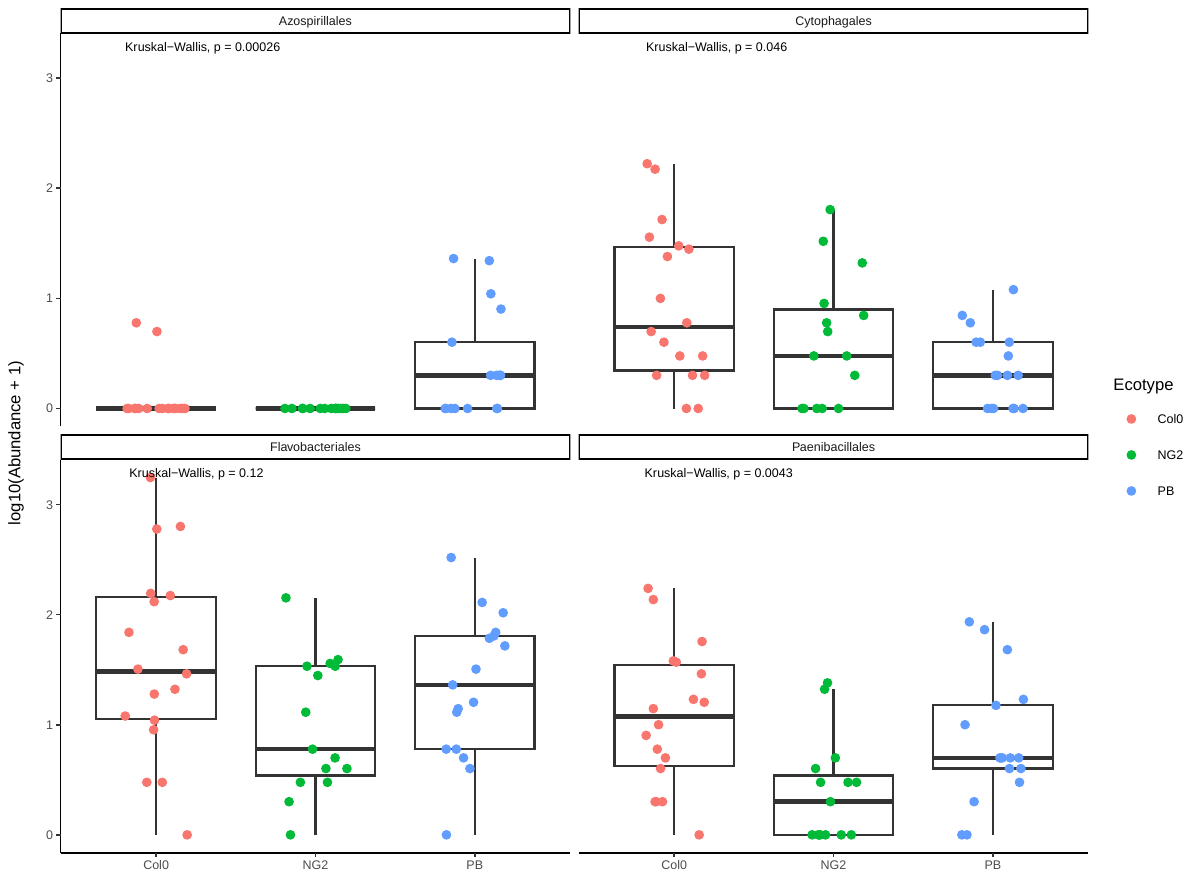

**Figure S8.** Bacterial taxa that were differentially enriched between ecotypes in mature leaf samples (combined SD21, 28, 35) in plants inoculated at D0. A. ASV-level taxa, B. Genus-level taxa, and C. Order-level taxa. Detections are based on DESeq analysis with p < 0.01. Kruskall-wallis p-values are additionally shown for each plot.

**Figure S9**

A.	B.

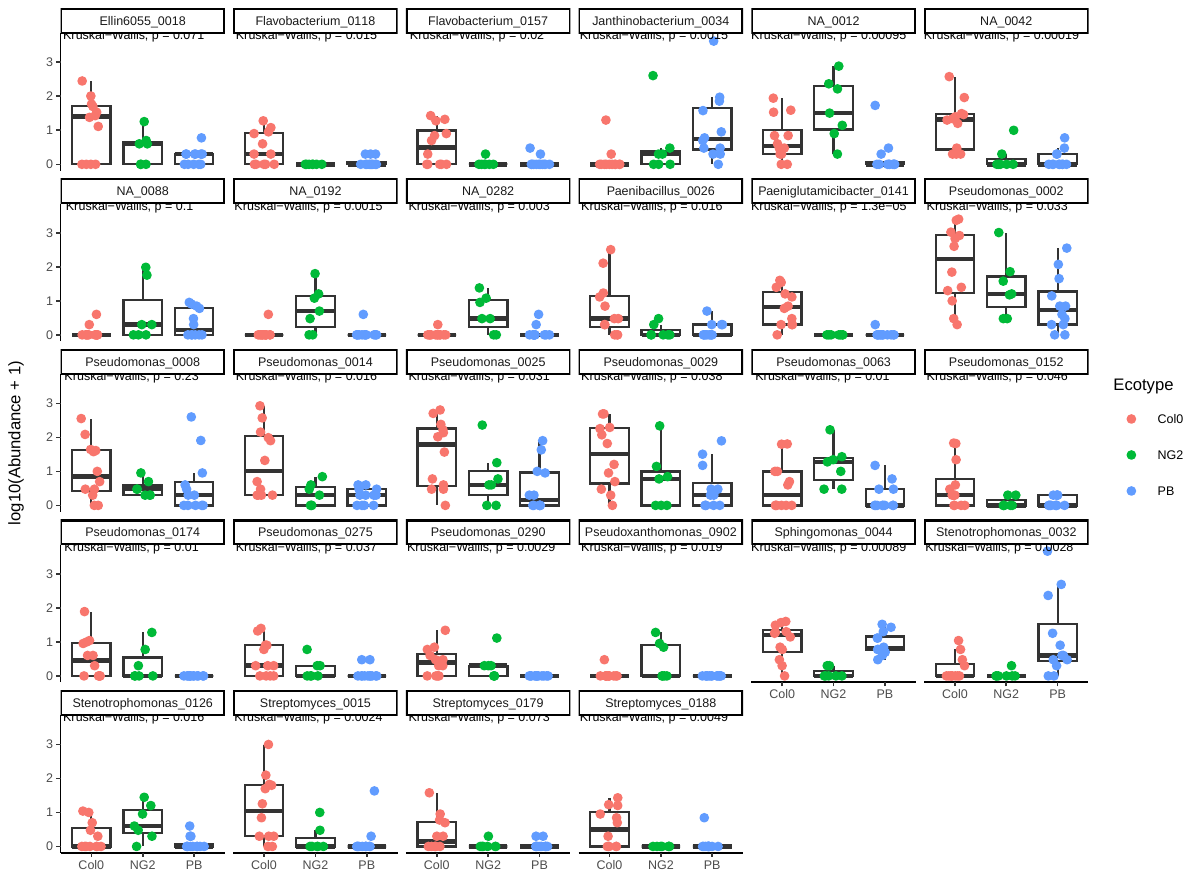

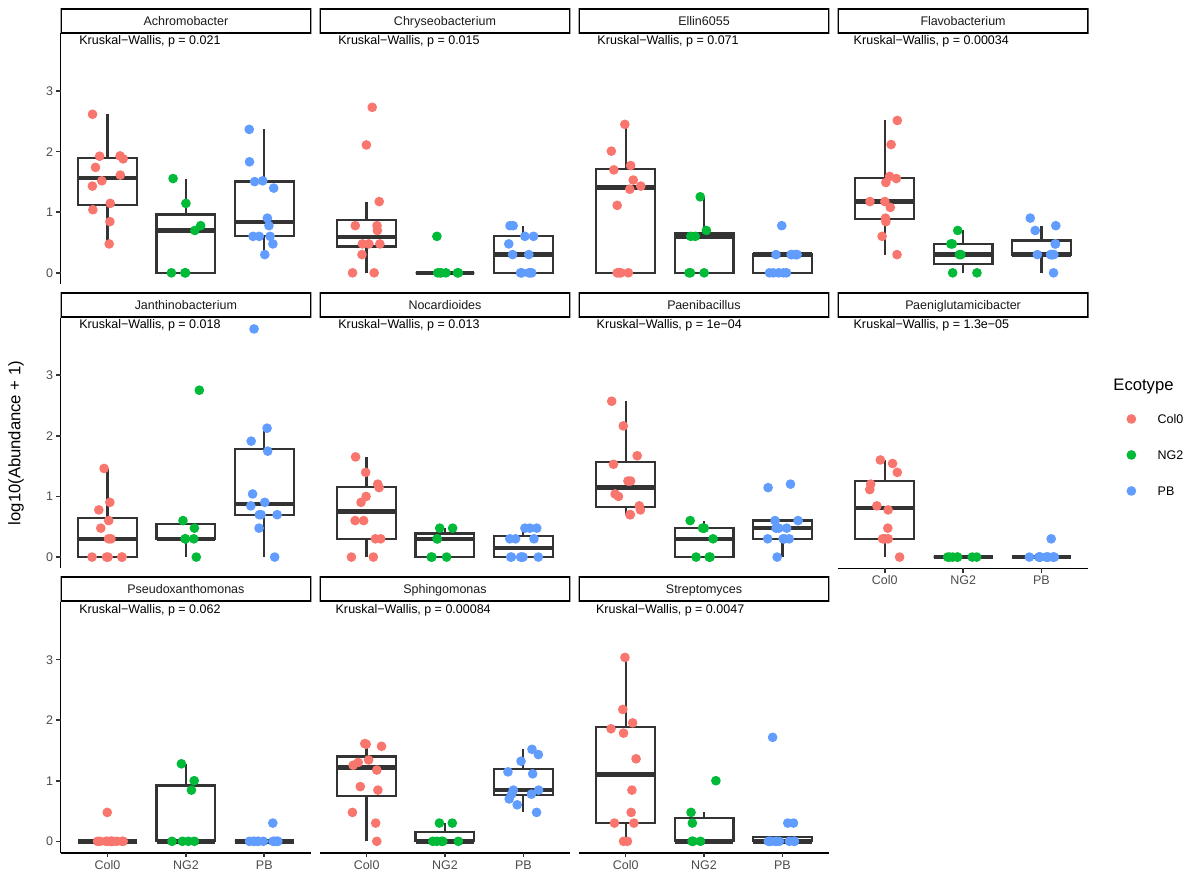

C.

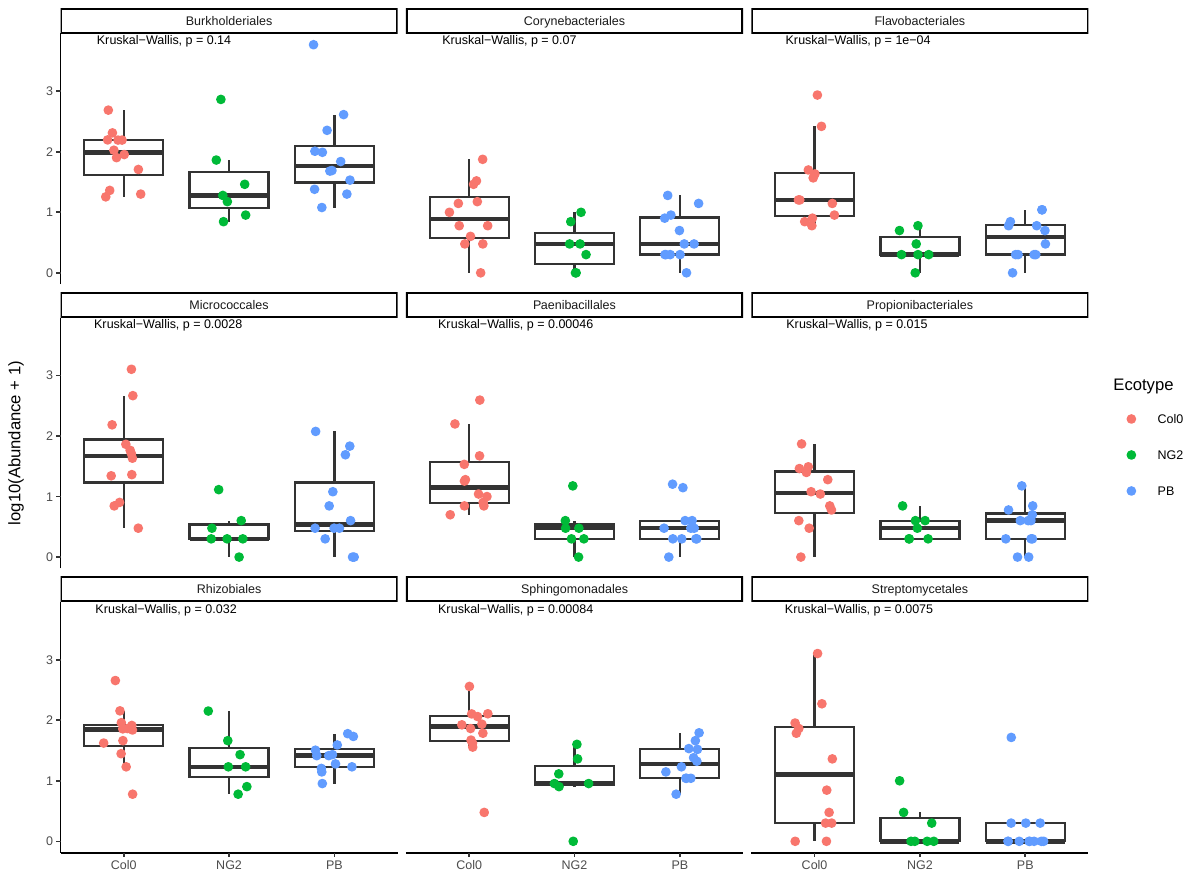

**Figure S9.** Bacterial taxa that were differentially enriched between ecotypes in mature leaf samples (combined SD21, 28, 35) in plants inoculated at D14. A. ASV-level taxa, B. Genus-level taxa, and C. Order-level taxa. Detections are based on DESeq analysis with p < 0.01. Kruskall-wallis p-values are additionally shown for each plot.

**Figure S10**

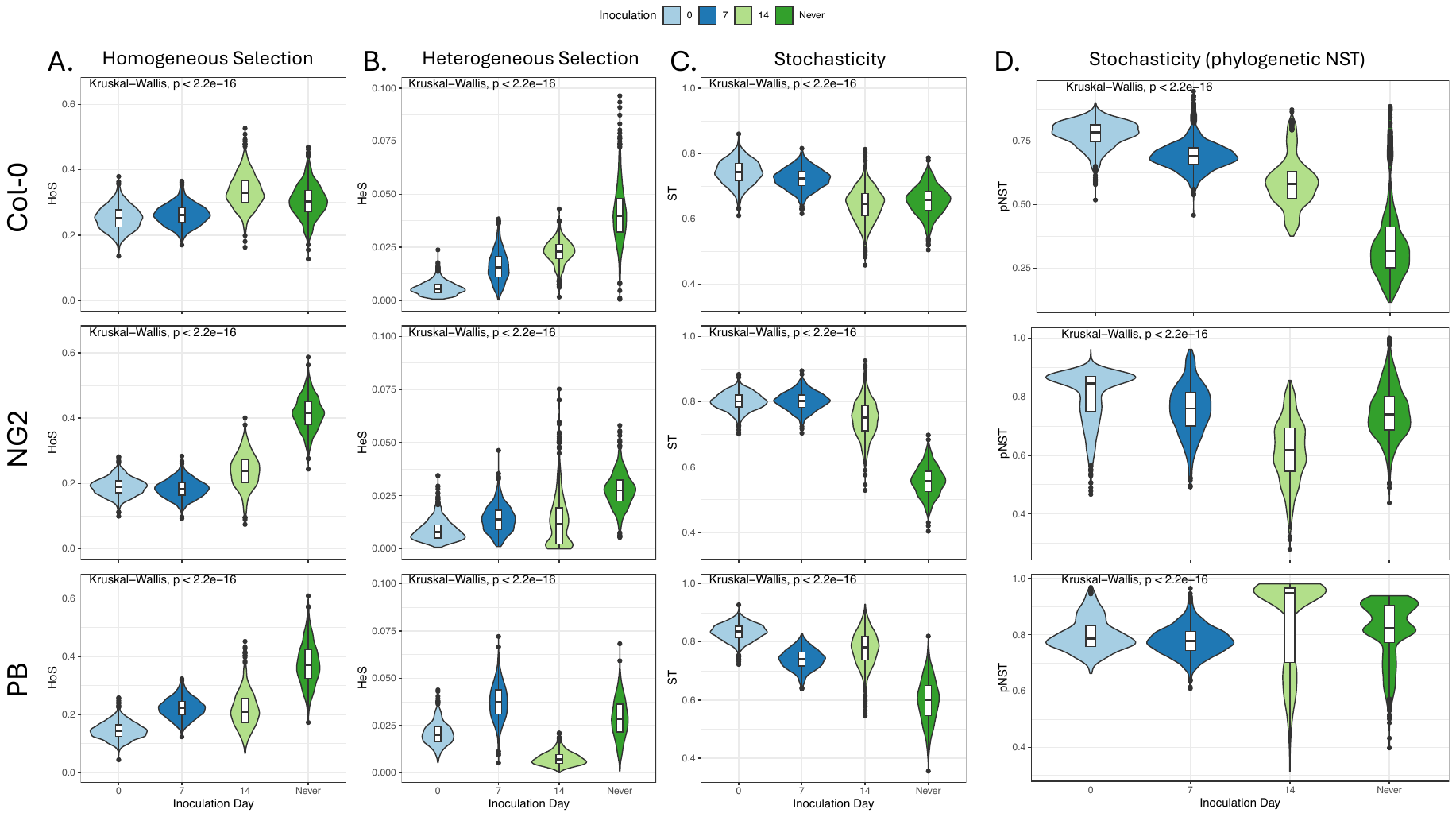

**Figure S10.** **Processes shaping bacterial colonization of leaves of the *A. thaliana* genotypes Col-0, NG2 and PB depend on both inoculation time and genotype.** A.–C. are based on the phylogenetic bin-based null model analysis in iCAMP. A.-B. The relative importance of deterministic processes (homogeneous and heterogeneous selection, respectively) and C. the relative importance of of stochastic processes (the sum of Dispersal limitation, Drift, and Homogenizing dispersal) in colonization of the leaves of plants inoculated at ID0, ID7, ID14 or never inoculated. D. The phylogenetic normalized stochasticity ratio (phylogenetic NST) is also provided as an independent measure of stochasticity for comparison. The top row is Col-0, the second row is NG-2 and the third row is PB.

**Figure S11**

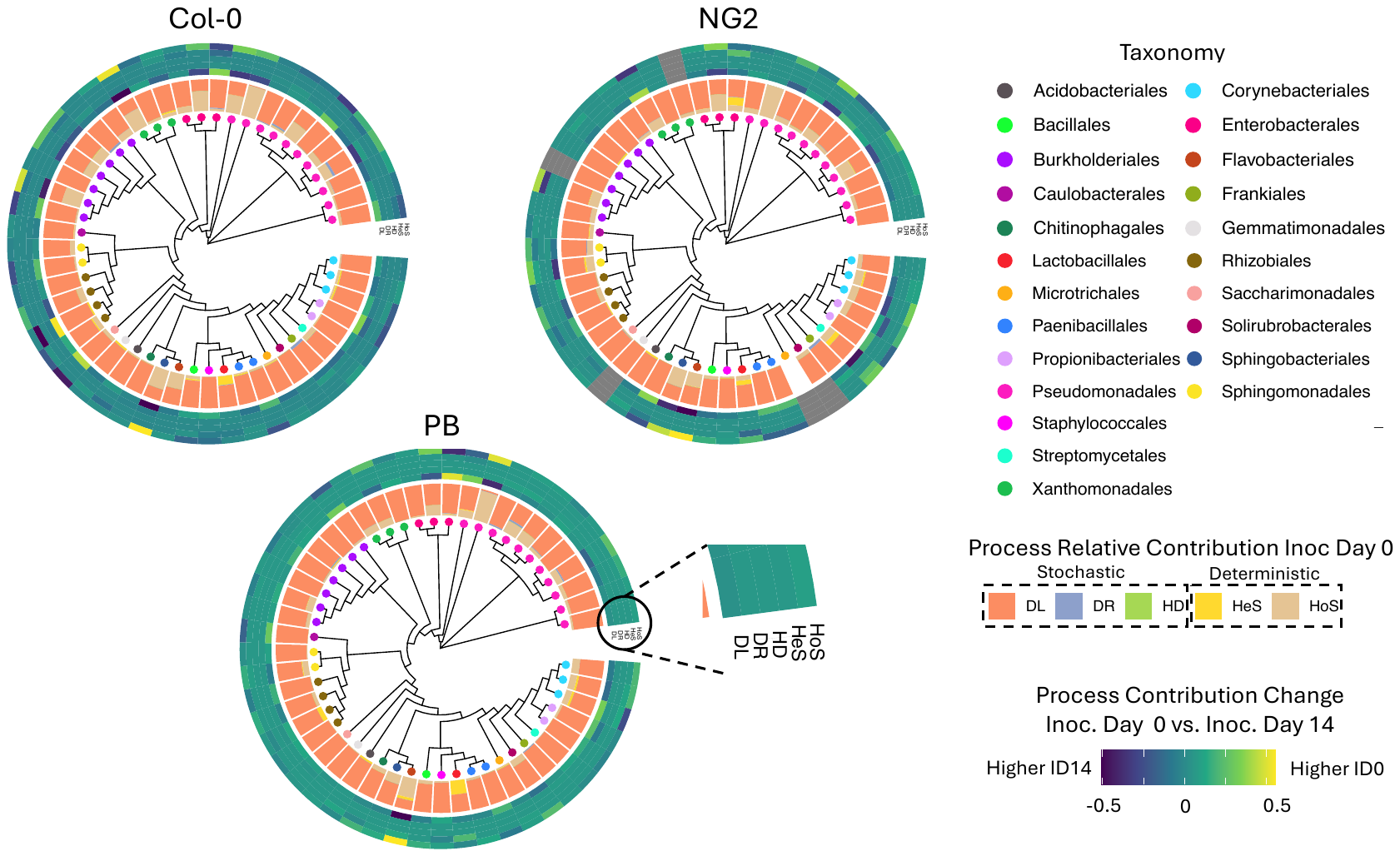

**Figure S11.** **Processes shaping bacterial colonization of leaves of the *A. thaliana* genotypes Col-0, NG2 and PB depend on both inoculation time and genotype.** Assembly processes at the level of taxonomic bins. The trees are all three identical and are annotated with, from inside to outside: Colored dots representing the majority taxonomy of each bin, A barchart showing for plants inoculated at ID0 the relative contribution of each community assembly process for each bin (DL: Dispersal limitation, DR: Drift, HD: Homogenizing dispersal, HeS: Heterogeneous selection, HoS: Homogeneous selection), and A heatmap showing the change in relative importance of each assembly process for each bin between plants inoculated at ID0 and those inoculated at ID14 (yellow is higher at ID0 dark blue is higher at ID 14). Figure 3B in the main text also shows labeled the taxa that are mentioned in the text that were selected deterministically at ID0 and stochastically at ID14 (homologous selection increased by at least 20% to reach >50% total contribution in D0).

**Figure S12**

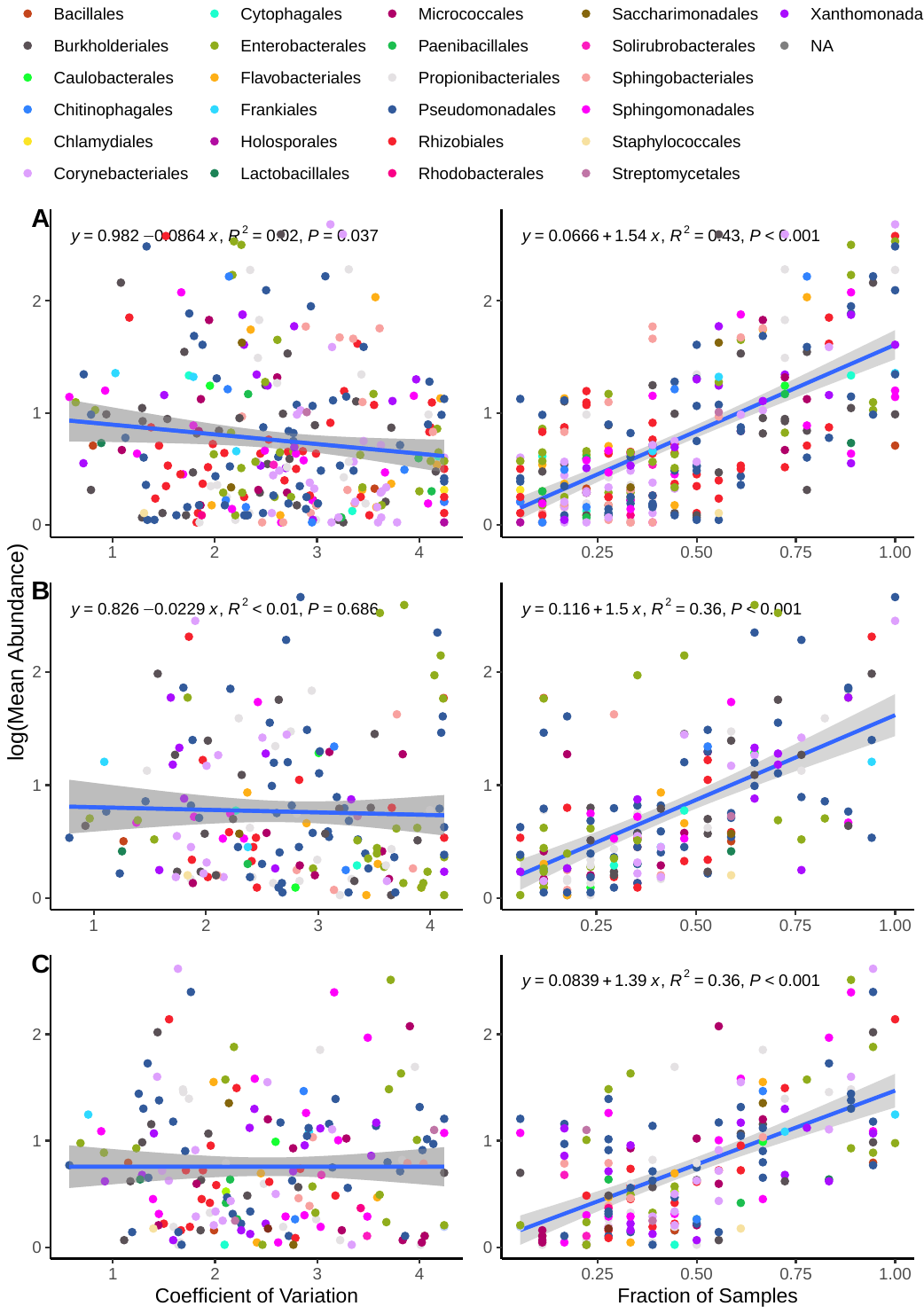

**Figure S12.** Correlation of the log mean abundance of ASVs to either the coefficient of variation (a measure of stochasticity of abundance) or the fraction of samples the taxa was detected in. Measurements are shown for ecotypes Col-0 (**A**), NG2 (**B**) and PB (**C**). The measures are calculated using data from leaf samples collected at SD 21, 28 and 35.

**Figure S13**

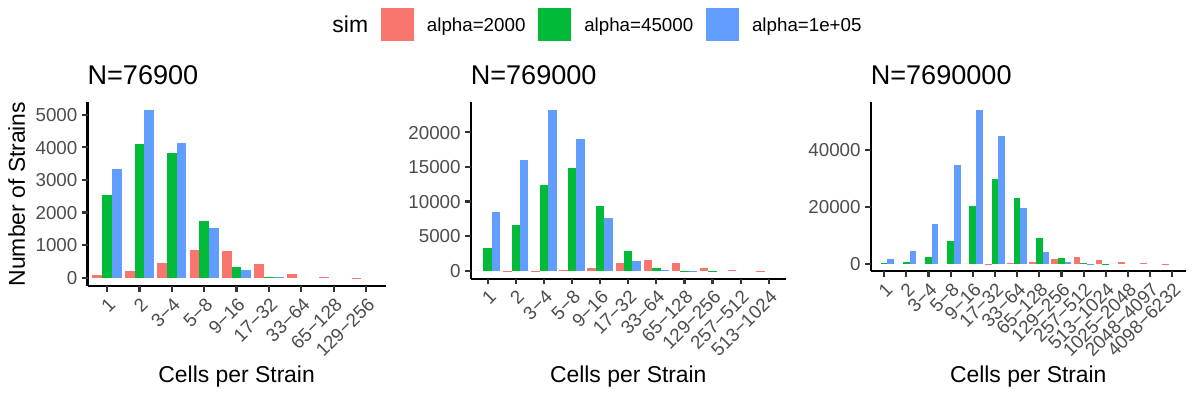

**Figure S13.** Preston’s Octave plots for the number of strains (y-axis) that fall within a range for a given number of cells per strain (x-axis). The three plots correspond to three orders of magnitude of N, the density of bacterial cells in soil (cells/mm^3^) and each plot is calculated for three levels of alpha, Fisher’s diversity index. The equations that the plots are based on can be found in the supplementary results.

**Figure S14**

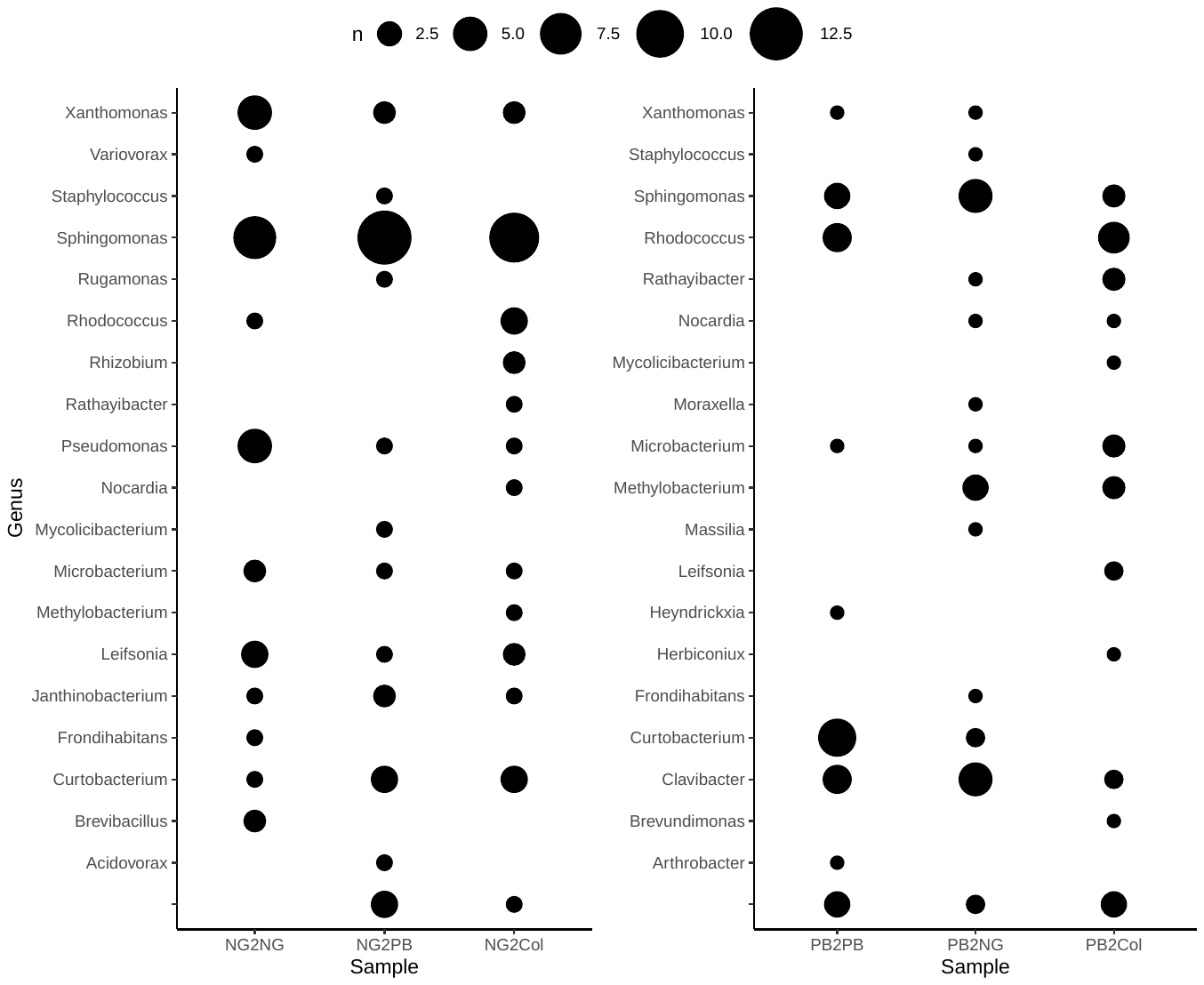

**Figure S14.** **Genus-level taxonomy and number of isolates of each genus recovered from *in-planta* isolation, where NG or PB-derived bacteria were inoculated onto NG, PB or Col-0 genotypes.** Associated statistics can be found in **Table S7**.

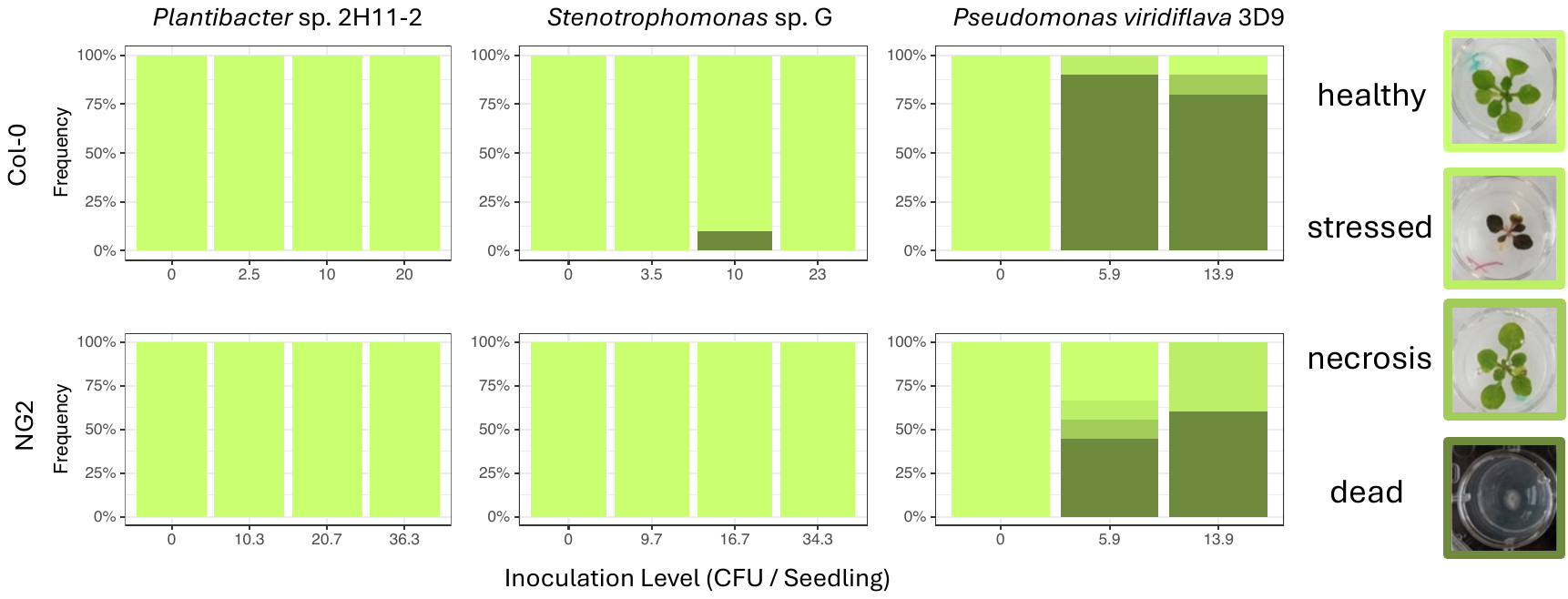

**Figure S15.** **Pseudomonas viridiflava 3D9 is an opportunistic pathogen of *A. thaliana*.** Germinating *A. thaliana* Col-0 or NG2 seedlings were inoculated with very low levels of three bacteria (CFU/seedling is indicated). After 14 days, plant phenotypes were recorded. healthy = no symptoms, stressed = purple/discolored or chlorotic leaves, necrotic = spots of dead tissue on at least one leaf, or dead. N=10.

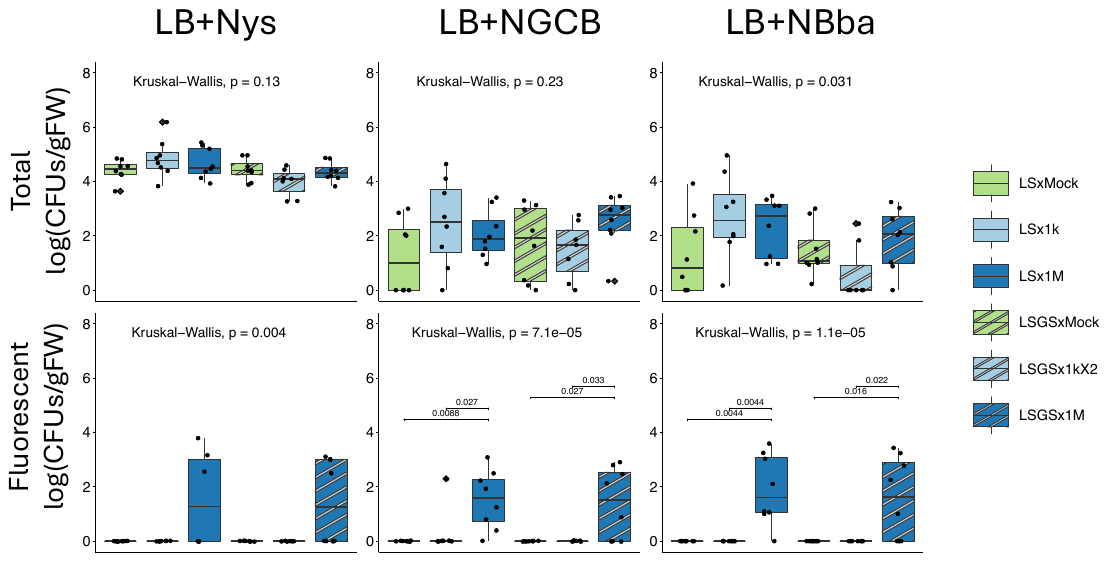

**Figure S16. *Pseudomonas viridiflava*** **3D9 transitions from soil to leaves based on fluorescent cell counts.** Seeds were germinated in standard laboratory soil (LS) or LS amended with an extract of a natural garden soil (LS+GS). The three treatments in each soil are no *Pseudomonas viridiflava* 3D9, low 3D9 (~1000 cells/cm^3^) or moderate 3D9 (~1x10^6^ cells/cm^3^). 14-d old plants were harvested and bacterial CFU counts were made on LB medium with nystatin (LB+N – total bacterial count), LB+N with gentamycin, chloramphenicol and bacitracin (LB+NGCB – selection against gram + bacteria) or LB+N with bacitracin and boric acid (LB+NBba – selection of Pseudomonads). The same plates were also observed for fluorescent colonies with a stereo microscope equipped with a GFP filter to estimate CFUs expressing mTurquoise2 (*Pseudomonas viridiflava* 3D9).

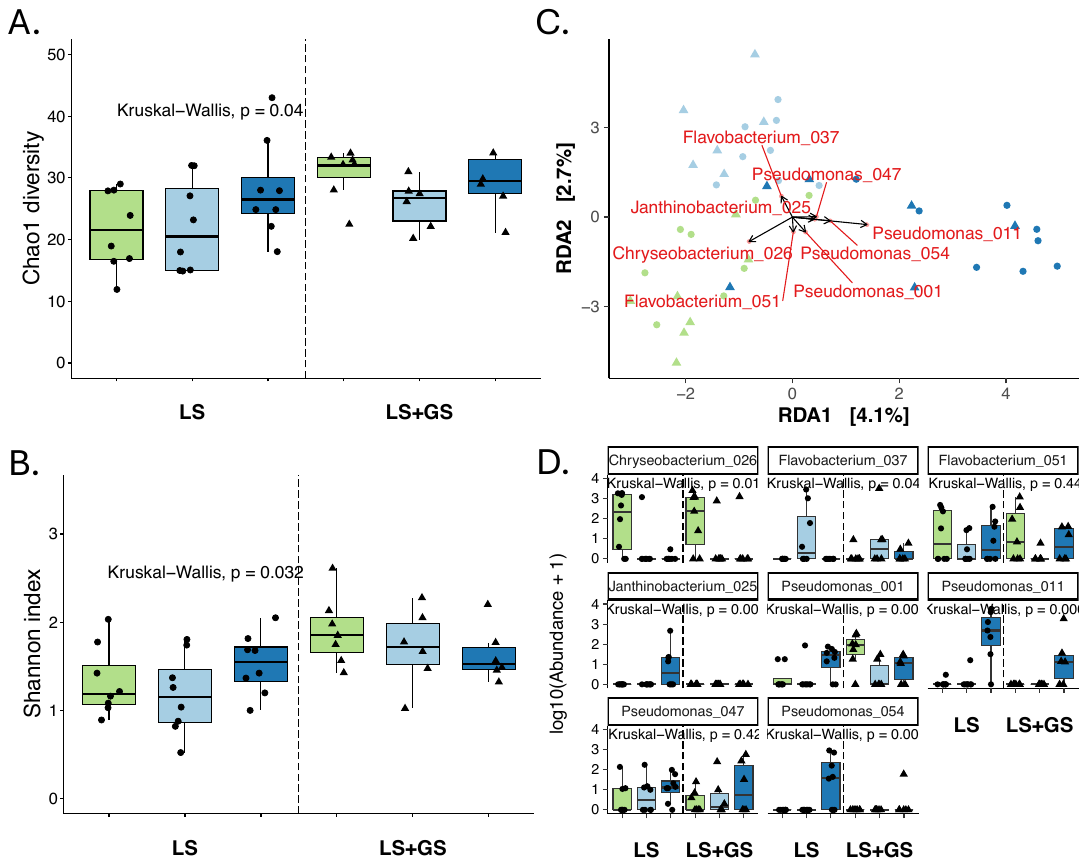

**Figure S17.** **Bacteria that deterministically transition from soil to mature leaves are influenced by *P. viridiflava* 3D9 soil amendment (ASV level analysis to complement Figure 5).** (A. and B.) Alpha diversity of bacteria in mature leaf tissues (14 days post germination) assessed by Chao1 metric (total estimated diversity) and Shannon index (diversity considering evenness of distribution of taxa). A Kruskal-Wallis test was performed on the data overall and when this was significant a Wilcoxon test was used to evaluate differences between groups. Significant p-values are shown. (C.) Principal components analysis of Aitchison distance between samples, constrained for treatment, showing biplot arrows for the top 1% of ASVs that correlate to the two axes. (D.) Boxplots showing the abundance of the bacteria identified using the biplot analysis in (C.) with p-value result of a Kruskal-Wallis test for significant differences between groups.

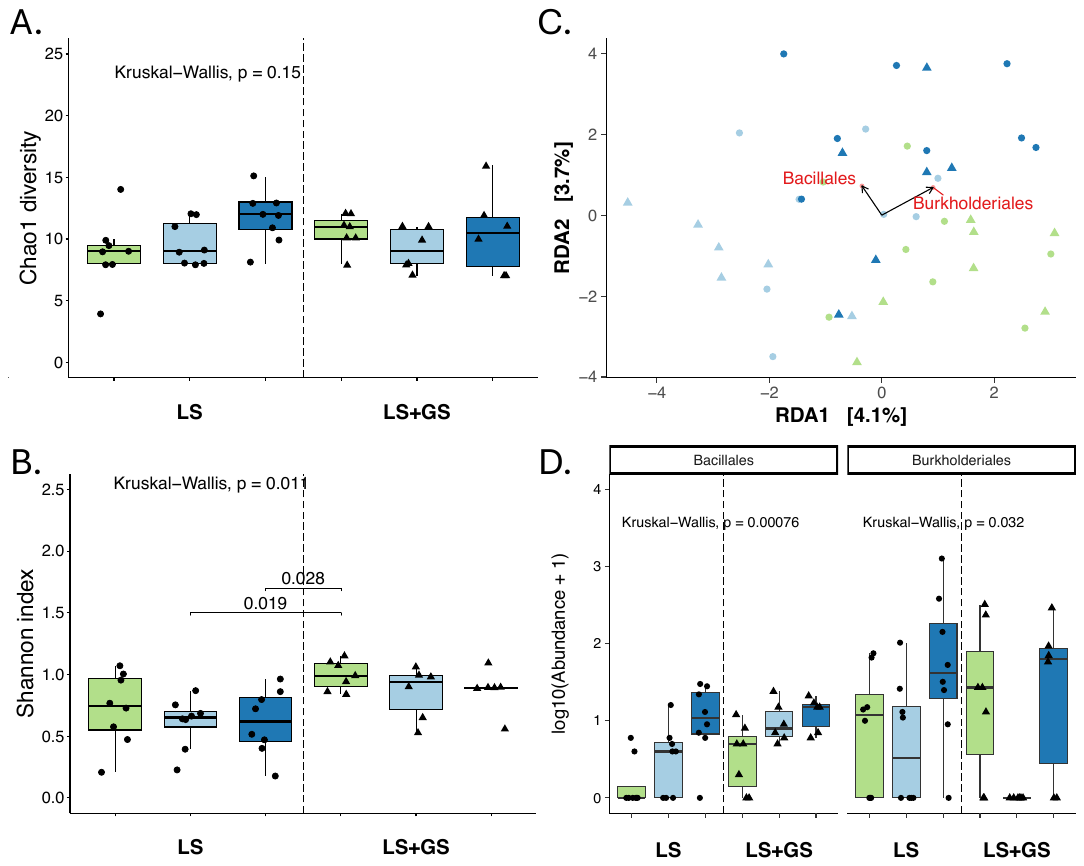

**Figure S18.** **Bacteria that deterministically transition from soil to mature leaves are influenced by *P. viridiflava* 3D9 soil amendment (Order-level analysis to complement Figure 5).** (A. and B.) Alpha diversity of bacteria in mature leaf tissues (14 days post germination) assessed by Chao1 metric (total estimated diversity) and Shannon index (diversity considering evenness of distribution of taxa). A Kruskal-Wallis test was performed on the data overall and when this was significant a Wilcoxon test was used to evaluate differences between groups. Significant p-values are shown. (C.) Principal components analysis of Aitchison distance between samples, constrained for treatment, showing biplot arrows for the top 3% of taxa that correlate to the two axes. (D.) Boxplots showing the abundance of the bacteria identified using the biplot analysis in (C.) with p-value result of a Kruskal-Wallis test for significant differences between groups.

**Table S1:** Significance of factors in describing the variation in bacterial communities between samples. Results are based on the Aitchison distance between samples collected at all time points at the level of ASVs. Analysis was carried out with PERMANOVA with 1000 permutations.

| **All Samples** | **Df** | **SumsOfSqs** | **MeanSqs** | **F.Model** | **R2** | **Pr(>F)** |
| --- | --- | --- | --- | --- | --- | --- |
| **Inoculation Day** | 3 | 26738.992 | 8912.99735 | 4.38676152 | **0.05721708** | **0.001** |
| **Sampling Day** | 3 | 16583.9481 | 5527.98269 | 2.72073925 | **0.03548694** | **0.001** |
| **Ecotype** | 2 | 11195.2208 | 5597.61039 | 2.75500832 | **0.02395595** | **0.001** |
| **Inoculation Day x Sampling Day** | 8 | 21582.5743 | 2697.82179 | 1.32780258 | **0.04618319** | **0.001** |
| **Inoculation Day x Ecotype** | 6 | 16935.0508 | 2822.50847 | 1.38917034 | **0.03623825** | **0.001** |
| **Sampling Day x Ecotype** | 6 | 13137.6584 | 2189.60974 | 1.07767291 | 0.02811245 | 0.084 |
| **Inoculation Day x Sampling Day x Ecotype** | 15 | 32001.2047 | 2133.41365 | 1.05001455 | 0.06847736 | 0.102 |
| **Residuals** | 162 | 329150.687 | 2031.79437 |  | 0.70432879 |  |
| **Total** | 205 | 467325.336 |  |  | 1 |  |
| **Col-0 Samples** |  |  |  |  |  |  |
| **Inoculation Day** | 3 | 16393.9947 | 5464.66488 | 3.00508769 | **0.11061315** | **0.001** |
| **Sampling Day** | 3 | 9549.28811 | 3183.09604 | 1.75042439 | **0.06443072** | **0.001** |
| **Inoculation Day x Sampling Day** | 8 | 16795.571 | 2099.44638 | 1.15451187 | 0.11332266 | 0.014 |
| **Residuals** | 58 | 105471.319 | 1818.47102 |  | 0.71163347 |  |
| **Total** | 72 | 148210.173 |  |  | 1 |  |
| **NG-2 Samples** |  |  |  |  |  |  |
| **Inoculation Day** | 3 | 11908.5664 | 3969.52214 | 2.21946062 | **0.10239978** | **0.001** |
| **Sampling Day** | 3 | 7769.06882 | 2589.68961 | 1.44796122 | **0.06680493** | **0.001** |
| **Inoculation Day x Sampling Day** | 8 | 16134.3728 | 2016.79661 | 1.12764219 | 0.13873678 | 0.013 |
| **Residuals** | 45 | 80482.8409 | 1788.50758 |  | 0.69205852 |  |
| **Total** | 59 | 116294.849 |  |  | 1 |  |
| **PB Samples** |  |  |  |  |  |  |
| **Inoculation Day** | 3 | 12382.7648 | 4127.58825 | 2.03726879 | **0.07812923** | **0.001** |
| **Sampling Day** | 3 | 10577.4528 | 3525.81762 | 1.74025066 | **0.06673859** | **0.001** |
| **Inoculation Day x Sampling Day** | 7 | 15994.2198 | 2284.88855 | 1.12776078 | 0.10091576 | 0.024 |
| **Residuals** | 59 | 119536.366 | 2026.0401 |  | 0.75421642 |  |
| **Total** | 72 | 158490.803 |  |  | 1 |  |

**Table S2:** Significance of factors in describing the variation in bacterial communities between samples. Results are based on the Aitchison distance between samples collected at all time points at the Order level. Analysis was carried out with PERMANOVA with 1000 permutations.

| **All Samples** | **Df** | **SumsOfSqs** | **MeanSqs** | **F.Model** | **R2** | **Pr(>F)** |
| --- | --- | --- | --- | --- | --- | --- |
| **Inoculation Day** | 3 | 2129.214 | 709.738 | 6.048 | **0.073** | **0.001** |
| **Sampling Day** | 3 | 1327.271 | 442.424 | 3.770 | **0.046** | **0.001** |
| **Ecotype** | 2 | 823.086 | 411.543 | 3.507 | **0.028** | **0.001** |
| **Inoculation Day x Sampling Day** | 8 | 1676.639 | 209.580 | 1.786 | **0.058** | **0.001** |
| **Inoculation Day x Ecotype** | 6 | 1324.624 | 220.771 | 1.881 | **0.045** | **0.001** |
| **Sampling Day x Ecotype** | 6 | 812.275 | 135.379 | 1.154 | 0.028 | 0.121 |
| **Inoculation Day x Sampling Day x Ecotype** | 15 | 2025.143 | 135.010 | 1.150 | 0.070 | 0.044 |
| **Residuals** | 162 | 19012.078 | 117.359 | NA | 0.653 | NA |
| **Total** | 205 | 29130.329 | NA | NA | 1.000 | NA |
| **Col-0 Samples** |  |  |  |  |  |  |
| **Inoculation Day** | 3 | 1765.993 | 588.664 | 5.413 | **0.178** | **0.001** |
| **Sampling Day** | 3 | 674.715 | 224.905 | 2.068 | **0.068** | **0.001** |
| **Inoculation Day x Sampling Day** | 8 | 1191.842 | 148.980 | 1.370 | **0.120** | **0.013** |
| **Residuals** | 58 | 6307.801 | 108.755 | NA | 0.635 | NA |
| **Total** | 72 | 9940.351 | NA | NA | 1.000 | NA |
| **NG-2 Samples** |  |  |  |  |  |  |
| **Inoculation Day** | 3 | 734.543 | 244.848 | 2.258 | **0.101** | **0.002** |
| **Sampling Day** | 3 | 659.753 | 219.918 | 2.028 | **0.090** | **0.001** |
| **Inoculation Day x Sampling Day** | 8 | 1016.835 | 127.104 | 1.172 | 0.139 | 0.104 |
| **Residuals** | 45 | 4879.014 | 108.423 | NA | 0.669 | NA |
| **Total** | 59 | 7290.145 | NA | NA | 1.000 | NA |
| **PB Samples** |  |  |  |  |  |  |
| **Inoculation Day** | 3 | 851.916 | 283.972 | 2.456 | **0.089** | **0.001** |
| **Sampling Day** | 3 | 743.900 | 247.967 | 2.145 | **0.077** | **0.001** |
| **Inoculation Day x Sampling Day** | 7 | 1196.352 | 170.907 | 1.478 | **0.124** | **0.001** |
| **Residuals** | 59 | 6821.658 | 115.621 | NA | 0.710 | NA |
| **Total** | 72 | 9613.825 | NA | NA | 1.000 | NA |

***Table S3:*** Significance of factors in describing the variation in bacterial communities between samples. Results are based on the Aitchison distance between samples collected at 21 DPI at the ASV level. Analysis was carried out with PERMANOVA with 1000 permutations.

| **All** | **Df** | **SumsOfSqs** | **MeanSqs** | **F.Model** | **R2** | **Pr(>F)** |
| --- | --- | --- | --- | --- | --- | --- |
| **Inoculation Day** | 2 | 10126.521 | 5063.260 | 2.380 | **0.096** | **0.001** |
| **Ecotype** | 2 | 6097.501 | 3048.750 | 1.433 | **0.058** | **0.001** |
| **Inoculation Day x Ecotype** | 4 | 10022.720 | 2505.680 | 1.178 | **0.095** | **0.009** |
| **Residuals** | 37 | 78723.642 | 2127.666 | NA | 0.750 | NA |
| **Total** | 45 | 104970.383 | NA | NA | 1.000 | NA |
| **Inoculation Day 0** |  |  |  |  |  |  |
| **Ecotype** | 2 | 3290.990 | 1645.495 | 1.204 | **0.147** | **0.021** |
| **Residuals** | 14 | 19131.147 | 1366.511 | NA | 0.853 | NA |
| **Total** | 16 | 22422.137 | NA | NA | 1.000 | NA |
| **Inoculation Day 7** |  |  |  |  |  |  |
| **Ecotype** | 2 | 5265.983 | 2632.991 | 1.201 | **0.146** | **0.009** |
| **Residuals** | 14 | 30700.531 | 2192.895 | NA | 0.854 | NA |
| **Total** | 16 | 35966.513 | NA | NA | 1.000 | NA |
| **Inoculation Day 14** |  |  |  |  |  |  |
| **Ecotype** | 2 | 2613.147 | 1306.573 | 1.552 | **0.256** | **0.005** |
| **Residuals** | 9 | 7574.759 | 841.640 | NA | 0.744 | NA |
| **Total** | 11 | 10187.906 | NA | NA | 1.000 | NA |

***Table S4:*** Significance of factors in describing the variation in bacterial communities between samples. Results are based on the Aitchison distance between samples collected at 21 DPI at the Order level. Analysis was carried out with PERMANOVA with 1000 permutations.

| **All** | **Df** | **SumsOfSqs** | **MeanSqs** | **F.Model** | **R2** | **Pr(>F)** |
| --- | --- | --- | --- | --- | --- | --- |
| **Inoculation Day** | 2 | 844.842 | 422.421 | 3.521 | **0.135** | **0.001** |
| **Ecotype** | 2 | 375.465 | 187.733 | 1.565 | **0.060** | **0.011** |
| **Inoculation Day x Ecotype** | 4 | 616.310 | 154.077 | 1.284 | **0.098** | **0.032** |
| **Residuals** | 37 | 4438.948 | 119.972 | NA | 0.707 | NA |
| **Total** | 45 | 6275.565 | NA | NA | 1.000 | NA |
| **Inoculation Day 0** |  |  |  |  |  |  |
| **Ecotype** | 2 | 226.675 | 113.337 | 1.387 | 0.165 | 0.066 |
| **Residuals** | 14 | 1144.305 | 81.736 | NA | 0.835 | NA |
| **Total** | 16 | 1370.979 | NA | NA | 1.000 | NA |
| **Inoculation Day 7** |  |  |  |  |  |  |
| **Ecotype** | 2 | 194.193 | 97.096 | 0.887 | 0.112 | 0.735 |
| **Residuals** | 14 | 1532.558 | 109.468 | NA | 0.888 | NA |
| **Total** | 16 | 1726.751 | NA | NA | 1.000 | NA |
| **Inoculation Day 14** |  |  |  |  |  |  |
| **Ecotype** | 2 | 266.004 | 133.002 | 1.711 | **0.276** | **0.010** |
| **Residuals** | 9 | 699.442 | 77.716 | NA | 0.724 | NA |
| **Total** | 11 | 965.446 | NA | NA | 1.000 | NA |

**Table S5**: Number of bacterial colonizers recovered by traditional medium-based isolation methods in comparison with our in-planta approach (2018) and comparison between cross-inoculations of leaf inocula onto different plant genotypes (2019).

| **Isolation method** | **Inoculum** | **Inoculation time point** | **n (total)** | **n (positive)** | **positive inocula [%]** |
| --- | --- | --- | --- | --- | --- |
| **NG2 🡪 R2A liquid (2018)** |  |  |  |  |  |
|  | 0-1 cells per plant (10^-4^ dilution) | - | 300 drops | 142 | 47.3 |
|  | 0-1 cells per plant (10^-5^ dilution) | - | 1200 drops | 66 | 5.8 |
| **NG2 🡪 R2A agar (2018)** |  |  |  |  |  |
|  | 0-1 cells per plant (10^-4^ dilution) | - | 300 drops | 153 | 51.0 |
|  | 0-1 cells per plant (10^-5^ dilution) | - | 1160 drops | 91 | 7.9 |
| ***In-planta* (2018)** |  |  |  |  |  |
| NG2 🡪 NG2 | 0-1 cells per plant (10^-4^ dilution) | 2-d old plant | 91 | 22 | 24.2 |
|  | 0-1 cells per plant (10^-5^ dilution) | 2-d old plant | 88 | 0 | 0.0 |
|  | 0-1 cells per plant (10^-4^ dilution) | 6-d old plant | 65 | 15 | 23.1 |
|  | 0-1 cells per plant (10^-5^ dilution) | 6-d old plant | 88 | 3 | 3.4 |
| ***In-planta* (2019)** |  |  |  |  |  |
| NG2 🡪 NG2 | 3.5 cells/plant | 3-d old plant | 120 | 39 | 32.5 |
| NG2 🡪 PB | 3.5 cells/plant | 3-d old plant | 120 | 30 | 25.0 |
| NG2 🡪 Col-0 | 3.5 cells/plant | 3-d old plant | 120 | 66 | 55.0 |
| PB 🡪 PB | 1.45 cells/plant | 3-d old plant | 120 | 33 | 27.5 |
| PB 🡪 NG2 | 1.45 cells/plant | 3-d old plant | 120 | 67 | 55.8 |
| PB 🡪 Col-0 | 1.45 cells/plant | 3-d old plant | 120 | 41 | 34.2 |

***Table S6****:Counts and p-values for the likelihood that the number of times a genus was observed in-planta is the same as the expected number of observations based on counts in R2A medium. P-values are based on a Χ^2^ test comparing the expected number of observations (calculated based on the frequency observed in R2A medium) vs. the actual number of observations.*

|  | **in-planta** | | |  | **R2A** | | |  | **in-planta exp. based on R2A freq** | | | |
| --- | --- | --- | --- | --- | --- | --- | --- | --- | --- | --- | --- | --- |
| **10^-4^ dilution** | Count | Total Inocs | Freq |  | Count | Total Inocs | Freq |  | Expected | Total |  | ChiSq.test  p-vallue |
| Pseudomonas | 13 | 156 | 0.08 |  | 18 | 600 | 0.03 |  | 4.68 | 156 |  | 1.2E-4 |
| Xanthomonas | 17 | 156 | 0.11 |  | 6 | 600 | 0.01 |  | 1.56 | 156 |  | 4.2E-35 |
| Rhizobiaceae | 2 | 156 | 0.01 |  | 5 | 600 | 0.008 |  | 1.3 | 156 |  | 0.54 |
| **10^-5^ dilution** | Count | Total Inocs | Freq |  | Count | Total Inocs | Freq |  | Expected | Total |  | ChiSq.test |
| Pseudomonas | 1 | 176 | 0.006 |  | 34 | 2360 | 0.01 |  | 2.54 | 176 |  | 0.33 |
| Xanthomonas | 1 | 176 | 0.006 |  | 12 | 2360 | 0.005 |  | 0.89 | 176 |  | 0.91 |
| Rhizobiaceae | 0 | 176 | 0 |  | 20 | 2360 | 0.008 |  | 1.49 | 176 |  | 0.22 |

***Table S7****: fdr-adjusted p-values for genus-level taxonomy and number of isolates of each genus recovered from in-planta isolation, where NG or PB-derived bacteria were inoculated onto NG, PB or Col-0 genotypes. P-values are based on a X^2^ test to determine whether a genus significantly more often originated from PB or NG (origin) or was recovered significantly more often in PB, Col or NG. p-values are only shown for genera where there were at least two observations in the tested conditions.*

| **Genus** | **Origin**  **(PB vs NG)** | **Recipient**  **(PB --> PB, Col or NG)** | | **Recipient**  **(NG --> PB, Col or NG)** |
| --- | --- | --- | --- | --- |
| Acidovorax |  |  | |  |
| Arthrobacter | | |  | |
| Brevibacillus | | |  | |
| Brevundimonas | | |  | |
| Clavibacter | **1.2E-10** | 3.3E-01 | |  |
| Curtobacterium | 5.6E-01 | **8.2E-02** | | 6.7E-01 |
| Frondihabitans | 1.0E+00 |  | |  |
| Herbiconiux |  |  | |  |
| Heyndrickxia | | |  | |
| Janthinobacterium | **1.2E-17** |  | | 7.7E-01 |
| Leifsonia | 3.8E-01 |  | | 6.7E-01 |
| Massilia |  |  | |  |
| Methylobacterium | 1.1E-01 | 1.0E+00 | |  |
| Microbacterium | 1.0E+00 | 6.1E-01 | | 7.7E-01 |
| Moraxella |  |  | |  |
| Mycolicibacterium | 1.0E+00 |  | |  |
| Nocardia | 1.0E+00 |  | |  |
| Pseudomonas | **2.4E-15** |  | | 1.4E-01 |
| Rathayibacter | 4.7E-01 | 8.0E-01 | |  |
| Rhizobium |  |  | |  |
| Rhodococcus | 1.6E-01 | 1.0E+00 | | 6.7E-01 |
| Rugamonas |  |  | |  |
| Sphingomonas | **6.6E-03** | 5.1E-01 | | 5.7E-01 |
| Staphylococcus | 1.0E+00 |  | |  |
| Variovorax |  |  | |  |
| Xanthomonas | 1.0E-01 |  | | 5.1E-01 |

***Table S8****: P-values for the likelihood that the number of times a genus was observed co-colonizing with another taxa was by chance alone. P-values are based on a Χ^2^ test comparing the expected number of observations (87.5% of total observations alone and 12.5% of total observations with a co-colonizer – based on the totals in the experiment) vs. the actual number of times the genus was observed alone and with a co-colonizer. Genera in bold are those who were mentioned in the text because they were found more often paired with another bacterium than alone. Orange colored genera denote the Burkholderiales.*

| **Genus** | **Observed**  **Paired** | **Observed**  **Alone** | **Observed**  **Total** | **Expected**  **Paired** | **Expected**  **Alone** | ***Χ*^2^ test**  **p-value** |
| --- | --- | --- | --- | --- | --- | --- |
| Sphingomonas | 10 | 36 | 46 | 5.75 | 40.25 | 0.05812607 |
| **Methylobacterium** | **3** | **5** | **8** | **1** | **7** | **0.03250944** |
| Rathayibacter | 2 | 3 | 5 | 0.625 | 4.375 | 0.06297905 |
| Microbacterium | 3 | 6 | 9 | 1.125 | 7.875 | 0.05878172 |
| **Janthinobacterium** | **4** | **0** | **4** | **0.5** | **3.5** | **1.2132E-07** |
| **Variovorax** | **1** | **0** | **1** | **0.125** | **0.875** | **0.00815097** |
| **Acidovorax** | **1** | **0** | **1** | **0.125** | **0.875** | **0.00815097** |
| **Rugamonas** | **1** | **0** | **1** | **0.125** | **0.875** | **0.00815097** |
| Xanthomonas | 2 | 9 | 11 | 1.375 | 9.625 | 0.5688114 |
| Curtobacterium | 1 | 17 | 18 | 2.25 | 15.75 | 0.37299848 |
| Rhodococcus | 1 | 14 | 15 | 1.875 | 13.125 | 0.49452467 |
| Nocardia | 1 | 2 | 3 | 0.375 | 2.625 | 0.27523352 |
| **Clavibacter** | **5** | **9** | **14** | **1.75** | **12.25** | **0.00862942** |
| Staphylococcus | 1 | 1 | 2 | 0.25 | 1.75 | 0.10880943 |
| Leifsonia | 1 | 7 | 8 | 1 | 7 | 1 |
| Rhizobium | 1 | 1 | 2 | 0.25 | 1.75 | 0.10880943 |
| **All Burkholderiales** | **7** | **0** | **7** | **0.875** | **6.125** | **2.5596E-12** |

**Supplementary Methods**

**Flow Pot Experiments**

**Preparations of flow pots**

*Potting soil.* The potting soil consisted of 1 part Floraton 3 Floragard Potting soil, 0.5 parts perlite (Perligran Premium Perlite), 0.25 parts sandbox sand and 7.1g fertilizer per liter of soil (Substral Osmocote 6-month garden flower fertilizer). All ingredients were mixed dry and distributed in 500g aliquots into Sun bags (Merck), autoclaved (sterilizing for 20 min at 121°C and drying for 10 min at 121°C) and allowed to cool down over night.

*Flow Pot assembly*. The axenic plant growth system (Flow Pots) is adapted from Kremer et al. (1). All individual parts were autoclaved before Flow Pot assembly (sterilizing for 20 min at 121°C and drying for 10 min at 121°C). In short, 50mL syringes were cut at the 20mL mark. 20 glass beads (diameter 3mm; Merck) are added to each Flow Pot followed by ~10g of double-autoclaved potting soil. A 7x7 centimeter mesh was put over the soil and attached with a heat-stable cable tie (Kabelbinder Discount). Nine prepared Flow Pots were placed into each microbox (SacO2) by attaching them in the holes of two empty tip racks. A 20cm tube (inner diameter 4mm, outer diameter 6mm, wall thickness 1mm) was connected to the luer lock on the bottom of the Flow Pot and the other end was guided up through one hole of the tip stand so that inoculation can be performed later from above. Assembled boxes were autoclaved (sterilizing for 20 min at 121°C and drying for 10 min at 121°C) and allowed to cool down to room temperature before the start of the experiment.

*Soil inoculum.* Garden soil was collected (22.07.2019) by taking soil >6cm below ground, drying at room temperature for 1 week, then sieving to remove larger debris. The prepared soil was stored at 4°C. To generate an inoculum slurry, 50g of this soil was mixed with 950g of water for 20min at 250rpm. It was left to rest for one hour to let the solid material settle down. The supernatant was then transferred into a new bottle. This inoculum was always prepared from the dried, sieved soil one day before inoculation. For the control the soil inoculum was prepared and autoclaved 2 times (sterilizing for 25 min at 121°C) one day before the inoculation and allowed to cool down in between the autoclaving runs and over night before it was used. Both inoculums were mixed in equal parts (1:1) with full MS basal salt mix (Murashige and Skoog basal salt mix) before applying it to the plant soil. For the inoculation performed at later timepoint using the airbrush systems, 0.02% of Silwet-77 was added to the Soil-MS mix to reduce surface tension.

*Seed vernalization.* Three days before the start of the experiment, seeds of the plant ecotypes NG2 (NASC N2110865), PB (until available via NASC it is available form the authors upon request.) and Col-0 were sterilized by soaking in 70% ethanol for 1 min, followed by 2% bleach for 1 min, followed by three washes with sterile water. Finally, seeds were left in 0.1% agar in the dark at 4°C for vernalization for three days. NG2 and PB are isolates of wild populations that have been selfed to generate homozygous lines before they were used in this experiment (2).

***In-planta* isolation of leaf bacteria experiment**

*Assessing the appropriate dilution of the leaf microbe extract inoculum.* In both years, we serially diluted the glycerol stock in PBS/S to obtain dilutions from 10^-1^ to 10^-6^. For each dilution, 5 µL were inoculated into 48 wells of a 96-well plate containing 200 µL R2A medium per well and positive wells were counted after three days. For experimental inoculations, we identified near-extinction dilutions (where most wells did not have any growth). Based on CFU counts, final inoculations in 2018 contained 0-1 cell/seedling (NG2, 2018). Because the frequency of colonized plants was low, in 2019 we used a lower dilution to 3.5 and 1.45 cells/seedling (NG2 and PB, 2019, respectively) which still resulted in colonization mostly by one strain per plant (but not only - see details in results).

*Identifying efficient early bacterial colonizers in NG2 (2018 experiments).*

Two weeks after inoculation, we checked via PCR how many plants were colonized by one bacterial taxon to identify efficient early colonizers, that can colonize a plant from one bacterial cell. We first extracted bacterial DNA from plant material by bead beating the leaves in 500 µL SDS extraction buffer (0.5% filter-sterilized SDS, 50 mM Tris-HCl pH = 8.0, 200 mM NaCl, 2 mM EDTA, prepared with NFW) using ~0.2 g (0.25-0.5 mm diameter) and ~0.4 g (1.25-1.55 mm) glass beads (Carl Roth) for 45 s at 2,400 rpm. We then purified DNA by a magnetic bead cleanup using home-made Sera-Mag purification beads (3) and re-suspended in 40 μL 10 mM Tris-HCl pH = 8.0. We amplified the V5-V7 region of the 16S rRNA gene using 799F/1391R primers (Suppl. Tab. S2). One 50 µL PCR reaction contained 1x Buffer B, 0.2 mM dNTPs, 2 mM MgCl2, 0.5 µL 500 U Taq Polymerase (Biodeal), 0.2 µM of each primer, 34.5 µL nuclease free water, 2 µL template DNA and was amplified with 3 min initial denaturation at 94 °C, followed by 35 cycles of denaturation at 94 °C for 1 min, annealing at 55 °C for 1 min, elongation at 72 °C for 30 s, and a final elongation at 72 °C for 10 min. The products were Sanger sequenced (Eurofins Genomics) with 1391R primers or a modified version of 1391R that avoids chloroplast 16S rRNA genes (Suppl. Tab. S2).

*Isolation and identification of colonizers in cross-inoculation experiments (2019 experiments).*

To both identify colonizers and generate a culture collection for later experiments, we isolated bacteria from leaves collected two weeks after inoculation and sequenced their 16S rRNA genes. In short, we added two sterilized metal beads (3 mm) to each well of 96-well plates containing the frozen leaf tissue from all six cross-inoculated treatments. The leaves were crushed by manual shaking for 5 min. We then added 100 μL PBS to each sample and prepared a 10-fold dilution series up to 10-3. 5 μL of each sample of all dilutions were plated on R2A and incubated for up to six days at 30 °C. Per treatment, we picked 30 random isolates. To identify them we extracted DNA from overnight cultures in R2A broth using 600 μL SDS extraction buffer (see above) per sample. ~0.2 g glass beads (0.25-0.5 mm) were added to each sample, incubated for 10 min at 37 °C, and bead-beat for 30 s at 13,400 rpm. Cell debris was centrifuged for 5 min at 20,000 x g and the supernatant was recovered. It was acidified with 1/3 vol. 5 M KOAc, cleaned-up with 1.5x Sera-Mag beads and re-suspended in 40 μL 10 mM Tris-HCl pH = 8.0. For identification, we amplified the V1-V9 hypervariable regions of the 16S rRNA genes with 8F/1492R primers (Suppl. Tab. S2) and Sanger sequenced (Eurofins Genomics) with the 1492R primer, repeating with 8F primers if necessary.

*Analysis of Sanger sequencing data.*

The resulting sequences were blasted against the NCBI 16S rRNA database. We used ten results for each sequence to assign taxonomy using the last-common-ancestor algorithm in MEGAN6 Community Edition 6.19.8 (4). Default settings were used except “Percent to Cover” was changed to 70% and “Min Percent Identity” to 97% to assign most isolates to the genus level. To check for significant differences in numbers of specific taxa isolated between inocula and recipient plant genotypes and to check for taxa that were isolated with a partner more often than expected, chi-square tests were performed in R (**Tables S6-S8**).

**P. viridiflava 3D9-141 soil amendment experiment**

*Media for CFU counting of total and 3D9-141 load*

Different media were used to determine CFU counts representing either total bacterial load or 3D9-141 bacterial loads (fluorescent colonies). To count total CFUs we used a media that restricts fungal growth (LB+N) containing 25 g/L Luria/Miller broth, 50 mg/L nystatin and 15g/L agar. To select for antibiotic resistance encoded in the cassette inserted in the 3D9-141 genome and against gram positive bacteria, LB+N was supplemented (LB+NGCB) with 15mg/L gentamycin, 15 mg/L chloramphenicol and 100mg/L bacitracin. To select for Pseudomonads (SP+NBba) according to Inoue et. al (5), we also used a Sucrose-Peptone medium with containing 20 g/L sucrose, 5 g/L peptone, 0.2 g/L MgSO4 7H2O, 1 g/L KH2PO4, 50 mg/L nystatin, 100 mg/L bacitracin, 10 mg/L ampicillin, 150 g/L boric acid and 15 g/L agar.

*qPCR estimation of 3D9-141 abundance in leaf tissues*

*mTurquoise2* should be present in only one copy in the 3D9-141 genome, making it a good proxy for the number of 3D9-141. A primer pair targeting *mTurquoise2* (mT2 fwd: AACGGCATCAAGGCCAACTTC, mT2 rev: ATGTGATCGCGCTTCTCGTTG, expected product size: 179 bp) was used in a qPCR carried out in a 96-well plate using KCQ S00 SYBR Green readymix (Merck Sigma-Aldrich). One microliter of sample gDNA was used as template in technical duplicates. A nuclease-free water template served as the no-template-control (NTC), while gDNA of axenically-grown *A. thaliana* NG2 was used as a negative control (NC). 3D9-141 gDNA was used as a positive control (PC). NTC, NC, and PC were added in triplicates. The amplification program included a denaturation step at 95^o^C for 30s, followed by 40 cycles of fluorescence reading, denaturation (95^o^C, 5s) and hybridization/elongation (63^o^C, 30s).

To convert Ct values to gene copies, isolated pMRE-Tn7-141 plasmid was linearized completely using high-fidelity EcoRI (NEB). Briefly, in a 200 µL digestions reaction, 1 µg of pMRE-Tn7-141 plasmid DNA was incubated in rCutSmart buffer with EcoRI-HF (20 u/µL) at 37 °C for 15 min. The results of this digestion were confirmed using gel electrophoresis. Thereafter, the qPCR was performed on a 1:8 dilution series of the linearized plasmid (from 32 to 0.0078 ng/µL) in triplicates. Using this standard curve, and the known pMRE-Tn7-141 copies, the mTurquoise2 copies of each sample were calculated.

The same samples were amplified in a qPCR using *A. thaliana* *EF1-α* primers (6) to normalize copy numbers to the host. This time, 3D9-141 gDNA served as a negative control, and the NG2 gDNA served as a positive control. The protocol was the same except the hybridization temperature was 62 °C. Further, a 1:8 dilution series of NG2 gDNA (from 679 to 0.02 ng/µL) was used to estimate the number of *EF1-α* gene copies in each sample.

**DNA extraction from leaf tissues.**

For all experiments, leaf samples frozen at -80°C were immediately bead-beaten (2 or 3x 30 s at 1,400 strokes/minute) in a BioSpec mini bead beater 96. The samples were briefly centrifuged and 200uL of CTAB extraction buffer (100mM TRIS pH 8.0, 20mM EDTA pH 8.0, 1.5 M NaCl, 2% cetylimethylammoniumbromid, 1%, polyvinylpyrrolidone (MW 40,000) in nuclease-free water) was added and mixed with the plant material by inverting the tubes a few times. Tubes were centrifuged and allowed to incubate at 37°C for 10 min.

For the samples from the inoculation time experiment, the tubes were centrifuged at high-speed (20.000g) for 5 minutes. The supernatant was recovered (about 200uL) into a round bottom deep-well plate. Then, the DNA was precipitated by mixing in 500 uL of ice-cold 100% ethanol and 20 uL 3M sodium acetate and centrifuging at maximum speed (2204g) at 4°C for 30 minutes. The supernatant was removed, and the pellets were washed twice with 200uL of 70% ethanol. The pellets were allowed to air dry for 15min and the DNA was re-hydrated in 100uL of 1x Tris-HCl (10mM). Next, the DNA was cleaned with in-house prepared Speed Beads (7). Samples were incubated with beads for 5 minutes at room temperature and were then placed on a magnetic stand until all the beads were drawn to the magnet and the supernatant was completely clear. The supernatant was removed, and samples were washed two times with 200uL of 70 % ethanol and then air dried for about 5 minutes. 40uL of Tris-HCl (10mM, pH 8) was added and samples were eluted by shaking at room temperature at 180 rpm for ~1 hour. After that samples were again placed on a magnetic stand and the supernatant was recovered into a new 96-well plate, which was frozen at -20°C until further use.

For samples from the 3D9-141 experiment, the supernatant from the CTAB extraction was mixed with 200uL of ice-cold phenol-chloroform-isoamylalcohol (25:24:1), centrifuged at top speed for 5 minutes, then the supernatant was precipitated overnight at -20C in an equal volume of isopropanol. After removing the isopropanol, the pellet was washed twice with 80% ethanol, air dried and resuspended in Tris-HCl (10mM, pH 8).

**16S rRNA gene amplicon sequencing library preparation**

*First amplification step*

Samples from 4 96-well plates were amplified in one target region (16S) in parallel on the same 384-well plate. First, 40uL reactions were prepared with 0.8 μL Kapa taq DNA polymerase (KAPA Biosystems), 1X Kapa GC Buffer, 0.08 μM of each forward and reverse primer, 0,25 μM of each blocking oligonucleotide, 0.3 mM dNTP and 26.4 μL nuclease free water. The primers were the same as those used in Unger et al. (2).The mixes were divided into four 9uL reactions with a 96-well pipetter (Platemaster, Gilson) on to a 384-well plate for independent amplification. To each of the master mixes 1uL of a 1:4 diluted genomic DNA sample template was added. These were run on a thermocycler block at 95 °C for 3 min, followed by 5 cycles of 98 °C for 20 sec, 55 °C for 1 min, 72 °C for 2 min with a final elongation at 72 °C for 2 min.

*Second amplification addition of indices and sequencing adapters*

The four replicate reactions were transferred into one 96-well plate. 10uL of the combined replicate reactions were cleaned enzymatically using Antarctic phosphatase and Exonuclease I (New England Biolabs, Inc)) to degrade leftover primers and inactivate nucleotides (0.5 μL each enzyme with 1.22 μL Antarctic phosphatase buffer at 37 °C for 30 minutes followed by 80 °C for 15 min). 5uL of the products of the enzymatic cleanup reaction were then used as template in a second, 20 μL PCR reaction with an identical setup as in the first step and for 20 cycles. The only difference was the use of concatenated barcoded primers that each had a 12-bp index and sequencing adapters.

*Library normalization*

Final products were brought to a volume of 50uL by adding 30uL of Tris-HCl (10mM, pH 8). They were then purified and normalized by using 0.5x volume home-made Sera-Mag purification beads and eluted in 40 μL TRIS-HCl. This ratio of beads to library also removes primer dimers. To 50uL PCR product 25uL SeraMag Speed Beads were added, mixed well and allowed to rest for 15 minutes. They were placed on a magnetic stand for 15 minutes to make sure that the beads are completely pulled down. The supernatant was discarded, and each sample was washed twice by carefully dripping 80% ethanol onto the well and carefully removing it. After a brief drying step 40uL of 10mM TRIS buffer was added and allowed to elute for 30 minutes while shaking at 900rpm on a Thermoshaker (VWR Thermal Shake lite). 1uL of the recovered DNA was added to a new 96-well plate and mixed with 9uL TE-buffer. 10uL of 200x diluted Picogreen (Thermo Fisher Scientific, Inc.). Fluorescence was measured on a qPCR machine (qTOWER3, Analytik Jena). The well with the highest fluorescence was used to create a standard curve (10x, 30x, 70x, 100x, 300x) based on which the relative concentrations of the individual samples was calculated. Samples were combined in similar ranges into sub-pools which were combined after a new fluorescence reading into one final pool to get equimolar libraries. Pools were again cleaned with 0.5x volume of speed beads to remove remaining primer dimers and eluted in 3.16x less volume (e.g., to a pool of 500uL 250uL of speed beads were added and eluted in 158uL). This step was repeated with a bead volume of 1.5x of the pool until one final pool with a volume of 100uL to 500uL was reached.

*Quantification and Sequencing of the pools.*

The exact length of the amplicons from each pool was analyzed on a 2100 Bioanalyzer Instrument and the library was quantified on a Qubit (Thermo Fisher Scientific, Inc.). The final library was denatured and then loaded onto a MiSeq lane spiked with 10% PhiX genomic DNA to ensure high enough sequence diversity. Sequencing was performed for 600 cycles to recover 300 bp of information in the forward and reverse directions with standard Illumina sequencing primers.

**Supplementary Results**

*Seeds germinating in natural soil will be exposed to limited cells of any given bacterial strain*

The success and effects of colonization of germinating seeds by a given bacterium in the soil will presumably depend on the inoculum size (in other words, the amount of the bacterium in the vicinity of the germinating seed). Thus, to design experiments that manipulate early colonization, we needed a realistic idea of inoculum sizes. Therefore, we applied a simple model to estimate the distribution of bacterial strains in the zone of influence (Z) of a germinating *A. thaliana* seed. First, we estimated the density of strains in soil (S) based on a large range of estimates covering several orders of magnitude for the density of bacterial cells in soil (N) and for their diversity (alpha) based on the equation for Fisher’s diversity index (Eq. 1).

$S= \alpha ln(1+\frac{N}{\alpha})$,

**(Eq. 1)**

where:

$S$= Density of bacterial strains in soil

$N$= Density of bacterial cells in soil

$\alpha$= Fisher’s diversity Index

Since strains are unlikely to be evenly distributed in soil, we then modeled a log-normal distribution using the estimates of N and S. In short, the results show that the exact distribution depends on alpha and N, but most bacterial strains in the zone of influence of a seed are likely to only be present in low abundance with <50 cells (**Figure S13**). Only in soil with extremely low alpha and very large N do abundances approach higher values, but even so, few strains reach more than about 500 cells. Because the total number of cells (N) and the number of species (S) scales with the seed zone of influence for small Z, these results do not depend on the size of the zone of influence of the seed. Since many leaf-associated bacterial strains in soil are likely to be among the relatively low-abundance soil members, we reason that most should have the capacity to reach and colonize leaves from just a few cells. Even though this is a drastic simplification of complex soil matrices and each seed in nature will likely have a unique microbial environment due to heterogeneity, it still provides a guideline to understand the lower limits of bacterial strain abundance so that we can investigate dynamics in this range.

*In-planta bacterial isolation and detection with PCR*

An extinction-diluted inoculum derived from crushed leaves collected from wild *A. thaliana* plants from the NG2 population was inoculated onto liquid or solid R2A medium to determine what culturable bacteria were present, or onto 2-d or 6-d old axenic *A. thaliana* NG2 (corresponding to seeds with emerged radicles or cotyledons, respectively) to determine which were able to colonize leaves (**Fig. 4A**). Inoculations on media showed that viable cells were present in at least 47% or 6% of drop inoculations (10^-4^ and 10^-5^ dilution, respectively, **Table S5**), showing that the dilution level was appropriate. These included diverse plant-associated taxa spanning the classes Actinobacteria, Alpha-, Beta-, and Gammaproteobacteria, Sphingobacteria, Flavobacteria, and Bacilli (**Fig. S14**). After adding the same inoculum onto germinating seedlings, after two weeks we detected leaf colonization with PCR in only about half (24% and 3% in 10^-4^ and 10^-5^ dilution, respectively, significantly less than in medium, ꭔ^2^: p<0.01 **Table S5**) of inoculations. Thus, many viable cells did not reach detectable levels on leaves. Colonizers in positive plants were mostly *Pseudomonas* sp. and *Xanthomonas* sp., but two were also positive for a *Rhizobiaceae* and one for a Burkholderiales. The number of plants with *Pseudomonas* sp.*, Xanthomonas* sp. or *Rhizobiaceae* was equal to or more than the expected amount based on the number of colonies of those taxa recovered on media (**Figure 4B and Table S6**). This indicates, remarkably, that nearly every viable (culturable) cell of these taxa that was inoculated onto germinating seedlings successfully colonized the plant. Thus, nearly every culturable cell of these taxa that was inoculated onto germinating seedlings reached detectable levels in leaves two weeks later.
